# Lactose blocks intercellular spreading of Galectin-1 from cancer cells to T-cells and activates tumor immunological control

**DOI:** 10.1101/2023.12.19.572484

**Authors:** Yu Hong, Xiaofang Si, Wenjing Liu, Xueying Mai, Yu Zhang

## Abstract

Understanding the mechanisms by which the immune system surveils cancer is the key to developing better tumor immunotherapy strategies. By CRISPR/Cas9 screenings, we identified that inactivation of beta-1,4-galactosyltransferase-1 (B4GALT1), a key enzyme in glycoconjugate biosynthesis, leads to enhanced T-cell receptor (TCR) activation and functions of CD8^+^ T-cells. Via proximity-dependent-intercellular-protein-spreading (PDICPS), cancer cells transfer surface-bound galectin-1 (Gal-1) proteins, which recognize and bind galactosylated membrane proteins, to CD8^+^ T-cells, thereby suppressing T-cell-mediated cytolysis. B4GALT1-deficiency leads to reduced cell-surface galactosylation and Gal-1 binding of CD8^+^ T-cells. Proteomic analysis revealed reduced binding of Gal-1 with TCR and its coreceptor CD8 on B4GALT1-deficient CD8^+^ T-cells, leading to enhanced TCR-CD8 colocalization and T-cell activation. Lactose, a structure-mimicking competitive inhibitor of N-glycan galactosylation, enhances the functions of CD8^+^ T-cells and tumor immunosurveillance. Results from various preclinical tumor models demonstrate that lactose and its derivatives are a new class of immune checkpoint inhibitors for tumor immunotherapy.

## Introduction

Cancer is the leading cause of death worldwide^1^. In 2020, there were 9.96 million cancer-related deaths and 19.29 million new cancer cases globally. Due to the rapid ageing of the global population, environmental changes, the development of new diagnostic techniques, and other factors, it has been predicted that the annual number of new cancer cases will increase to 30 million in 2040. To date, few types of cancers have been shown to be curable; furthermore, how to efficiently treat most cancers is still a major scientific and economic challenge.

Tumor immunotherapies have been rapidly developed recently, and they are one of the most effective strategies for curing cancers^2–4^. It remains unclear why tumor immunotherapies only show significant efficacy for some cancer types and some patients^2^. Understanding the mechanisms by which the immune system surveils cancer is the key to developing better therapeutic strategies and identifying patients who will respond well to immunotherapy. On the other hand, one hidden obstacle for the real-world implication of tumor immunotherapies, including blocking antibodies for immune checkpoints and chimeric antigen receptor (CAR) T-cell therapy, is the high cost of treatment, which is often underestimated^5^. For example, even though the annual cost of anti-PD-1 antibody therapy has recently decreased significantly, it is still not affordable for most patients. It is necessary to develop more economic strategies, such as small molecule inhibitors for immune checkpoints, in order to cure cancer patients, thereby potentially saving millions of lives, especially in developing countries.

Cytotoxic CD8^+^ T-cells play a central role in tumor immunotherapy. The presence and activity of CD8^+^ T-cells in tumors are efficient biomarkers to predict survival and therapeutic efficacy for patients receiving immunotherapy^6,7^. Immune checkpoint inhibitors (e.g., anti-PD-1, anti-PD-L1, and anti-CTLA4 antibodies) targeting the reactivation of tumor-infiltrated cytotoxic CD8^+^ T-cells have shown amazing clinical benefits in treating various human tumors^8–12^.

CAR-T and T-cell receptor-engineered T (TCR-T) cells, which are exogenous cytotoxic CD8^+^ T-cells that are genetically engineered to directly target cancer cells, have demonstrated promising clinical efficacy against some human tumors^13,14^. Fully elucidating the mechanisms regulating the development, differentiation, and functions of cytotoxic CD8^+^ T-cells would be helpful for the development of better tumor immunotherapies.

The expression of PDCD1 (PD-1) can be induced in a wide variety of immune cell types^15–17^. For example, TCR activation induces the expression of PD-1 on the T-cell surface. The interaction between the PD-1 receptor and its ligand, PD-L1, reduces TCR signals to suppress the immune system. When tumors evade the immune response, cancer cells may upregulate their surface PD-L1 expression to prevent endogenous cytotoxic CD8^+^ T-cell attack. Antibodies blocking the interaction between PD-1 and PD-L1 have been proven to be effective in human tumor immunotherapy and show high efficacy in many tumor types^9,10^. Understanding how immune cells regulate their PD-1 expression is one of the most important aspects of tumor immunology research. Although a few regulators of PD-1 expression in cytotoxic CD8^+^ T-cells have been identified recently^18–22^, unbiased systematic screening is still lacking.

Here, by combining ex vivo primary memory CD8^+^ T-cell culture and an in vivo syngeneic mouse tumor model with genome-wide and custom CRISPR/Cas9 screenings, we systematically identified genes and pathways that regulate PD-1 expression and TCR activation in CD8^+^ T-cells. Among them, inactivation of a key enzyme in glycoconjugate biosynthesis, i.e., B4GALT1, enhances TCR activation and the functions of CD8^+^ T-cells both in vitro and in vivo. Via proximity-dependent intercellular protein spreading (PDICPS), cancer cells transfer surface-attached Galectin-1 to CD8^+^ T-cells to suppress T-cell functions, while this checkpoint is impaired in B4GALT1-deficient CD8^+^ T-cells. Proteomic analysis of membrane proteins of CD8^+^ T-cells demonstrated reduced binding of Gal-1 with TCR and its coreceptor CD8 in the absence of B4GALT1, thus leading to enhanced TCR-CD8 colocalization and T-cell activation. Importantly, targeting this novel immune checkpoint by lactose, a natural compound mimicking N-galactosylation, depletes cell surface galectins on both immune and cancer cells, which subsequently leads to the activation of cytotoxic CD8^+^ T-cells within the tumor microenvironment and efficient suppression of tumor growth in syngeneic mice. Lactose is a new class of immune checkpoint inhibitors for tumor immunotherapy.

## Results

### Ex vivo and in vivo CRISPR/Cas9 screenings identify genes and pathways that regulate PD-1 expression, T-cell activation, and functions of CD8^+^ T-cells

We set up an ex vivo genome-wide CRISPR/Cas9 screening system to identify genes and pathways that regulate PD-1 expression in mouse primary CD8^+^ T-cells (Fig. 1a). In brief, splenic CD8^+^ T-cells from Cas9-EGFP/OT-1 mice were infected with a retroviral whole-genome guide RNA (gRNA) library following ovalbumin (Ova) peptide stimulation. After puromycin selection, the memory CD8^+^ T-cells were restimulated by coculturing with B16F10-OVA cells. The highest and lowest 5% PD-1^+^ cells were isolated by fluorescence-activated cell sorting (FACS) and defined as PD-1^high^ and PD-1^low^ populations. The distributions of individual gRNAs in the whole-genome library in those subpopulations, as well as in input cells, were revealed by next-generation sequencing. As shown in Fig. 1b, Pdcd1 and previously identified PD-1 regulators, such as Satb1^18^ and Fut8^20^, were successfully identified as positive controls. Most of the top candidates could be verified individually by FACS and RT‒qPCR assays (Fig. S1a). Gene set enrichment analysis (GSEA) identified several KEGG (Kyoto Encyclopedia of Genes and Genomes) pathways significantly involved in the regulation of PD-1 expression in CD8^+^ T-cells (Fig. 1d). As expected, genes that are known to be involved in the TCR activation pathway such as Cd3d, Zap70, and Lat were among the top ones identified. Interestingly, genes involved in the protein export pathway (such as Srp14, Srp68, Sec16A) and in aminoacyl tRNA biosynthesis (such as Mars, Hars2, Eprs) were significantly increased in PD-1^low^ populations. In addition, we also identified and verified genes required for N-glycan biosynthesis, including B4galt1, Mgat2, and Dpm3, which could negatively regulate the expression of PD-1 (Fig. S1b-d).

**Figure 1.**
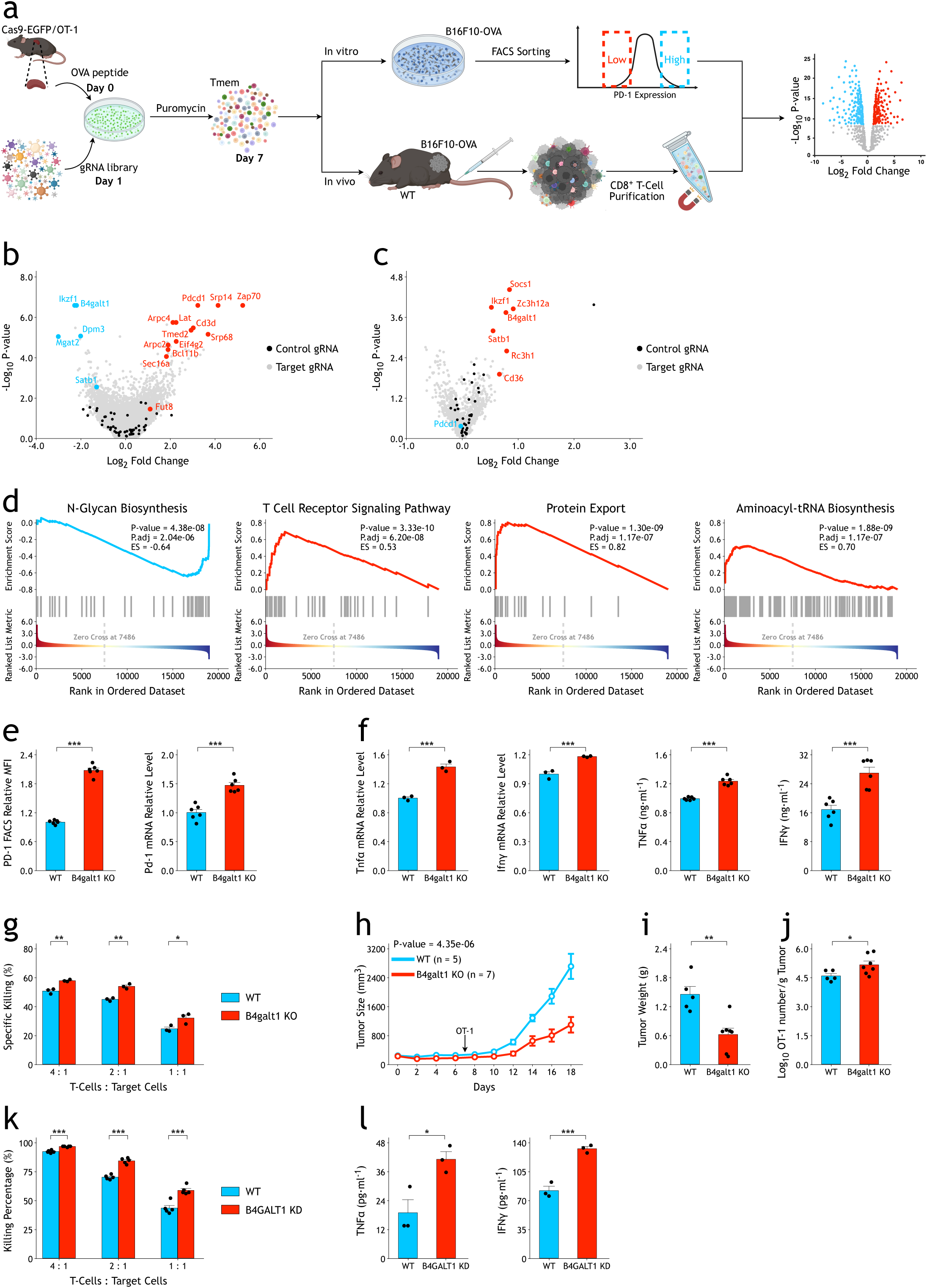
Ex vivo and in vivo CRISPR/Cas9 screenings identify genes and pathways that regulate PD-1 expression and functions of CD8^+^ T-cells. a. Schematic view of ex vivo and in vivo CRISPR/Cas9 screenings in mouse primary CD8^+^ T-cells.
b. Volcano plot showing the results of ex vivo CRISPR/Cas9 genome-wide screenings. The p values were calculated by the α-robust rank aggregation (α-RRA) algorithm in MAGeCK.
c. Volcano plot showing the results of in vivo CRISPR/Cas9 screenings with a small custom library. The p values were calculated by the α-RRA algorithm in MAGeCK.
d. GSEA of significantly enriched KEGG pathways in genome-wide screenings. The enrichment score (ES) and statistical significance were calculated by the clusterProfiler (version 3.12.0) R package.
e. CRISPR/Cas9 knockout of B4GALT1 (sgB4galt1) in CD8^+^ T-cells increases PD-1 expression. The mean fluorescence intensities (MFIs) of PD-1 were measured by FACS. The relative mRNA expression levels of PD-1 were measured by quantitative RT‒qPCR. The p values were calculated by a two-tailed Student’s t test.
f. CRISPR/Cas9 knockout of B4GALT1 in CD8^+^ T-cells increases the expression of TNFα and IFNγ after coculture with B16F10-OVA cells. The relative mRNA expression levels of Tnfα and Ifnγ were measured by quantitative RT‒qPCR. The secreted TNFα and IFNγ in the medium were measured by ELISA. The p values were calculated by a two-tailed Student’s t test.
g. CRISPR/Cas9 knockout of B4GALT1 in OT-1 CD8^+^ T-cells increases in vitro specific killing activities on B16F10-OVA cells. The p values were calculated by a two-tailed Student’s t test.
h. CRISPR/Cas9 knockout of B4GALT1 in OT-1 T-cells enhances growth control of B16F10-OVA tumors in vivo. The p value was calculated by two-way ANOVA.
i. Compared with control wild-type OT-1 T-cells, B16F10-OVA tumors were significantly smaller when B4GALT1 knockout OT-1 T-cells were transplanted. The p value was calculated by a two-tailed Student’s t test.
j. CRISPR/Cas9 knockout of B4GALT1 increases the number of OT-1 T-cells infiltrated into B16F10-OVA tumors. The p value was calculated by a two-tailed Student’s t test.
k. Knockdown of B4GALT1 in human NY-ESO-1 TCR T-cells by shRNA increases in vitro specific killing activities on A375 cells. The p values were calculated by a two-tailed Student’s t test.
l. Knockdown of B4GALT1 in human NY-ESO-1 TCR T-cells increases the expression of TNFα and IFNγ after coculture with A375 cells. The secreted TNFα and IFNγ in the medium were measured by ELISA. The p values were calculated by a two-tailed Student’s t test. Data are shown as the mean ± SEM. *P < 0.05; **P < 0.01; ***P < 0.001.

To verify these candidate genes in a high-throughput manner and test their potential functions in the tumor microenvironment (Fig. 1a), we synthesized a custom gRNA library containing 4,617 gRNAs targeting the top candidate genes obtained from whole-genome screenings and 105 intergenic control gRNAs. The gRNA library-infected Cas9-EGFP/OT-1 memory CD8^+^ T-cells were restimulated by coculture in vitro or transplanted into wild-type C57BL/6J mice inoculated subcutaneously with B16F10-OVA tumors to screen for genes regulating CD8^+^ T-cell functions in vivo. After 7 days, CD8^+^ T-cells were collected from the tumors for gRNA sequencing. In our screenings, we successfully identified several positive control genes^23–26^ such as Socs1, Regnase-1 (Zc3h12a), Rc3h1, and Cd36, indicating our in-vivo screening system worked robustly (Fig. 1c). In consistent with these genes, inactivating B4galt1, which encodes beta-1,4-galactosyltransferase 1, showed significant phenotypes in both ex vivo and in vivo screenings.

### Ablation of B4GALT1 in CD8^+^ T-cells activates TCR signaling and enhances T-cell-mediated tumor immunotherapy

B4GALT1 is one of the seven beta-1,4-galactosyltransferases that transfer galactose in a beta 1-4 linkage to similar acceptor sugars, including N-acetylglucosamine (GlcNAc), glucose (Glc), and xylose (Xyl). More specifically, B4GALT1 uses UDP-galactose and N-acetylglucosamine for the production of galactose beta-1,4-N-acetylglucosamine^27^. In addition to glycoconjugate biosynthesis, B4GALT1 can also form a heterodimer with α-lactalbumin (LALBA) as lactose synthetase in lactating tissues. Although B4GALT1 is expressed ubiquitously, its roles in regulating the interaction and adhesion of immune cells have been demonstrated^28–30^.

To determine the functions of B4GALT1 in CD8^+^ T-cells, we infected Cas9-EGFP/OT-1 CD8^+^ T-cells with different gRNAs targeting B4GALT1, and it showed a significantly increased surface PD-1 expression and PD-1 mRNA levels, comparing with control T-cells (Fig. 1e and Fig. S1d). Such phenotypes could be rescued by overexpression of either the short or long isoform of mouse B4galt1 cDNA (Fig. S2a), suggesting that the biosynthetic function, but not ligand-induced signal transduction, of B4GALT1^27^ is responsible for the suppression of TCR activation. In addition, compared with control cells, B4GALT1 knockout OT-1 T-cells showed significantly increased targeted cell killing activity in vitro, as well as enhanced expression of T-cell activation and cytotoxic markers such as IFNγ and TNFα (Fig. 1f-g). Whole-genome RNA sequencing analysis confirmed the enhanced TCR activation in B4GALT1 knockout CD8^+^ T-cells (Fig. S2b-d). GSEA of DEGs (differentially expressed genes) between control and B4galt1 gRNA-infected CD8^+^ T-cells revealed that the TCR signaling pathway was at the top of the significantly altered pathways (Fig. S2d).

When transplanted into wild-type mice with B16F10-OVA cells inoculated subcutaneously, B4galt1 gRNA-infected OT-1 T-cells showed significantly higher tumor killing activity than control gRNA-infected cells (Fig. 1h-i). Analysis of tumor-infiltrated lymphocytes (TILs) demonstrated more infiltrated OT-1 T-cells in tumors when B4galt1 gRNA was infected (Fig. 1j). Mechanistically, B16F10-OVA tumors had similar numbers of infiltrated B4galt1 gRNA-infected OT-1 T-cells as control gRNA-infected cells 24 hours after intravenous injection (Fig. S3a), suggesting that B4GALT1 has no significant effect on tumor infiltration of OT-1 T-cells. On the other hand, CFSE (carboxyfluorescein succinimidyl ester) analysis at a later time point (6 days after infusion) showed increased OT-1 T-cell proliferation within tumors when B4GALT1 was inactivated (Fig. S3b). Finally, inactivation of B4GALT1 also significantly enhanced the target cell killing activity of human NY-ESO-1 TCR T-cells in vitro (Fig. 1k-l). These results suggest that inhibition of N-galactosylation could enhance the functions of CD8^+^ T-cells and that B4GALT1 could be a potential target to modulate the activity of TCR T-cells.

### Cancer cells transfer cell surface Galectin-1 to CD8^+^ T-cells to inhibit T-cell activation, which is impaired in B4GALT1-deficient CD8^+^ T-cells

To dissect the molecular mechanism by which B4GALT1 regulates T-cell activation, we first used FACS staining with biotin-labeled *Erythrina cristagalli* lectin (ECL) and succinyl-wheat germ agglutinin (sWGA) to profile the surface expression of terminal βGal and βGlcNAc, respectively. B4GALT1 knockout OT-1 T-cells showed slightly but significantly decreased ECL staining and significantly increased sWGA staining, comparing with WT control OT-1 T cells (Fig. S4). Surprisingly, FACS staining with biotin-labeled recombinant Galectin-1 showed a more dramatic difference between control and B4GALT1 knockout OT-1 T-cells (Fig. 2a, d and Fig. S4c).

**Figure 2.**
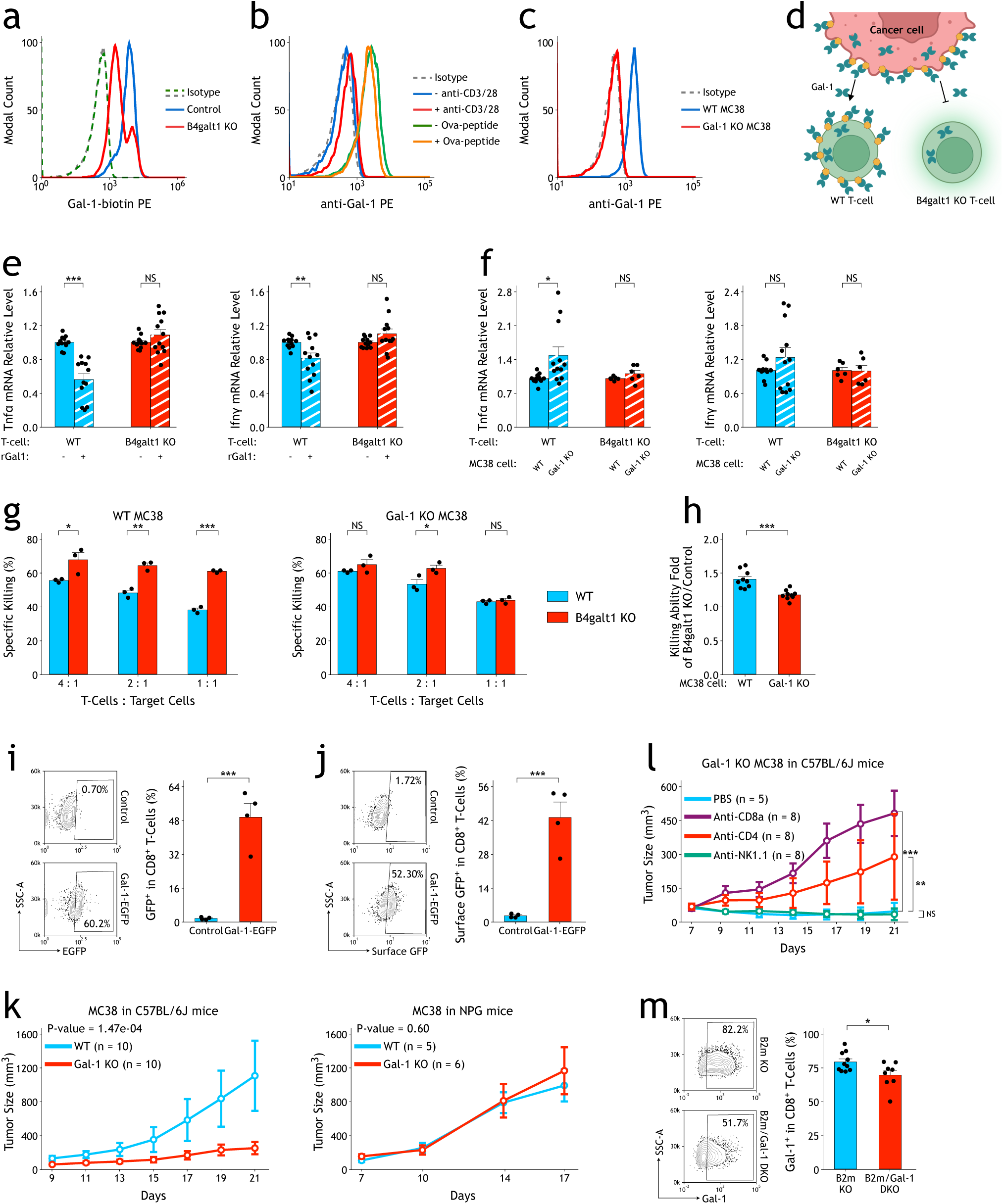
Cancer cells transfer cell surface Gal-1 to CD8^+^ T-cells to inhibit TCR activation, which is impaired in B4GALT1-deficient CD8^+^ T-cells. a. B4GALT1 knockout and control OT-1 cells were incubated with biotin-recombinant Gal-1 (rGal-1) and then stained with streptavidin-PE. Blue and red curves indicate control and B4GALT1 knockout OT-1 cells, respectively; gray and green dotted curves indicate control and B4GALT1 knockout OT-1 cells, respectively, stained with streptavidin-PE only.
b. OT-1 cells with (red) or without (blue) TCR activation by anti-CD3/28 antibodies and OT-1 cells cocultured with MC38 cells with (orange) or without (green) prior Ova-peptide pulsing were stained with anti-Gal-1 antibody. The gray dotted curve indicates OT-1 cells stained with isotype control antibody.
c. OT-1 cells show no significantly increased surface Gal-1 after coculture with Gal-1 knockout MC38 cells (red curve) compared with OT-1 cells without coculture (gray dotted curve). The blue curve indicates OT-1 cells cocultured with control wild-type MC38 cells (blue curve).
d. Schematic of surface Gal-1 transfer from cancer cells to T-cells.
e. rGal-1 treatment (2.5 µg/ml) decreases the mRNA expression levels of Tnfα and Ifnγ after TCR activation by anti-CD3/CD28 antibodies in control wild-type OT-1 cells (1×10^5^ cells/ml) but shows no significant effect on B4GALT1 knockout OT-1 cells. The p value was calculated by a two-tailed Student’s t test.
f. Compared with wild-type MC38 cells, coculture with Gal-1 knockout MC38 cells increased the mRNA expression levels of Tnfα and Ifnγ in control wild-type OT-1 cells after activation by anti-CD3/CD28 antibodies. Such an effect is absent for B4GALT1 knockout OT-1 cells. The p value was calculated by a two-tailed Student’s t test.
g. B4GALT1 KO and control wild-type OT-1 cells were used to kill wild-type control (left panel) and Gal-1 knockout (right panel) MC38 cells pulsed with Ova-peptide. The p value was calculated by a two-tailed Student’s t test.
h. The fold change in the killing ability of B4GALT1 knockout OT-1 cells versus wild-type control OT-1 cells was calculated when wild-type or Gal-1 knockout MC38 cells were used. Data for a 2:1 ratio (T-cells: MC38 cells) were used. The p value was calculated by a two-tailed Student’s t test.
i. MC38 cells infected with exogenous expression of Gal-1-EGFP fusion or control EGFPs were inoculated subcutaneously into wild-type C57BL/6J mice. After 3 weeks, tumor-infiltrated CD8^+^ T-cells showed a GFP signal only in Gal-1-EGFP tumors. The p value was calculated by a two-tailed Student’s t test.
j. Tumor-infiltrated CD8^+^ T-cells showed a surface GFP signal by anti-GFP antibody staining only for Gal-1-EGFP tumors. The p value was calculated by a two-tailed Student’s t test.
k. Gal-1 knockout (sgGalectin-1) MC38 and wild-type control MC38 cells were inoculated subcutaneously into wild-type C57BL/6J (left panel) and NPG (NOD-Prkdc^scid^IL2rg^null^) mice (right panel). The p value was calculated by two-way ANOVA.
l. Gal-1 knockout MC38 cells were inoculated subcutaneously into wild-type mice depleted of CD8^+^ T-cells, CD4^+^ T-cells, and NK cells. The p value was calculated by two-way ANOVA.
m. B2m knockout and Gal-1/B2m double knockout MC38 cells were inoculated subcutaneously into wild-type C57BL/6J mice. After 3 weeks, tumor-infiltrated CD8^+^ T-cells were stained with anti-Gal-1 antibody. The p value was calculated by a two-tailed Student’s t test. Data are shown as the mean ± SEM. *P < 0.05; **P < 0.01; ***P < 0.001. NS, not significant.

RNA sequencing analysis revealed that galectin-1, galectin-3, and galectin-9 (Gal-1, Gal-3, Gal-9) were the most highly expressed galectins in OT-1 T-cells and MC38 cells (Fig. S5a). Intracellular FACS staining confirmed that all galectins were well expressed in both OT-1 T-cells and MC38 cells (Fig. S5b). However, Gal-1, -3, and -9 were absent on the OT-1 T-cell surface, while Gal-1 was more significantly present on the surface of MC38 cells than Gal-3 and Gal-9 (Fig. S5b-c).

After coculture with MC38 cells either with or without Ova-peptide pulsing, Gal-1 staining on the OT-1 T-cell surface increased dramatically, suggesting that interaction between the TCR receptor and MHC-1/Ova complex or TCR activation was not necessary (Fig. 2b). On the other hand, activation by anti-CD3/CD28 antibodies alone could not increase the cell surface Gal-1 staining of OT-1 T-cells in the absence of MC38 cells (Fig. 2b). In accordance with these findings, compared with wild-type controls, B4GALT1 knockout OT-1 T-cells did not show significantly enhanced T-cell activation by anti-CD3/28 antibodies in the absence of MC38 cells (Fig. S6). Taken together, these results suggest that increased Gal-1 staining on the OT-1 T-cell surface might come from cancer cells in coculture. Indeed, Gal-1 staining on the OT-1 T-cell surface only increased dramatically after coculture with wild-type but not Gal-1 knockout MC38 cells (Fig. 2c, Fig. S7 and Fig. S8). Incubation with conditioned medium from neither wild-type nor Gal-1 knockout MC38 cells increased surface Gal-1 staining on OT-1 T-cells (Fig. S8). Moreover, when OT-1 T-cells and MC38 cells were physically separated by a membrane in a Boyden chamber or a thin layer of low-melting agarose, Gal-1 protein could not be efficiently transferred from MC38 cells to OT-1 T-cells, suggesting that cell‒cell interaction and/or close proximity is necessary (Fig. S8). Different from known mechanisms for intercellular transfer of cell-surface proteins^31,32^, such as trogocytosis, exosomal protein transport, and shuttle of proteins through nanotubes, we named this process as proximity-dependent intercellular protein spreading (PDICPS).

Adding exogenous recombinant Gal-1 protein to wild-type OT-1 T-cells could significantly inhibit T-cell activation by anti-CD3/28 antibodies, while this effect was absent for B4GALT1-deficient OT-1 T-cells (Fig. 2e). OT-1 T-cells cocultured with Gal-1 knockout MC38 cells without Ova pulsing also showed enhanced T-cell activation by anti-CD3/28 stimulation compared with cells cocultured with wild-type MC38 cells (Fig. 2f). In contrast, B4GALT1 knockout OT-1 T-cells showed similar T-cell activation in such conditions. Compared with Gal-1 knockout MC38 cells, wild-type MC38 cells were more sensitive to B4GALT1 knockout OT-1 T-cell-mediated specific killing than wild-type OT-1 cells (Fig. 2g-h). These results demonstrated the essential roles of cancer cell-surface transferred Gal-1 in regulating T-cell activation.

We hypothesized that in the tumor microenvironment, cancer cells may transfer their cell surface Gal-1 to proximal tumor-specific CD8^+^ T-cells to suppress immune control. We overexpressed Gal-1-GFP fusion protein in MC38 cells to trace Gal-1 transfer in vivo. CD8^+^ T-cells infiltrated in Gal-1-GFP MC38 tumors showed strong total GFP and surface GFP signals (Fig. 2i-j). Moreover, compared with control wild-type cells, the growth of subcutaneously inoculated Gal-1 knockout MC38 cells was greatly reduced in wild-type C57BL/6J but not immunodeficient NPG (NOD-Prkdc^scid^IL2rg^null^) mice (Fig. 2k). When subcutaneously inoculated into wild-type mice depleted of different immune cell types by specific antibodies (Fig. S9), the growth of Gal-1 knockout MC38 tumors could be significantly rescued by CD8^+^ and CD4^+^ T-cell depletion but not NK-cell depletion (Fig. 2l). B2m knockout also rescued the growth of Gal-1 knockout MC38 tumors in wild-type mice (Fig. S10). We examined endogenous surface Gal-1 staining on tumor-infiltrated CD8^+^ T-cells in vivo. As shown in Fig. S12, surface Gal-1 expression could hardly be detected on CD8^+^ T-cells from the peripheral blood and spleen of wild-type mice. In contrast, tumor-infiltrated CD8^+^ T-cells, which were mostly PD-1-positive and represented exhausted CD8^+^ T-cells, were positive for surface Gal-1 staining (Fig. S11). Compared with B2m-deficient MC38 tumors, CD8^+^ T-cell surface Gal-1 signals were significantly reduced in Gal-1/B2m double knockout MC38 tumors, suggesting that MC38 cells are major donors for Gal-1 on the CD8^+^ T-cell surface in vivo (Fig. 2m). To summarize, our results demonstrated a novel immune checkpoint mechanism in which cancer cells could transfer their surface Gal-1 to the proximal CD8^+^ T-cell surface to suppress T-cell activation both in vitro and in vivo.

### B4GALT1 regulates TCR-CD8 colocalization and TCR activation via surface Gal-1

To identify the substrates of B4GALT1 on the CD8^+^ T-cell surface, we used a recombinant Gal-1 affinity column to purify Gal-1 binding proteins from membrane protein extracts of OT-1 CD8^+^ T-cells (Fig. S12a). Analysis of proteins that were significantly different between wild-type and B4GALT1 knockout OT-1 T-cells revealed that both TCRα/β (OT-1) and CD8α/β were among the top list (Fig. S12b). KEGG analysis showed a significant enrichment of the TCR signaling pathway (Fig. S12c). We verified the reduced Gal-1 binding with CD8β in B4GALT1 knockout T-cells by western blotting (Fig. S12d). Interestingly, the migration of CD8β in SDS‒PAGE was different between wild-type control and B4GALT1 knockout T-cell extracts, suggesting that CD8β is a direct substrate of B4GALT1 (Fig. S12d). Indeed, treating Gal-1 pull-down products with PNGase F to remove all glycosylation on proteins omitted the migration difference of CD8β (Fig. S12e). It has been suggested that regulation of TCR-CD8 colocalization by galectin binding might be one major mechanism of tumor immunological control^33,34^. We measured the interaction between TCR and CD8 by FRET assay^34^ (Fig. S12f). As shown in Fig. S12g, by adding recombinant Gal-1 to wild-type OT-1 cells, the TCR-CD8 FRET signal was significantly reduced, and this phenotype could be reversed by lactose treatment. Such an effect of rGal-1 on TCR/CD8 FRET was less significant for B4GALT1 knockout OT-1 T-cells.

Taken together, our results support a model in which cancer cells transfer their surface Gal-1 to the T-cell surface, where Gal-1 proteins inhibit TCR activation by interfering with the interaction between TCR and CD8 T-cells. Accordingly, inactivation of B4GALT1 could not affect CAR-T-mediated target cell killing, which is CD8-independent (Fig. S13).

### Targeted removal of cell surface Gal-1 by lactose, a structure-mimicking competitive inhibitor of N-galactosylation, activates CD8^+^ T-cells in vitro and in vivo

In the milk-secreting mammary gland, B4GALT1 interacts with LALBA to form lactose synthase, which transfers the galactose moiety to glucose. Since the galactose moiety of lactose can compete and interfere with the functions of beta-galactosides in glycoproteins and glycolipids ^33,35–38^, we hypothesized that lactose and its derivatives might be a new class of immune checkpoint inhibitors to deplete Gal-1 on the CD8^+^ T-cell surface. Indeed, we found that lactose enhanced the specific killing of Ova-pulsed MC38 cells (Fig. 3a) by OT-1 T-cells. In this in vitro killing system, lactose treatment removed Gal-1 from the OT-1 cell surface and significantly increased the expression of effector cytokines such as Ifnγ and Tnfα in OT-1 T-cells, suggesting enhanced T-cell activation (Fig. 3b). However, when stimulated by anti-CD3/CD28 antibodies in the absence of exogenous Gal-1, lactose treatment could not significantly affect Ifnγ and Tnfα expression by OT-1 T-cells (Fig. S15). Lactose treatment relieved the inhibitory effect of recombinant Gal-1 and MC-38-transferred Gal-1 on OT-1 cells (Fig. 3c, Fig. S14 and Fig. S16). This treatment also depleted surface Gal-1 and significantly enhanced Ifnγ and Tnfα expression in exhausted CD8^+^PD-1^+^ T-cells isolated from MC38 tumors (Fig. S17).

**Figure 3.**
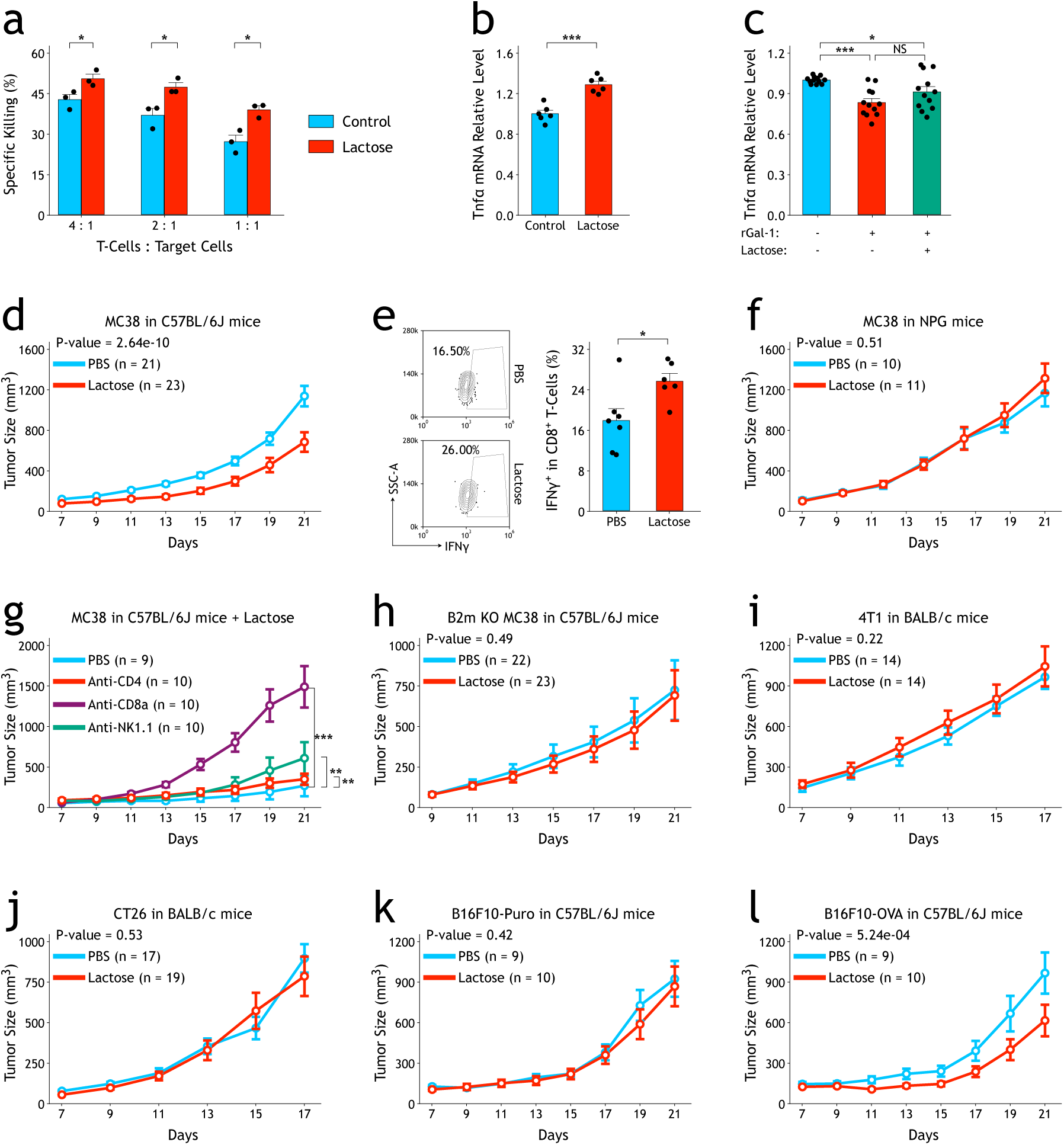
Lactose, a competitive inhibitor of galectins, enhances tumor immunosurveillance in vitro and vivo. a. Lactose treatment increases in vitro specific killing activities of OT-1 CD8^+^ T-cells on OVA-peptide pulsed MC38 cells. The p values were calculated by a two-tailed Student’s t test.
b. Lactose treatment increases the expression of Tnfα in OT-1 CD8^+^ T-cells after coculture with OVA peptide-pulsed MC38 cells. The relative mRNA expression levels of Tnfα were measured by quantitative RT‒qPCR. The p values were calculated by a two-tailed Student’s t test.
c. rGal-1 treatment reduces the expression of Tnfα in OT-1 CD8^+^ T-cells after anti-CD3/CD28 antibody stimulation. Such an inhibitory effect of rGal-1 could be reversed by lactose treatment. The relative mRNA expression levels of Tnfα were measured by quantitative RT‒qPCR. The p values were calculated by a two-tailed Student’s t test.
d. The effect of intravenous injection of lactose on growth of subcutaneous MC38 tumors in syngeneic C57BL/6J wild-type mice. The p value was calculated by two-way ANOVA.
e. Intravenous lactose injection increases percentages of MC38 tumor-infiltrated IFNγ-positive CD8^+^ T-cells. The p value was calculated by a two-tailed Student’s t test.
f. The effect of intravenous lactose injection on the growth of subcutaneous MC38 tumors in immunodeficient NPG mice. The p value was calculated by two-way ANOVA.
g. The effect of intravenous lactose injection on the growth of MC38 tumors in syngeneic C57BL/6J wild-type mice depleted of CD8^+^ T-cells, CD4^+^ T-cells, and NK cells. The p values were calculated by two-way ANOVA.
h. The effect of intravenous lactose injection on the growth of B2m knockout MC38 tumors in syngeneic C57BL/6J wild-type mice. The p value was calculated by two-way ANOVA.
i. The effect of intravenous lactose injection on the growth of 4T1 tumors in syngeneic BALB/c wild-type mice. The p value was calculated by two-way ANOVA.
j. The effect of intravenous lactose injection on the growth of CT26 tumors in syngeneic BALB/c wild-type mice. The p value was calculated by two-way ANOVA.
k. The effect of intravenous lactose injection on the growth of B16F10-puro tumors in syngeneic C57BL/6J wild-type mice. The p value was calculated by two-way ANOVA.
l. The effect of intravenous lactose injection on the growth of B16F10-OVA tumors in syngeneic C57BL/6J wild-type mice. The p value was calculated by two-way ANOVA. Data are shown as the mean ± SEM. *P < 0.05; ***P < 0.001.

To test the effect of lactose in vivo, we injected lactose solution intravenously every two days into wild-type mice subcutaneously inoculated with MC38 tumors. In comparison with the PBS injection, the lactose injection significantly reduced the growth of MC38 tumors and increased IFNγ-positive CD8^+^ T-cells in tumors (Fig. 3d-e, Fig. S18), while this effect was absent when immunodeficient NPG mice were used (Fig. 3f). Depleting CD8^+^ T-cells in wild-type mice significantly compromised the effect of lactose injection on MC38 tumors, while CD4^+^ T-cells and NK cells showed less contribution (Fig. 3g). B2m-deficient MC38 tumors did not respond to lactose treatment (Fig. 3h). Moreover, intravenous injection of lactose did not synergistically increase the efficacy of anti-PD-1 or anti-PD-L1 antibody treatment on MC38 tumors in syngeneic mice (Fig. S19).

Intravenous lactose treatment also did not significantly affect the growth of 4T1 and CT26 tumors in wild-type syngeneic mice (Fig. 3i-j), suggesting being immune “cold” tumors^39^. In comparison with control cells, exogenous overexpression of Ova protein in B16F10 cells significantly increased its sensitivity to lactose treatment (Fig. 3k-l) by converting it to a “hot” tumor^40^.

In the absence of functional mammalian transporters, lactose cannot enter cells efficiently (Fig. S20a) and is chemically stable in the circulation system^41^. Therefore, we hypothesized that intravenous administration of lactose inhibits tumor immune checkpoints not mainly through its degradation of monosaccharide products (galactose and glucose). To confirm that the galactose moiety is the main functional group in lactose that binds lectins and galectins, we tested sucrose, N-acetyllactosamine (LacNAc), and lactose-BSA (Fig. S20b). Compared with lactose, LacNAc could compete with the binding of ECL and Gal-1 on the cell surface more efficiently but could also compete with the binding of sWGA (Fig. S20c). In vivo, LacNAc also demonstrated significant inhibition of MC38 tumor growth (Fig. S20d). Lactose in blood is depleted quickly via the kidney^42^. We coupled lactose to BSA efficiently^43^ (Fig. S20b), which may improve its pharmacological dynamics in vivo^44^. All of the findings support our model and indicate that galactose motifs compete with the binding of galectins on the cell surface and subsequently modulate cellular receptor activities.

Finally, we checked the safety profiles of intravenous lactose treatment for wild-type C57BL/6J mice. As shown in Fig. S21, neither short-term (24 hours) (Fig. S21a) nor long-term (24 days) (Fig. S21b-c) treatment with intravenous lactose injection every two days showed significant general toxicity or other obvious side effects.

### Lactose alters the tumor microenvironment and enhances the expansion and functions of tumor-specific CD8^+^ T-cells

Bulk RNA sequencing analysis of lactose- and control-treated subcutaneous MC38 tumors in wild-type mice identified 793 significantly upregulated and 742 significantly downregulated genes (Fig. S22a-b). The HALLMARK pathways, including interferon gamma response, hypoxia, TNFα signaling via NFκB, and Myc targets (V1 and V2), were significantly enriched after lactose treatment (Fig. S22c). Analysis of gene expression signatures (Fig. S22d-e) revealed that lactose-treated tumors had significantly higher expression of the T accumulation positive signature^45^ and T-cell cytotoxicity signature^46^ than PBS control.

To further dissect the potential effect of lactose treatment on the tumor microenvironment, especially tumor-infiltrated T-cells, we performed single-cell RNA sequencing and TCR sequencing for FACS-sorted CD3-positive T-cells from lactose- and control-treated MC38 tumor samples (Fig. S23). T-cells were then annotated by ProjectTILs^47^. In lactose-treated tumors, the percentages of exhausted CD8^+^ T (CD8_Tex) cells increased from 39.8% to 57.2%, while naïve-like CD8^+^ T (CD8_NaiveLike) and CD4^+^ Treg (Treg) cells decreased from 15.5% to 7.4% and from 25.0% to 15.9%, respectively (Fig. S23c). TCR clonality analysis demonstrated that lactose-treated tumors had significantly more expanded larger TCR clones than controls (Fig. S24a-d).

The largest TCR clone, CN1 (CDR3: TRA;TRB=CAIDPYNQGKLIF; CASSPGQGYAEQFF), against a specific neoantigen (ASMTNMELM in the Adpgk gene) of MC38 cells^48^ increased from 2.3% to 34.2% in all CD3^+^, 2.3% to 32.4% in CD8_Tex, 0.03% to 0.8% in CD8_Tpex, and 0.05% to 0.9% in CD8_EffectorMemory after lactose treatment (Fig. S24e-f). Both bulk and single-cell RNA sequencing results demonstrate that lactose treatment enhances the expansion and functions of tumor-specific CD8^+^ T-cells in the tumor microenvironment.

### LALBA expression activates the tumor immune response by de novo synthesis of lactose

We hypothesized that exogenously overexpressing LALBA in cancer cells could lead to de novo lactose biosynthesis within the tumor microenvironment, which subsequently may activate tumor immunological control. As shown in Fig. S25a-c, overexpressing the LALBA protein using a lentiviral expression system in MC38 cells led to B4GALT1-dependent de novo synthesis and secretion of lactose, while D107A and A126K mutations in LALBA^49,50^ resulted in inactivation of lactose synthase activity. After subcutaneous inoculation in wild-type mice, the growth of MC38 tumors with LALBA overexpression was significantly inhibited as compared with that of controls (GFP only) (Fig. 4a). Importantly, in more than 50% of mice, we observed complete regression (CR) of MC38-LALBA tumors. In contrast, LALBA-overexpressing MC38 cells showed tumor growth similar to that of MC38 control cells in immunodeficient NPG mice (Fig. 4b). We then rechallenged MC38-LALBA-CR mice with MC38-control cells. Compared with age-matched naïve wild-type control mice, MC38-LALBA-CR mice were completely resistant to MC38-control tumors, suggesting that tumor-specific immunological memory had been established (Fig. 4c). We also subcutaneously inoculated a mixture of MC38-control with MC38-LALBA cells at a 1:1 or 9:1 ratio in wild-type mice. As shown in Fig. S26, mixing MC38-control with MC38-LALBA cells in the same tumor enhanced tumor growth control (6/22 CR for a 1:1 ratio and 3/22 CR for a 9:1 ratio). All these results suggest that exogenous expression of LALBA in tumors, even partially, could enhance tumor immunological control and memory.

**Figure 4.**
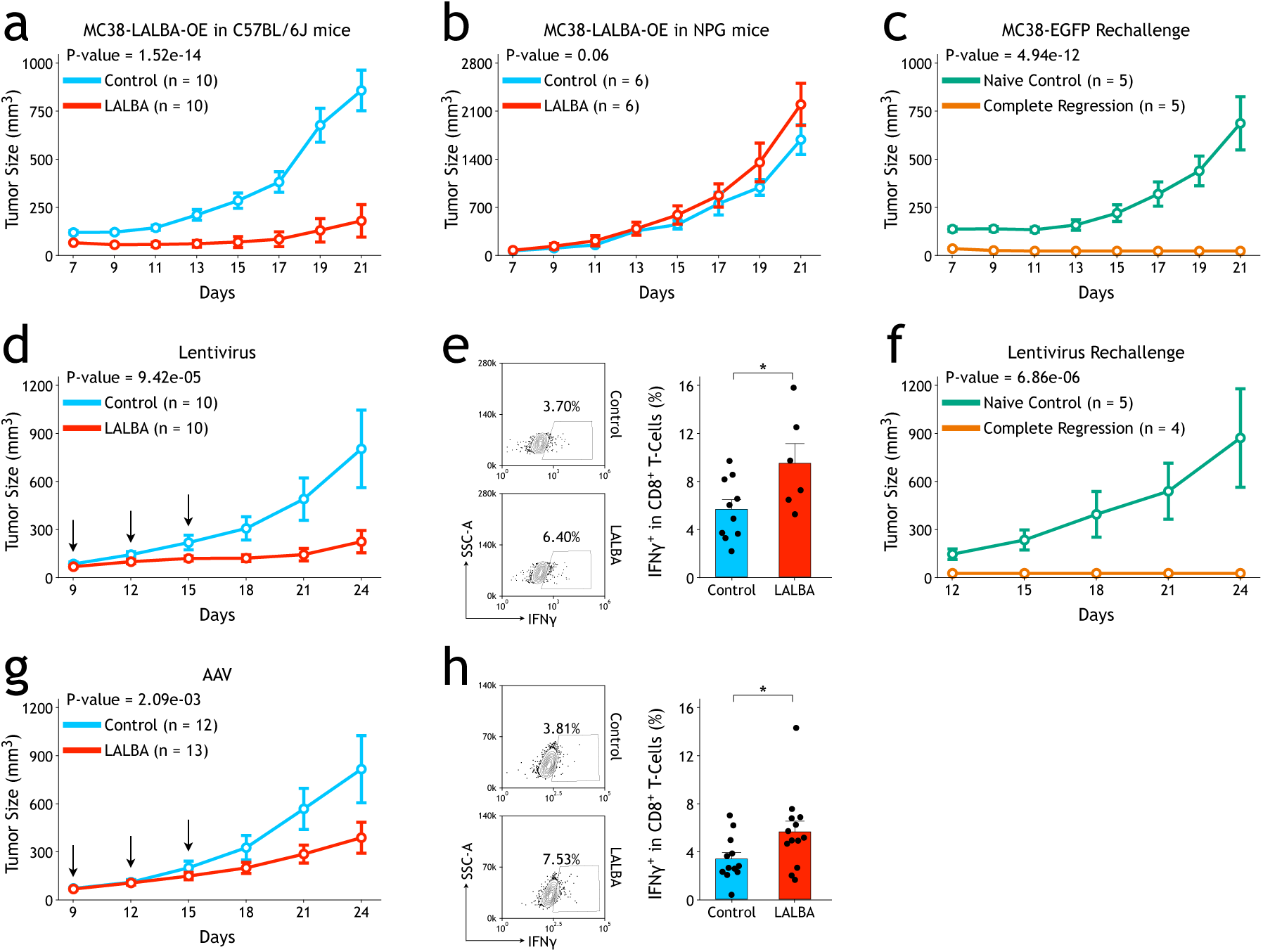
Intratumoral overexpression of α-lactalbumin (LALBA) leads to de novo synthesis of lactose within the tumor microenvironment and activates the tumor immune response. a. Overexpression of LALBA in MC38 cells suppresses tumor growth in wild-type mice. The p value was calculated by two-way ANOVA.
b. Overexpression of LALBA in MC38 cells does not affect tumor growth in immunodeficient NPG mice. The p value was calculated by two-way ANOVA.
c. Compared with age-matched wild-type (naïve) mice, MC38-GFP tumors do not grow in wild-type CR mice, which have completely eliminated their MC38-LALBA tumors (complete regression, CR). The p value was calculated by two-way ANOVA.
d. Intratumoral injection of LALBA-expressing lentivirus reduces the growth of subcutaneous MC38 tumors in syngeneic wild-type mice. Arrows indicate the time points for lentivirus injection. The p value was calculated by two-way ANOVA.
e. Intratumoral injection of lenti-LALBA increases the percentages of tumor-infiltrated IFNγ-positive CD8^+^ T-cells. The p value was calculated by a two-tailed Student’s t test.
f. MC38 tumors cannot grow in mice in which subcutaneous MC38 tumors were completely eliminated by lenti-LALBA treatment. The p value was calculated by two-way ANOVA.
g. Intratumoral injection of AAV-LALBA reduces the growth of subcutaneous MC38 tumors in syngeneic wild-type mice. Arrows indicate the time points for AAV injection. The p value was calculated by two-way ANOVA.
h. Intratumoral injection of AAV-LALBA increases the percentages of tumor-infiltrated IFNγ-positive CD8^+^ T-cells. The p value was calculated by a two-tailed Student’s t test. Data are shown as the mean ± SEM. *P < 0.05.

To achieve exogenous overexpression of LALBA within tumors, we intratumorally injected a lentivirus encoding control or LALBA overexpression cassette. As shown in Fig. 4d, three intratumoral injections of lenti-LALBA led to significant inhibition of MC38 tumor growth in wild-type mice (CR=4/10), while LALBA mutants (D107A and A126K) showed no significant effect (Fig. S27). FACS analysis revealed increased IFNγ^+^ CD8^+^ T-cell numbers in lenti-LALBA-injected tumors than that in control lentivirus-injected tumors (Fig. 4e). In addition, CR mice after lenti-LALBA injections were completely resistant to rechallenge with MC38 cells (Fig. 4f). We also used adeno-associated virus (AAV) to overexpress LALBA in MC38 tumors and observed a similar effect (Fig. 4g-h).

### Lactose activates human tumor-residential T-cells within the tumor microenvironment and architecture

To test the potential translational implications of lactose treatment for human tumor immunotherapy, we treated NPG mice engrafted with human immune cells and human SGC7901 gastric tumor xenografts by intravenous administration of lactose solution or PBS. Similar to pembrolizumab (Fig. S28), an anti-human PD-1 antibody, lactose treatment significantly reduced the growth of subcutaneously engrafted SGC7901 tumors in humanized NPG mice (Fig. 5a). After lactose treatment, SGC7901 tumors contained higher percentages of infiltrated human IFNγ^+^ CD8^+^ T-cells compared with control samples (Fig. 5b). Importantly, this tumor suppression effect of lactose treatment was absent in NPG mice without engrafted human immune cells (Fig. 5c).

**Figure 5.**
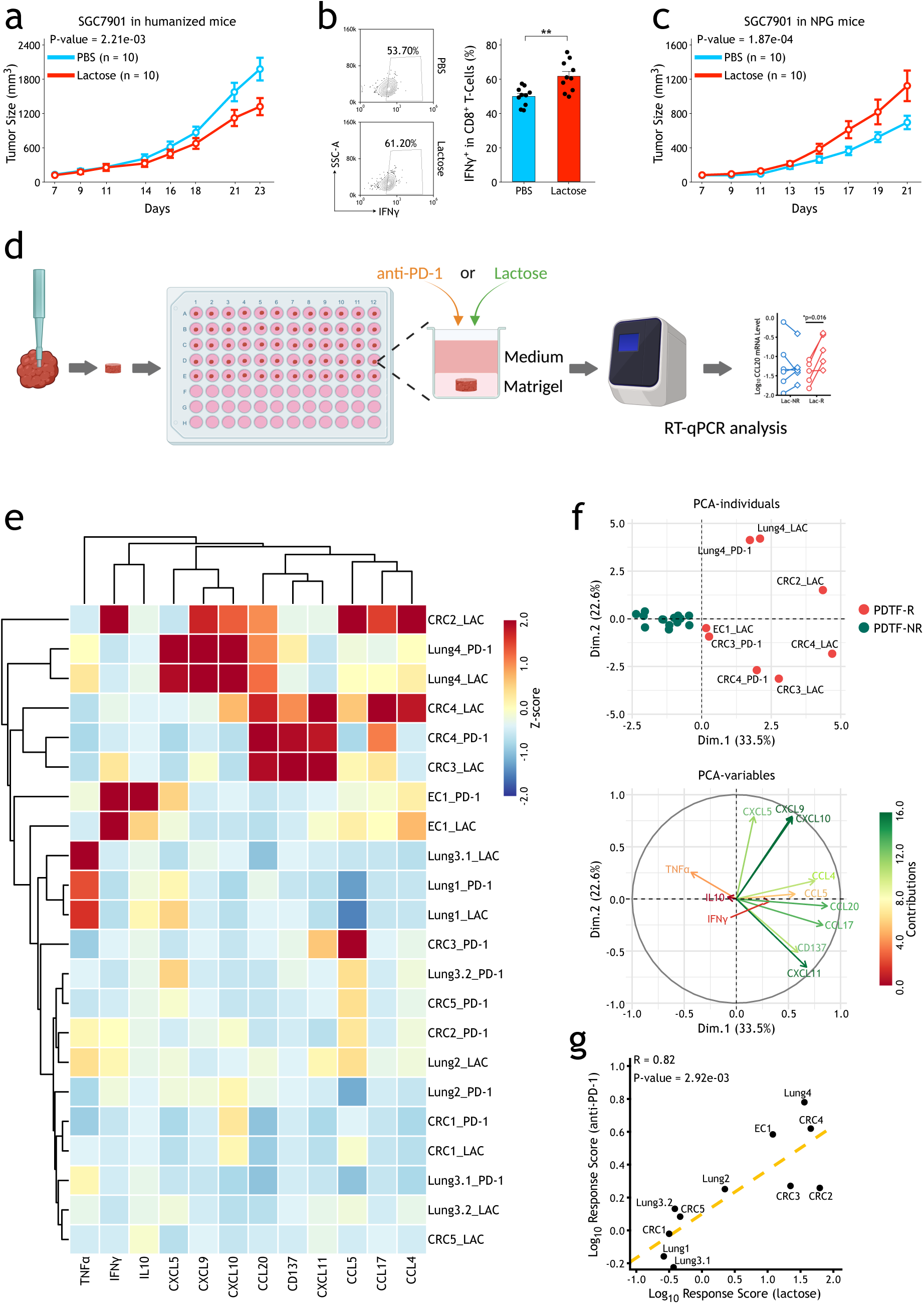
Lactose activates human tumor-residential T-cells within the tumor microenvironment and architecture. a. Effect of intravenous lactose treatment on growth of human SGC7901 tumors in immunodeficient NPG mice reconstituted with human immune system. The p value was calculated by two-way ANOVA.
b. Intravenous lactose treatment increases percentages of SGC7901 tumor-infiltrated IFNγ-positive human CD8^+^ T-cells. The p value was calculated by a two-tailed Student’s t test.
c. Effect of intravenous lactose treatment on the growth of human SGC7901 tumors in immunodeficient NPG mice. The p value was calculated by two-way ANOVA.
d. Schematic view of ex vivo culture of patient-derived tumor fragments.
e. Heatmap displaying normalized z scores between lactose- or anti-PD-1-treated and untreated conditions for 12 parameters.
f. PCA of the data in e, showing separation of samples (top) and parameters (bottom).
g. Correlation between response scores for lactose and those for anti-PD-1 treatment. The correlation value and p value were calculated by Pearson’s product-moment correlation analysis in R. Data are shown as the mean ± SEM. **P < 0.01.

Finally, we used ex vivo culture of patient-derived tumor fragments^51^ to profile the effect of pembrolizumab and lactose treatment on 11 human primary tumor samples from 3 different cancer types (lung cancer, colorectal cancer, and esophageal cancer) (Fig. 5d-e and Fig. S29). As shown in Fig. 5f, the PCA results suggested that 5 out of 11 tumor samples showed a positive response to lactose treatment. Interestingly, the ex vivo response of human primary tumor tissues to pembrolizumab and lactose treatment could be correlated very well (Fig. 5g). These preclinical results suggest that lactose treatment could be beneficial for cancer patients who are predicted to respond to anti-PD-1 antibody therapy.

## Discussion

Therapeutically targeting PD-1 molecules on immune cells, especially cytotoxic CD8^+^ T-cells, is one of the most successful strategies to treat tumors^10^. Understanding the mechanisms that regulate PD-1 expression on cytotoxic CD8^+^ T-cells presents important clues to optimize current tumor immunotherapy strategies, including CAR-T/TCR-T and immune checkpoint inhibitors, especially anti-PD-1 and anti-PD-L1 blocking antibodies. By genome-wide CRISPR/Cas9 screenings, we systematically identified genes and pathways regulating PD-1 expression in primary CD8^+^ T-cells. Subsequent custom screenings revealed that knockout of B4GALT1, a key gene in N-glycan biosynthesis, enhances T-cell activation and functions both in vitro and in vivo. Interestingly, we demonstrated that inactivation of B4GALT1 led to defective transfer of galectin-1 from cancer cells to the CD8^+^ T-cell surface, which represents a new type of immune checkpoint regulation. Finally, systematic intravenous administration or tumor-specific de novo synthesis of lactose, a structure-mimicking competitive inhibitor of galectins, could be used as an effective tumor immunotherapy strategy. This strategy has been tested in several preclinical mouse and human tumor models.

It is well accepted that the immune system plays critical roles in all stages of cancer development^52^. In immunoproficient patients, transformed cancer cells escape immunological controls via multiple mechanisms. Cancer cells can intrinsically reduce immune recognition, such as neoantigen presentation^53,54^. On the other hand, cancer cells also activate immune checkpoints to suppress the immune response within the tumor microenvironment. Recent advances in tumor immunotherapy have proven that suppression of immune checkpoints, such as anti-PD-1 and anti-PD-L1 antibodies, leads to efficient immune rejection of tumors^9,10^. The fact that only a few patients respond to anti-PD-1/PD-L1 antibodies suggests the existence of additional immune checkpoints. Indeed, malignant cancer cells can also actively modulate the immunosuppressive tumor microenvironment via secretion of cytokines and metabolites^55–57^. Galectins are a family of conserved galactosylation-binding proteins that have been proposed to modulate the functions of immune cells during normal physiological processes and various diseases, including tumors^58,59^. Our results support the model that cancer cells could transfer aberrantly high levels of cell surface Galectin-1 to proximal immune cells, especially tumor-targeting cytolytic CD8^+^ T-cells (Fig. 6). Subsequently, tumor-derived Galectin-1 interacts with galactosylated surface proteins on cytolytic CD8^+^ T-cells such as TCRs^33,34^ to regulate T-cell activation and functions. Targeting this novel immune checkpoint would be a new strategy for tumor immunotherapy. Indeed, targeted alteration of protein galactosylation by knockout of B4GATL1 leads to reduced tumor-transferred Gal-1 on CD8^+^ T-cells and enhanced T-cell activation and functions. On the other hand, lactose and its derivatives could inhibit such immune checkpoints by depleting cell surface Gal-1 on both cancer cells and immune cells. Lactose treatment activates infiltrated CD8^+^ T-cells in tumors mainly by depleting Gal-1 on CD8^+^ T-cells and relieving its repression of T-cell activation. Systematic administration of lactose or de novo synthesis of lactose within the tumor microenvironment by overexpressing LALBA could enhance CD8^+^ T-cell-mediated immunotherapy in various mouse and human preclinical tumor models.

**Figure 6.**
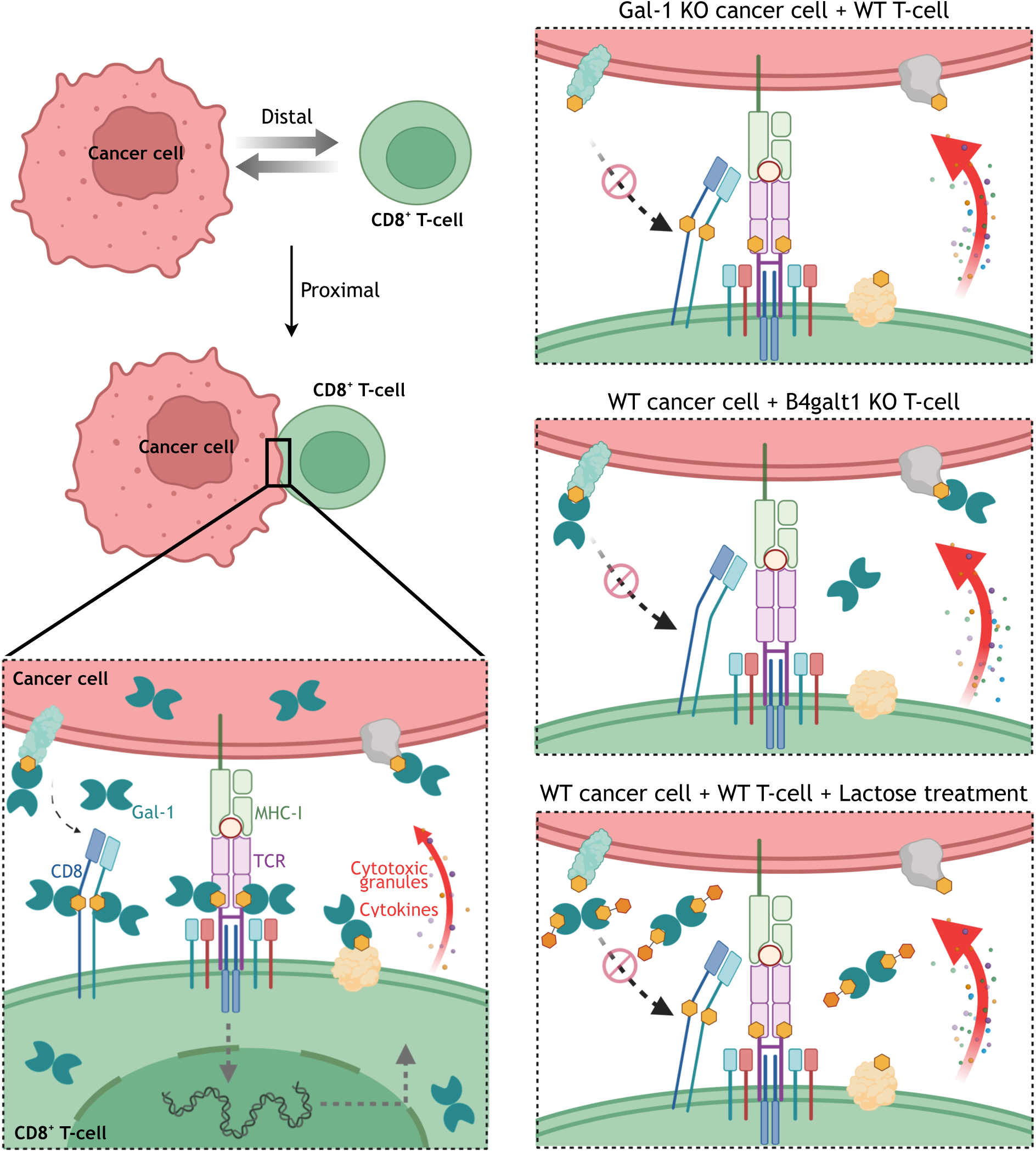
Schematic view of targeting a novel immune checkpoint involving N-glycome and galectins by lactose. Cancer cells transfer their surface Gal-1 proteins to proximal CD8^+^ T-cells, where Gal-1 proteins recognize and bind to galactosylated membrane proteins to suppress T-cell-mediated cytolysis by interfering with interactions between TCR and CD8. There are multiple strategies to target this immune checkpoint pathway: 1, Gal-1 deficiency in cancer cells; 2, B4GALT1 deficiency in CD8^+^ T-cells leads to reduced T-cell surface galactosylation and Gal-1 protein binding; and 3, lactose, a structure-mimicking competitive inhibitor of Gal-1 protein, relieves the inhibitory effect and functions as a novel immune checkpoint inhibitor.

As one of the mechanisms for cell-cell communication, intercellular exchange of cell-surface proteins has been involved in regulation of immune response^31,32^. Comparing with TCR-mediated trogocytosis^60^, the Gal-1 transfer from cancer cells to CD8^+^ T-cells is not dependent on TCR activation and interaction between TCR and MHC-peptide complex. Knockout of Rab27a and Coro1a genes in MC38 cells could not affect the cell surface Gal-1 level on MC38, nor transferring of Gal-1 to OT-1 cells, suggesting exosome pathway^61^ might not be involved (data not shown). Our results demonstrate that intercellular proximity, cell surface galactosylation level, and cell surface Gal-1 level are the major driving forces for intercellular Gal-1 transfer. Especially relative low affinity binding of Gal-1 with its galactosylated targets is intrinsic for this proximity-dependent intercellular protein spreading (PDICPS). A systematic analysis is necessary to identify additional substrates of PDICPS and their implications in cell-cell communication and biological processes.

Immune checkpoint blockade therapy demonstrates promising results in an increasing number of cancer patients. However, high cost is still one of the most important hurdles for its broad implication. More than 500 million metric tons of cow milk (containing approximately 4.8% lactose) are produced worldwide annually. From that, more than 6 million tons of dairy permeate powder (containing >76% lactose) have been produced in the food industry, and tens of thousands of tons of pharmaceutical grade lactose are used widely as a drug filler for medications. The antitumor efficacy and economic cost of lactose will provide a widely accessible strategy for cancer treatment.

## Supporting information

Supplemental figres 1-29

## Acknowledgments

We thank members of Y.Z. lab for helpful discussion and support. We also thank Dr. Guomin Li (CIMR) and Dr. Chuanyuan Li (CIMR) for their careful reading and invaluable feedback on the manuscript. We thank Dr. Yan Ma and Yang Cao at Metabolomics Center (NIBS), and Dr. Songbiao Zhu at Proteomics Center (CIMR) for assistance with LC-MS analysis, Yue Yin at Sequencing Center (NIBS) for single cell library preparation. The research in Y.Z. lab is supported by grants from National Key R&D Program of China (2021YFA1101002), National Natural Science Foundation of China (81773304, 81572795), the “Hundred, Thousand and Ten Thousand Talent Project” by Beijing municipal government (2019A39). We thank the municipal government of Beijing and the Ministry of Science and Technology of China for funds allocated to NIBS and CIMR.

## Author Contributions

Y.H. and Y.Z. conceived the study. Y.H., X.S., W.L., X.M. performed experiments and analyzed data. Y.Z. analyzed data and wrote the manuscript with support from all authors.

## Declaration of Interests

Part of this research has been submitted for patents.

## Methods

### Cell lines

HEK293T, CT26, and 4T1 cell lines were purchased from ATCC. B16F10 cell line was provided by laboratory of Dr. Ting Chen (National Institute of Biological Sciences, Beijing). MC38 cell line was provided by laboratory of Dr. Xiaodong Wang (National Institute of Biological Sciences, Beijing). Nalm6 cell line was provided by laboratory of Dr. Zhaoqing Ba (National Institute of Biological Sciences, Beijing). A375 cell line was provided by laboratory of Dr. Feng Shao (National Institute of Biological Sciences, Beijing). SGC7901 cell line was provided by Beijing Vitalstar Biotechnology Co., Ltd. PK136 hybridoma was purchased from ATCC for anti-mouse NK1.1 antibody production.

B16F10-OVA cell line was constructed by infecting B16F10 cells with lentivirus encoding chicken ovalbumin. MC38-Cas9 cells were generated by infecting MC38 cells with lentivirus encoding Cas9 protein and selected by blasticidin S (Gibco A1113903). MC38-EGFP and MC38-Gal-1-EGFP cell lines were generated by infecting MC38-Cas9 cells with EGFP-expressing and Gal-1-EGFP-expressing lentivirus, respectively. B2m knockout, Gal-1 knockout, and Gal-1/B2m double knockout MC38 cells were generated by infecting MC38-Cas9 cells with sgB2m-, sgGal-1-, and sgGal-1+sgB2m-expressing lentivirus, respectively. LALBA-overexpressing MC38 cells were generated by infecting MC38-Cas9 cells with lentivirus encoding LALBA cDNA and GFP^+^ cells were sorted as a pool. LALBA-overexpressing and B4GALT1 knockout MC38 cells were generated by infecting MC38-Cas9 cells with lentivirus encoding LALBA cDNA and B4galt1 sgRNA, and GFP^+^ cells were sorted as a pool.

B4galt1 rescue vectors were constructed based on pMSCV-U6 sgB4galt1-PGK-puro-2A-BFP vector. Long isoform (UniProt, P15535-1) and short isoform (UniProt, P15535-2) were amplified from cDNA of mouse T-cells, and used to replace the BFP fragment by seamless cloning kit (Biomed, CL116).

HEK293T, CT26, B16F10, MC38, A375 and 4T1 cells were cultured in DMEM (Gibco C11965500BT) supplemented with 10% fetal bovine serum (FBS), 2 mmol L-glutamine, 100 μg/ml penicillin, and 100 U/ml streptomycin (all purchased from Gibco). Nalm6 cells were maintained in RPMI-1640 (HyClone) medium supplemented with 15% FBS, 100 μg/ml penicillin, 100 U/ml streptomycin, 10 mM HEPES, 0.1 mM 2-mercaptoethanol, 2 mM L-glutamine and 1xMEM nonessential amino acids. SGC7901 cells were maintained in RPMI-1640 (HyClone) medium supplemented with 10% FBS, 100 μg/ml penicillin, and 100 U/ml streptomycin.

### Lentivirus and adeno-associated virus (AAV) preparation

Lentivirus was packaged by co-transfecting HEK293T cells with lentiviral vector, psPAX2, and pMD2.G. 48 hours post-transfection, supernatant was collected and filtered through a 0.45 μm filter to remove cell debris. Virus was used for infection directly or concentrated by centrifugation at 25,000 rpm (107,000 g) for 2.5 hours at 4 °C and resuspended in PBS.

To produce lentivirus for human CD8^+^ T-cell infection, HEK293T cells were seeded in Opti-MEM I Reduced Serum Medium (Gibco, 31985-070) supplemented with 5% FBS, 1 mM sodium pyruvate (Gibco, 11360-070), and 1xMEM nonessential amino acids (Gibco, 11140-050) the day before transfection. 6 hours post-transfection, the transfection medium was replaced with fresh medium supplemented with 1x ViralBoost (Alstem Bio, VB100). Lentivirus was collected and filtered through a 0.45 μm filter 24 hours and 48 hours after transfection separately, followed by addition of Lenti-X-Concentrator (Takara, 631232). Lentivirus was concentrated following manufacturer’s instructions and resuspended in X-VIVO 15 medium (Lonza, 04-418Q) in 1% of the original volume.

HEK293T cells were transfected with AAV8 vector along with packaging plasmid and adenovirus helper plasmid. After 72 hours, medium and cells were collected for precipitation and lysis, followed by density gradient centrifugation with iodixanol. The purified virus was then ultrafiltered and diluted with PBS.

### Animals

Female C57BL/6J and BALB/c mice were purchased from Beijing Vital River Laboratory Animal Technology Co., Ltd. OT-1 mice were provided by laboratory of Dr. Hai Qi (Tsinghua University). Cas9-EGFP/OT-1 mice were generated by breeding OT-1 and Cas9-EGFP knock-in mice ^62^ at animal facility of the National Institute of Biological Sciences, Beijing. NOD-Prkdcscid Il2rgtm1/Vst (NPG) mice were purchased from Beijing Vitalstar Biotechnology Co., Ltd. Six- to eight-week-old mice were used at the start of experiments. Animals were housed under specific pathogen-free conditions in individually ventilated cages in a controlled 12-h day-night cycle with standard food and water provided ad libitum. All animal experiments were conducted following the Ministry of Health national guidelines for housing and care of laboratory animals and performed in accordance with institutional regulations after review and approval by the Institutional Animal Care and Use Committee at the National Institute of Biological Sciences.

### Retroviral whole-genome CRISPR/Cas9 gRNA library construction

A whole-genome CRISPR knockout gRNA library (1000000096) was purchased from Addgene. The gRNA regions were PCR amplified with primer pair F:5’-GGCTTTATATATCTTGTGGAAAGGACGAAACACCG-3’ and R:5’-CTAGCCTTATTTTAACTTGCTATTTCTAGCTCTAAAAC-3’, and then transferred into MSCV-gRNA-PGK-PURO-2A-BFP vector by Gibson reaction.

### Retroviral small custom gRNA library construction

A total of 1,398 genes were selected according to whole-genome knockout screening results. On average, three gRNAs were selected from the initial library for each gene according to gRNA performance (Supplementary Table 3). A total of 105 intergenic control gRNAs were also added. Oligonucleotides containing the guide sequence were synthesized (Custom Array), PCR-amplified, and cloned into MSCV-gRNA-PGK-PURO-2A-BFP vector via Gibson reaction.

### Ex vivo T memory cell culture, infection, and adoptive transfer

At Day 0, splenocytes were prepared from 6- to 8-week-old Cas9-EGFP/OT-1 female mice and cultured with IL2 (10 ng/ml) and SIINFEKL peptide (10 ng/ml) in complete RPMI media (RPMI 1640, 10% FBS, 20 mM HEPES, 1 mM sodium pyruvate, 0.05 mM 2-mercaptoethanol, 2 mM L-glutamine, 100 U/ml streptomycin, and 100 μg/ml penicillin) at a density of 1×10^6^ cells/ml for 24 hours. On day 1, activated Cas9-EGFP/OT-1 T-cells were enriched by Percoll isolation as previously described^63^ and spin-infected (2,000 g, 30 °C, 1 hour, with no acceleration or brake) with retrovirus supplemented with polybrene (8 μg/ml) in 24-well-plate. After spin-infection, the plate was placed into a CO_2_ incubator at 37 °C for 5 hours and cultured with IL2 (2 ng/ml), IL7 (2.5 ng/ml), and IL15 (10 ng/ml) in complete RPMI media at a density of 3×10^5^ cells/ml. Two days after infection (Day 3), cells were selected by 3 μg/ml puromycin in the presence of IL2/IL7/IL15 for another 4 days. Day 7 cells were used for coculture experiments or adoptive transfer.

### Human T-cell isolation, culture and transduction

Human peripheral blood mononuclear cells (PBMC) were acquired from healthy donors (LV-BIOTECH). CD8^+^ T-cells were isolated using EasySep negative selection kit (STEMCELL, Cat#17953), and stimulated with anti-CD3/CD28 Dynabeads (Thermo Fisher Scientific, Cat#40203D). T-cells were cultured in X-VIVO medium with 10% FBS, 2 mmol L-glutamine, 100 μg/ml penicillin, 100 U/ml streptomycin, and human IL2 (500 IU/ml). 48 hours after activation, T-cells were transduced in a lentivirus-coated plate, centrifuged at 1,200 g for 90 minutes at 37°C. Lentivirus transduction was repeated once 24 hours later. Positive transduced cells were sorted 3 days later. During T-cell expansion, the cells were maintained at a concentration of 3 x 10^5^ cells/ml.

### Anti-CD19-CAR and NY-ESO-1 specific TCR T-cell generation

pMSCV-CD19 scFv (FMC63)-IRES-RFP-U6 sgRNA vector was constructed from a pMSCV-CD19 scFv-IRES-RFP vector provided by laboratory of Dr. Feng Shao (National Institute of Biological Sciences, Beijing). An U6 sgRNA cassette was assembled by seamless cloning kit (Biomed, CL116).

CD8^+^ T-cells were purified from splenocytes of Cas9-EGFP knock-in mice by a mouse CD8^+^ T-cell isolation kit (R&D, MAGM203). At day 0, CD8^+^ T-cells were stimulated with anti-CD3 (1 μg/ml, Cat#14-0031-86) and anti-CD28 (0.5 μg/ml, Cat#102116) in complete RPMI 1640 medium containing 20 ng/ml IL2 for 24 hours. On day 1, activated CD8^+^ T-cells were enriched by Percoll isolation and spin-infected (2,000 g, 30 °C, 1 hour, with no acceleration or brake) with retrovirus expressing anti-CD19-CAR and sgRNA supplemented with polybrene (8 μg/ml) in 24-well-plate. The RFP^+^ cells were sorted on day 3 for further culture. CD8^+^ T-cells were cultured with the same condition as T memory cells. In vitro killing assay was perform on day 7 by mixing anti-CD19-CAR T-cells with Nalm6.

NY-ESO-1 specific TCR (1G4)^64^ was synthesized and cloned into a lentiviral backbone with a SFFV promoter. T2A-BFP fragment and U6 shRNA cassette were assembled by seamless cloning kit (Biomed, CL116) simultaneously.

NY-ESO-1 specific TCR T-cells were generated following the human CD8^+^ T-cell isolation, culture, and transduction protocols. 3 days post-transduction, BFP^+^ cells were sorted for further culture. 7 days post-transduction, in vitro killing assay was performed by mixing T-cells with A375 cells.

### CRISPR/Cas9 screening

For ex vivo PD-1 expression screening, at Day 6, 3×10^6^ B16F10-OVA cells were plated in 15 cm dishes in DMEM (DMEM+10% FBS+ Pen/Strep). On Day 7, puromycin-selected OT-1 T-cells were resuspended in a final concentration of 6×10^5^ cells/ml with complete RPMI 1640 medium supplemented with IL2 (2 ng/ml), IL7 (2.5 ng/ml), and IL15 (10 ng/ml), and added to B16F10-OVA cells at a T-cell:B16F10-OVA cell ratio of 4:1. Cells were cocultured overnight at 37 °C. CD8^+^ T-cells were then collected and stained with PE anti-PD-1 in 2% FBS/PBS for 30 minutes on ice. The highest and lowest 5% PD-1^+^ cells were sorted via BD FACSAria.

For the secondary small custom library in vivo screening, following 4-day puromycin selection, an aliquot of 2×10^6^ infected OT-1 T-cells was saved as input (approximately 269X cell coverage per sgRNA). Library-infected OT-1 T-cells (2×10^6^ cells per recipient) were intravenously transferred into mice bearing Day 14 B16F10-OVA tumors. In total, 24 mice were used as recipients. At 7 days post-adoptive transfer, B16F10-OVA tumors were digested into single-cell suspensions, and tumor-infiltrated CD8^+^ T-cells were isolated by biotin anti-mouse CD8α (Biolegend, 100704) and streptavidin beads. Meanwhile, a 1/20 volume aliquot of single-cell suspension was used for OT-1 staining to estimate numbers of infiltrated OT-1 T-cells in each tumor. A total of 1×10^4^ to 1×10^5^ OT-1 T-cells were collected per tumor.

### Sequencing library preparation

Genomic DNA was extracted by phenol/chloroform extraction. Primary PCR was performed using Titanium Taq DNA Polymerase (Clontech, 639242) to amplify the sgRNA region. A secondary PCR was performed to add adaptors and indexes to each sample. Nova-seq 150-bp paired-end sequencing (Illumina) was performed.

### Analysis of screening results

Raw reads were preprocessed with sequence-grooming tools to remove adaptor sequences with Cutadapt (version 3.4)^65^ and to be merged with FLASH (version 1.2.11)^66^. Then, MAGeCK (version 0.5.9.5)^67^ was used to analyze the screening data. We used the MAGeCK “count” command to generate read counts of all samples. The raw read counts of all gRNAs for all samples were merged into a count matrix. We next used the MAGeCK “test” command to identify the top negatively and positively selected gRNAs or genes with default settings with software available from https://sourceforge.net/projects/mageck/. GSEA in the Kyoto Encyclopedia of Genes and Genomes (KEGG) functional pathway was performed using the GSEA() function of the R package clusterProfiler (version 3.12.0)^68^ with default parameters. The KEGG database was selected in the “C2” category of the R package msigdbr (version 7.5.1) (https://igordot.github.io/msigdbr/).

### Flow cytometry

For surface staining, cells were stained in 2% FBS/PBS on ice for 30 minutes. Intracellular staining was performed with a fixation/permeabilization kit (BD Biosciences) according to manufacturer’s instructions. The following antibodies were used: APC anti-mouse CD8α (Invitrogen, 17-0081-83), PE anti-mouse CD279 (PD-1) (BioLegend, 109104), APC anti-mouse CD279 (PD-1) (BioLegend, 109112), FITC anti-mouse IFNγ (BioLegend, 505806), FITC anti-mouse CD3 (BioLegend, 100204), PE-streptavidin (BioLegend, 405203), biotinylated erythrina cristagalli lectin (B-1145-5), biotinylated succinylated wheat germ agglutinin (B-1025S-5), biotinylated recombinant Gal-1, PE anti-mouse Galectin-1 (R&D, Cat#IC1245P), PE anti-mouse TCR Vα2 (BD, Cat#553289), APC anti-human CD8α (BioLegend, 300912), PE anti-human IFNγ (BioLegend, 506507), and FITC anti-mouse TCR Vα2 (BioLegend, 127806). Flow cytometry was performed by BD FACSAria and data were analyzed with FlowJo.

### ELISA

OT-1 T-cells (3×10^5^/ml) were cocultured with B16F10-OVA cells at 37 °C for 8 hours in the presence of IL2 (2 ng/ml), IL7 (2.5 ng/ml), and IL15 (10 ng/ml). Supernatants were collected after coculture. TNFα and IFNγ in the culture supernatant were measured using ELISA kits (ABclonal, Cat#RK00027, Cat#RK00019). Samples were plated in duplicate.

Human CD8^+^ T-cells (3 x 10^5^/ml) were cocultured with A375 cells at 37 °C for 24 hours in the presence of human IL2 (500 IU/ml). Supernatants were collected for measurement of TNFα and IFNγ secretion using ELISA kits (ABclonal, Cat#RK00030, Cat#RK00015).

### Mouse tumor models, drug treatment, and adoptive transfer experiments

4×10^5^ B16F10-OVA cells were injected subcutaneously into female C57BL/6J mice. Seven days post-injection, mice bearing tumors of similar size were randomly separated into 2 groups. A total of 2×10^6^ B4galt1 gRNA-transduced or control gRNA-transduced OT-1 T-cells were injected intravenously. Tumors were then measured every two days with an electronic digital caliper. Tumor volume was calculated as width x width x length x 1/2.

For lactose treatment, MC38 (1×10^6^ cells), B16F10 (4×10^5^ cells), B16F10-OVA (4×10^5^ cells), B16F10-Control (4×10^5^ cells), 4T1 (4×10^5^ cells), CT26 (4×10^5^ cells), and SGC7901 (5×10^6^ cells) cells were injected subcutaneously into female C57BL/6J, BALB/c, or NPG mice. The next day, unless otherwise indicated, 250 mM lactose in 200 μl PBS was injected intravenously every other day (until Day 21). For LacNAc treatment, 20 mM LacNAc in 200 μl PBS was injected intravenously every other day (until Day 21).

For the combination therapy, anti-PD-1 (Clone RMP1-14, BioXCell, BE0146) or anti-PD-L1 (Clone 10F.9G2, BioXCell, BE0101) at 100 μg each in 200 μl PBS was injected intraperitoneally on Days 6, 9, and 12 after tumor implantation.

For CD8^+^ T-, CD4^+^ T-, NK-cell depletion, 100 μg NK depletion antibody in 200 μl PBS was injected intraperitoneally at day −3 and day −1 prior to tumor inoculation; 250 μg CD8 depletion antibody (Clone 2.43, BioXCell, BE0061) in 200 μl PBS was injected intraperitoneally at day −2 and day −1 prior to tumor inoculation; 400 μg CD4 depletion antibody (Clone GK1.5, BioXCell, BE0003-1) in 200 μl PBS was injected intraperitoneally at day −2 and day −1 prior to tumor inoculation, respectively.

For the MC38 rechallenge tumor model, 1×10^6^ MC38-EGFP cells or MC38-LALBA cells were injected subcutaneously into female C57BL/6J mice. Twenty-one days after tumor inoculation, mice that completely cleared the MC38-LALBA tumors were rechallenged with 1×10^6^ MC38-EGFP cells. A control group of age-matched female C57BL/6J mice that had not been exposed to MC38-EGFP cells (naive) was used as control.

For the mixture of MC38-EGFP/MC38-LALBA tumor model, MC38-EGFP and MC38-LALBA cells were mixed at ratios of 9:1 or 1:1, and a total of 1×10^6^ cells were injected subcutaneously into female C57BL/6J mice. Mice injected with 1×10^6^ MC38-EGFP cells were included as control.

For intratumoral virus injection, 1×10^6^ MC38 cells were injected subcutaneously into female C57BL/6J mice. 9 days after tumor inoculation, mice with similar tumor sizes were used for intratumoral virus injection. Injection was performed on Days 9, 12, and 15. The titer of the lentiviral or AAV virus was determined by serial dilution method using MC38 cells. For each injection, the same units of concentrated lentiviral or AAV virus (defined by infection of 8×10^4^ MC38 cells in vitro) were injected into the tumors. Tumor growth was monitored every 3 days up to Day 24.

### Mice with a humanized immune system

To generate mice with a humanized immune system, 3- to 4-week-old NPG mice received a sublethal irradiation of 120 cGy followed by a tail vein injection of human CD34^+^ cord blood cells. 12 weeks after transfer, peripheral blood was analyzed for presence of human CD45^+^ cells to confirm human hematopoietic reconstitution. The mice were inoculated subcutaneously with 5 × 10^6^ SGC7901 cells at 13 weeks after hematopoietic stem cell transplantation. From Day 1 onward, the mice were intravenously injected with 250 mM lactose in 200 μl PBS as described above. Another group of mice injected with PBS was included as control. From Day 10 onward, anti-human PD-1 antibody (Pembrolizumab, Selleck, Cat# A2005) (10 mg/kg) was injected intraperitoneally twice a week for total 4 times. Tumor growth was measured every 2 or 3 days.

### Tumor-infiltrated lymphocyte (TIL) isolation

B16F10-OVA, MC38, and SGC7901 tumors were cut into small pieces in 6-well plates containing 5 ml RPMI 1640, 2% FBS, and 50 U/ml collagenase type IV (Invitrogen, V900893). Samples were incubated at 37 °C for 1 hour with rotation. Suspensions were passed through a 70 μm strainer and washed three times with PBS. Samples were then used for antibody staining and FACS analysis.

### T-cell infiltration and proliferation within tumors

Following 4-day puromycin selection, infected OT-1 T-cells were labeled with CFSE (carboxyfluorescein diacetate succinimidyl ester, Invitrogen), and 2×10^6^ labeled cells were intravenously transferred into Day 14 B16F10-OVA tumor-bearing mice. CFSE dilution was quantified by flow cytometry at 24 hours and Day 6 following transfer.

### In vitro T-cell activation by anti-CD3/CD28 antibodies

In vitro cultured Tmem cells at 1×10^5^ cells/ml were stimulated with anti-CD3 (1 μg/ml, Cat#14-0031-86) and anti-CD28 (0.5 μg/ml, Cat#102116) in complete RPMI 1640 medium containing 10 ng/ml IL2 for 6~8 hours. Tumor derived PD-1 positive CD8^+^ T-cells were sorted by FACS and stimulated at the same condition as mentioned above.

To test the effect of recombinant Gal-1 (rGal-1), T-cells were incubated with 2.5 μg/ml rGal-1 (Cat# 50100-MNAE) for 10 minutes before stimulation.

To load T-cells with endogenous Gal-1 from MC38 cells, 2×10^6^ OT-1 T-cells were cocultured with 1×10^6^ wild-type or Gal-1 knockout MC38 cells in conditional medium (50% of T-cell medium without cytokine, 50% of MC38 cell conditional medium) in 6-cm-dish for 8 hours. OT-1 T-cells were sorted by FACS.

### Gal-1 transfer assay

#### Coculture

OT-1 T-cells were cocultured with wild-type or Gal-1 knockout MC38 cells at a 2:1 ratio or indicated ratios for 8 hours in fresh medium, or wild-type or Gal-1 knockout MC38 cell conditional medium.

#### Boyden chamber

OT-1 T-cells were seeded onto the upper compartment of the Boyden chamber (Corning, Cat#3412) with MC38 cells on the lower compartment, or onto the lower compartment together with MC38 cells for an 8-hour culture.

#### Separation by agarose layer

Low-melting agarose (Cat#16520-050) dissolved in culture medium was added after MC38 cell attachment. Then OT-1 T-cells were added on the agarose layer or directly on MC38 cells. The cells were cultured for 8 hours.

### In vitro targeted cell killing

For OT-1 T-cell killing activity assay, B16F10-OVA cells and B16F10 cells were labeled as CFSE^hi^ (2.5 mM) and CFSE^lo^ (50 nM), respectively, and then cocultured with OT-1 T-cells in 96-well plates at the indicated ratios. Cancer cells without addition of OT-1 T-cells were used as controls. Following 24 hours of incubation, the ratios of CFSE^hi^ and CFSE^lo^ populations were detected by FACS.

For in vitro killing of MC38 cells, the cells were pulsed with 100 ng/ml Ova-peptide (SIINFEKL) at 37 °C for 3 hours prior to coculture with OT-1 T-cells.

For lactose treatment, in vitro killing experiment was performed with conditional medium (50% fresh T-cell medium without cytokine and 50% MC38 conditional medium) with or without 20 mM lactose. Killing result was checked by FACS 20 hours post-incubation.

Specific killing was calculated by following equation: specific killing percentage = [1-(CFSE^hi^/CFSE^lo^ of T-cell)/(CFSE^hi^/CFSE^lo^ of cancer cell only)] × 100%.

### FRET

TCR-CD8 FRET was measured by flow cytometry^34^. For rGal-1 treatment, 1.2×10^6^ control or B4GALT1 KO Tmem cells were treated with 2.5 μg rGal-1 in 100 μl PBS at room temperature for 10 minutes. For lactose treatment, 20 mM lactose were added during rGal-1 treatment. T-cells were then incubated with PE-anti Vα2 and APC-anti CD8α at 4°C for 30 minutes. Samples were stained with either antibody, both or neither. After staining, samples were fixed with fixation buffer at 4°C for 10 minutes. FRET emission was assessed and FRET efficiency was calculated in FRET units as reported previously^34^.

### Proteomic analysis of Gal-1 binding proteins in CD8^+^ T-cells

Recombinant Gal-1 was purified^69^ and coupled with sepharose beads (Cat#GE17-0906-01). B4GALT1 knockout OT-1 T-cells were cocultured with MC38 for 8 hours and then stained by anti-Gal-1 antibody for sorting of Gal-1 negative OT-1 T-cell population. Membrane extracts of OT-1 T-cells were prepared by Mem-PER^TM^ Plus Kit (Cat#89842Y). Extracts were incubated with Gal-1-sepharose beads. After washing, binding proteins were eluted by 20 mM lactose and precipitated by DOC/TCA, and then separated on SDS-PAGE followed by silver staining (Sigma, PROTSIL1). The samples were digested and subjected to LC-MS (LTQ ORBITRAP Velos mass spectrometer, Thermo Fisher Scientific, San Jose, CA, USA).

Raw data was processed by Proteome Discoverer (version1.4, https://www.thermofisher.com/hk/en/home/industrial/mass-spectrometry/liquid-chromatography-mass-spectrometry-lc-ms/lc-ms-software/multi-omics-data-analysis/proteome-discoverer-software.html) to obtain a PSM (peptide spectrum matches) raw intensity table of all identified proteins for further analysis with R package DEqMS^70^. Raw intensity values were log2 transformed, and for each PSM, the median of log2 intensity was subtracted to get a relative log2 ratio. Then different expression of protein calculation was enabled in eBayes() function with method ROC (receiver operating characteristic) analysis. Differentially expressed proteins were selected by criteria P < 0.05. Those proteins were used for functional enrichment analysis with R package ClusterProfiler (version 3.12.0)^71^.

### Western blot

10 μg cell membrane extracts of OT-1 T-cells were separated by SDS–PAGE gels. Gels were transferred to nitrocellulose membranes (Amersham™ Protran®, Cat#10600001). Anti-CD8β (Abcam, Cat# ab228965) (1:1000), Anti-Ly9 (Abcam, Cat# ab252931) (1:1000), anti-Igf2γ (Sino Biological, Cat# 107533-T40) (1:500), anti-Itga1 (Solarbio, Cat# K002895P) (1:500), anti-Itgb7 (Solarbio, Cat#K004272P) (1:500), anti-Ighg1 (Solarbio, Cat#K111394P) (1:500), anti-Lnpep (Santa Cruz, Cat#sc-365300) (1:100), and anti-Sell (Santa Cruz, Cat#sc-390756) (1:100) were used as primary antibodies. Secondary antibodies were IRDye® 680RD Donkey anti-Mouse IgG (LI-COR, P/N 926-68072) (1:10000), IRDye® 800CW Goat anti-Mouse IgG (LI-COR, P/N 926–32210) (1:10000), IRDye® 680RD Goat anti-Rabbit IgG (LI-COR, P/N 926-68071) (1:10000), IRDye® 800CW Goat anti-Rabbit IgG (LI-COR, P/N 926-32211) (1:10000). The membranes were scanned on an Odyssey imager (LI-COR).

### Lactose quantification by LC‒MS

Cells were collected and washed twice with PBS. For 2×10^6^ cells, 400 μl methanol/acetonitrile mixture (methanol:acetonitrile:H_2_O=4:4:2) was added and vortexed. The supernatant was placed at −80 °C for 1 h, and the tubes were spun at 14,000 × g for 15 minutes at 4 °C. The supernatant was transferred to a new tube and the last step was repeated once. The final supernatant was used for analysis.

Cell medium was collected and passed through a 0.45 μm filter to remove cell debris. Methanol was added at 2.5 times the volume of medium. The tubes were vortexed for 2 minutes and spun at 14,000 × g for 15 min. The supernatant was used for analysis.

A solution of lactose purchased from Sigma‒Aldrich at 100 mM was used to prepare working standard solutions of 100, 200, 500, 1000, 2000, 5000, and 10000 nM by serial dilution in 50% methanol in water. The standard solutions and biological extracts were transferred to LC vials with microinsert for analysis. LC‒MS analysis was performed on a Thermo Vanquish UHPLC equipped with a Thermo Q Exactive HF-X hybrid quadrupole-Orbitrap mass spectrometer in negative ESI mode. Separation was achieved on a Waters Acquity UPLC BEH Amide column (1.7 μm, 2.1 × 100 mm) at a column temperature of 35 °C under isocratic elution of 30% mobile phase A (5 mM ammonium formate in water) and 70% mobile phase B (acetonitrile). The flow rate was 0.3 ml/min, and the injection volume was 10 μl. The run time for each injection was 5 minutes.

Full-scan mass spectra were acquired in the range of m/z 66.7 to 1000 with the following ESI source settings: spray voltage: 2.5 kV, auxiliary gas heater temperature: 380 °C, capillary temperature: 320 °C, sheath gas flow rate: 30 units, auxiliary gas flow: 10 units. MS1 scan parameters included resolution 60000, AGC target 3e6, and maximum injection time 200 ms. Data processing was performed with Thermo Xcalibur software (version 4.2), and the [M + FA]-adduct of lactose (m/z 387.1138) was used for quantitation.

Note: The concentrations showed in figure S25 represent all detectable disaccharides with MW 342.3.

### Lactose-BSA coupling

Lactose-BSA was prepared following published protocol^43^. Briefly, 20 mg/ml BSA and 80 mg/ml lactose solutions were mixed at 1:1 volume ratio and lyophilized, which was then placed in an air oven at 120 °C for 30 minutes with the tube left open. The tube was allowed to cool down to room temperature. The powder was dissolved in water. Low molecular weight carbohydrates were removed by multiple dialysis.

### RT-qPCR and RNA-seq

Total RNA was extracted using TRIzol® Reagent (Ambion, Cat# 15596018). cDNA was synthesized by the ImProm-II™ Reverse Transcriptase system (Promega, Cat# A3801) using 100 ng RNA per reaction. The qPCR reactions were prepared with TB Green® Premix Ex Taq™ (Takara, Cat# RR420A) using 1 μl cDNA per reaction in a 20 μl reaction volume. The relative gene expression levels were normalized to GAPDH.

Total RNA was used as input material for the RNA sample preparations. Sequencing libraries were generated using NEBNext Ultra RNA Library Prep Kit for Illumina (NEB, Catalog #E7530L) following manufacturer’s recommendations and index codes were added to attribute sequences to each sample. Prepared libraries were quantified and sequenced by the Illumina NovaSeq 6000 S4 platform with 150 bp paired-end reads.

### RNA-Seq data analysis

Raw data was aligned to mouse reference genome (mm10) with Ensemble version 98 gene annotation by TopHat (version 2.1.1) (TopHat2: accurate alignment of transcriptomes in the presence of insertions, deletions and gene fusions). Then raw count of each gene was calculated by HTSeq (version 1.99.2)^72^ and normalized to Fragments Per Kilobase of transcript per Million mapped reads (FPKM) by cufflinks (version 2.2.1) (Differential gene and transcript expression analysis of RNA-seq experiments with TopHat and Cufflinks). A R-based toolkit, DESeq2 (version 1.38.0)^73^, was used for defining differentially expressed genes (DEGs) that selected by criteria P < 0.05 or P < 0.01. Those DEGs were performed functional enrichment analysis with R package ClusterProfiler (version 3.12.0)^71^. The average gene expression levels were used to estimate the signatures in a bulk-seq sample. Full lists of genes involved in each signature calculation are listed in Supplementary Table 6.

### Single cell RNA-seq sample preparation

CD3-positive cells were obtained from three pairs of PBS- or lactose-treated MC38 tumors. The same numbers of CD3^+^ cells from each tumor were pooled together and subjected to FACS sorting. The FACS-sorted viable CD3^+^ cells were resuspended at 5 × 10^5^–1 × 10^6^ cells/ml with a final viability of >85%. Single-cell library preparation was carried out using the 5′ V(D)J and gene expression platform as per the 10× Genomics protocol (10×Genomics, Pleasanton, CA, USA). Libraries were sequenced by NovaSeq 6000 S4 platform.

### Single-cell RNA-Seq data processing

Raw reads were processed to generate gene expression matrices by Cell Ranger toolkit (version 7.0.0, 10x Genomics, https://support.10xgenomics.com/single-cell-gene-expression/software/pipelines/7.0/what-is-cell-ranger), to remove low-quality reads, to aligned the mouse reference genome (mm10, version=”2020-A”) with Ensemble version 98 gene annotation, to assign cell barcodes, and to generate the unique molecular identifier (UMI) matrix.

A R-based toolkit, Seurat (version 4.3.0, https://doi.org/10.1016/j.cell.2021.04.048), was used to analyze the single-cell RNA-Seq (scRNA-seq) data. Specifically, the raw UMI matrix was processed to filter out genes detected in less than 3 cells, to remove cells that had either lower than 200 or higher than 5,000 expressed genes. We further quantified the numbers of gene and UMI count for each cell and kept high quality cells with less than 50% mitochondrial gene counts. R package DoubletFinder(version 2.0.3) was then applied to each sequencing library to remove potential doublets; cells with expected doublet rate of 7.5% were filtered out, to ensure that most of the CD3^+^ cells were included. Finally, 19,776 cells (9,516 from PBS and 10,260 from LAC) were obtained for downstream analysis.

### TCR sequences assembly and Quality control

Function vdj from Cell Ranger toolkit (version 7.0.0) provided by 10x Genomics was applied to align TCR-seq reads to the mouse reference genome (mm10, version=”2020-A”) and assemble TCR sequences. The preliminary TCR sequences were filtered to keep those characterized with high confident, full-length, productive, and assigned with a valid cell barcode and an unambiguous chain type. With R-based toolkit scRepertoire (version 1.8.0), each cell was assigned with one pair of alpha and beta chains with the highest UMI counts. Cells with identical TCR pairs were defined as clonal cells and considered to originate from the same ancestry. In total, 15,013 of T-cells (6,998 from PBS and 8,015 from LAC) were assigned with one pair of TCR sequence, and 14,428 cells (6,639 from PBS and 7,789 from LAC) of them had both TCR and scRNA-seq data.

### Major cell-type identification and TCR clonotype analysis

Seurat (version 4.3.0) was used to normalize expression matrices by function NormalizeData() with sctransform method and ScaleData() after filtration. Then R package ProjecTILs (version 3.0.0) was applied to identified different T-cell subtypes, function make.projection() was used to project all candidate cells onto a mouse interactive reference TIL Atlas. Function cellstate.predict() was used to summarizes TIL diversity using nine broad cell subtypes with distinct phenotypes, functions, metabolic lifestyles, and preferential tissue distributions^47^. At last, 14,344 cells (6,607 from PBS and 7,737 from LAC) were annotated in 9 subtypes and to perform further analysis with R package scRepertoire (version 1.8.0).

### Hematologic and biochemical analysis of blood samples

Blood samples (500 μl) were harvested in EDTA-coated tubes. A portion of the blood was taken for the evaluation of hematological parameters: white blood cells (WBC), lymphocytes (LYM), MID cells (MID), granulocytes (GRA), red blood cells (RBC), hemoglobin (HGB), hematocrit (HCT), mean corpuscular volume (MCV), mean corpuscular hemoglobin (MCH), mean corpuscular hemoglobin concentration (MCHC), red cell distribution width-standard deviation (RDW-SD), red cell distribution-coefficient of variation (RDW-CV), platelets (PLT), procalcitonin (PCT), mean platelet volume (MPV), and platelet distribution width (PDW) in a blood analyzer (TEK-II MINI Hematology analyzer). The remaining samples were centrifuged (1,500 g, 10 minutes, room temperature), and the supernatants were used for biochemical parameter analysis: alanine aminotransferase (ALT), aspartate aminotransferase (AST), alkaline phosphatase (ALP), creatinine (CR), blood urea nitrogen (BUN), lactate dehydrogenase (LDH), creatine kinase (CK), glucose (GLU), and inorganic phosphorus (Pi) were measured by an automated clinical biochemical analyzer (Beckman Coulter AU5800).

### Ex vivo culture of patient-derived tumor fragments

Tumor samples were collected from patients with lung cancer, colon cancer, rectal cancer, and esophageal cancer at Beijing Chaoyang Hospital and Guangdong Provincial People’s Hospital. Detailed patient characteristics were provided in supplementary figure 29. The study was performed following all relevant ethical regulation. All patients consented to research usage of material not required for diagnostic use either by opt-out procedure or via previous written informed consent.

Tumor lesions were collected in ice-cold collection medium (RPMI1640 (HyClone) supplemented with 2.5% FBS, 100 μg/ml penicillin, and 100 U/ml streptomycin) for subsequent processing. Tumor lesions were cut into slices of ~1 mm thickness on ice, followed by punching with a 2 mm-diameter-puncher. To prepare matrix, tumor growth medium (DMEM supplemented with 10% FBS, 100 μg/ml penicillin, 100 U/ml streptomycin, 2 mM L-glutamine, 1 mM sodium pyruvate, 1×MEM-NEAA, 50 μM β-mercaptoethanol), NaHCO_3_ in PBS (final conc. 1.1%), and collagen I solution (final conc. 1 mg/ml) were mixed slowly, followed by adding ice-cold Matrigel (Corning, Cat#354262). Next, pre-cooled 96-well-plate (flat-bottom) was coated with 40 μl matrix and solidified at 37°C to form a bottom layer. Tumor fragments were washed with tumor growth medium and placed on top of the pre-solidified matrix, with one fragment per well, followed by addition of a second layer of 40 μl matrix. After solidification at 37°C, 120 μl tumor growth medium containing lactose (final conc. 20 mM) or PD-1 antibody (Pembrolizumab, final conc. 10 μg/ml) was added on top of the matrix. Control cultures were carried out with only tumor growth medium. The tumor fragment cultures were kept at 37°C. 48 hours post-incubation, tumor fragments were collected in Trizol (Ambion, Cat#391309), and homogenized for RNA extraction and RT-qPCR.

The expression level of each gene was normalized by hGAPDH, and the fold-change (FC) between treatment and control was used to calculate the response score. Briefly, the average FCs of 10 genes (CCL17, CCL20, IFNγ, CCL4, CXCL9, CXCL10, CCL5, CD137, CXCL11, CXCL5) was used as response score for each sample.

### Statistical analysis

Statistical analyses were conducted with R 4.1.0. Unless otherwise stated, a two-tailed paired Student’s t test was used to determine statistical significance (*P < 0.05, **P < 0.01; ***P < 0.001; NS, not significant) with function t.test(). Two-way ANOVA was performed by function aov(). The mean and standard error of the mean (SEM) are presented in the figures, and the error bars represent the SEM value, and calculated by function sem(). Correlation of normalized values (z-score) was calculated by function scale(). Test for Correlation Between Paired Samples was calculated by function cor.test() with method of Pearson’s product moment correlation coefficient. Principal component analysis (PCA) was calculated by function PCA() of R package FactoMineR(version 2.4).

## Data availability

The raw sequence data reported in current study have been deposited in the Genome Sequence Archive in BIG Data Center, Beijing Institute of Genomics (BIG), Chinese Academy of Sciences, under accession numbers PRJCA010494 that can be accessed at https://ngdc.cncb.ac.cn/search/?dbId=&q=PRJCA010494.

## Supplemental information

Supplemental information includes 29 figures and 6 tables.

**Figure S1. Multiple components in the N-glycan biosynthesis pathway were identified in genome-wide screenings for PD-1 regulators in CD8^+^ T-cells.**

a. Verification of candidate genes by individual gRNAs. The relative expression levels of surface PD-1 protein and PD-1 mRNA were measured by FACS as the mean fluorescent intensity (MFI) and RT‒qPCR, respectively.

b. Schematic views of the N-glycan biosynthesis pathway. The genes identified in current genome-wide screenings are labeled.

c. Distribution of N-glycan biosynthesis-related genes in current screenings. The blue curve represents all target genes; the red curve represents all genes involved in the N-glycan biosynthesis pathway; and the gray curve represents control gRNAs.

d. Representative PD-1 FACS plots for gRNAs targeting the B4galt1, Mgat2, and Dpm3 genes.

**Figure S2. Ablation of B4GALT1 in CD8^+^ T-cells activates T-cell receptor (TCR) signaling.**

a-b. The effect of B4GALT1 deficiency on PD-1 surface expression could be rescued by overexpression of either the long or short isoform of B4GALT1 cDNA. The p values were calculated by a two-tailed Student’s t test.

c. Heatmap demonstrating differentially expressed genes (DEGs) between B4GALT1 knockout and control wild-type mouse OT-1 CD8^+^ T-cells after coculture. The genes in the TCR signaling pathway are labeled on the left side.

d. Volcano plot showing upregulated and downregulated genes (p value<0.01) in B4GALT1 knockout mouse OT-1 CD8^+^ T-cells after coculture. The genes in the TCR signaling pathway are labeled with dark blue and dark red. The top genes and some genes in the TCR signaling pathway are annotated. The p value was calculated by the Wald test, and p.adjust was calculated by the Benjamini‒Hochberg procedure with the R package DESeq2 (version 1.22.2).

e. Bar graph showing KEGG pathways significantly changed in B4GALT1 knockout mouse OT-1 CD8^+^ T-cells after coculture. The p value was calculated by the clusterProfiler (version 3.12.0) R package.

Data are shown as the mean ± SEM. **P < 0.01; NS, not significant.

**Figure S3. The effect of B4GALT1 knockout on tumor-infiltrated OT-1 T-cells.**

a. Tumor infiltration of CFSE-labeled OT-1 CD8^+^ T-cells 24 hours after transplantation. The p value was calculated by a two-tailed Student’s t test.

b. Enhanced proliferation of CFSE-labeled B4GALT1 knockout (sgB4galt1) OT-1 T-cells in tumors. The p value was calculated by a two-tailed Student’s t test.

Data are shown as the mean ± SEM. NS, not significant.

**Figure S4. CRISPR/Cas9 knockout of B4GALT1 (sgB4galt1) in OT-1 cells alters surface binding of lectins and galectin-1 (Gal-1).**

a-b. Cells were stained with biotin-ECL (a) or biotin-sWGA (b) and streptavidin-PE. Blue and red curves indicate control sgRNA- and sgB4galt1-infected OT-1 cells, respectively. Gray dotted curves indicate cells stained with streptavidin-PE only. The MFIs of each sample were measured by FACS. The p value was calculated by a two-tailed Student’s t test.

c. Blue and red bars indicate control sgRNA- and sgB4galt1-infected OT-1 cells, respectively, stained with biotin-rGal-1 and streptavidin-PE. The p value was calculated by a two-tailed Student’s t test.

Data are shown as the mean ± SEM. *P < 0.05; **P < 0.01; ***P < 0.001.

**Figure S5. Expression of galectins in OT-1 and MC38 cells.**

a. The mRNA expression levels of each galectin in OT-1 and MC38 cells are shown as FPKMs, which were generated by whole-genome RNA-sequencing analysis.

b. FACS analysis of surface and intracellular expression of Gal-1, Gal-3, and Gal-9 in OT-1 and MC38 cells.

c. For MC38 cells, the ratios of surface galectin MFIs versus intracellular galectin MFIs were calculated.

Data are shown as the mean ± SEM.

**Figure S6. In vitro B4GALT1 knockout does not enhance TCR activation by anti-CD3/28 antibodies.**

B4GALT1 knockout and control OT-1 cells (1×10^5^ cells/ml) were stimulated with anti-CD3 (1 μg/ml) and anti-CD28 (0.5 μg/ml) for 8 hours. The mRNA expression levels of Tnfα(a) and Ifnγ(b) were measured by quantitative RT‒ qPCR. The p value was calculated by a two-tailed Student’s t test.

Data are shown as the mean ± SEM. **P < 0.01. NS, not significant.

**Figure S7. MC38 cells transfer their surface Gal-1 to the OT-1 cell surface.**

OT-1 cells were cocultured with MC38 cells at 0:10, 1:10, 1:2, 2:1, 10:1 and 10:0 ratios. MFIs of surface Gal-1 on OT-1 and MC38 cells were measured by FACS with an anti-Gal-1 antibody.

Data are shown as the mean ± SEM.

**Figure S8. Close proximity is necessary for Gal-1 transfer from MC38 to OT-1 cells.**

a. OT-1 cells were cocultured with wild-type or Gal-1 knockout MC38 cells at a 2:1 ratio for 8 hours with MC38 cell conditional medium. MFIs of surface Gal-1 on OT-1 cells were measured by FACS with an anti-Gal-1 antibody. The p value was calculated by a two-tailed Student’s t test.

b. OT-1 cells were seeded onto the upper compartment of the Boyden chamber only (i), with MC38 cells on the lower compartment (ii), or onto the lower compartment together with MC38 cells (iii). MFIs of surface Gal-1 on OT-1 cells were measured by FACS with an anti-Gal-1 antibody. The p value was calculated by a two-tailed Student’s t test.

c. Low-melting agarose was added after MC38 cell attachment. Then, OT-1 cells were added to the well (i), to the agarose layer (ii), or directly to MC38 cells (iii). MFIs of surface Gal-1 on OT-1 cells were measured by FACS with an anti-Gal-1 antibody. The p value was calculated by a two-tailed Student’s t test.

Data are shown as the mean ± SEM. *P < 0.05; ***P < 0.001. NS, not significant.

**Figure S9. FACS analysis demonstrates successful depletion of CD8^+^ T-cells (a), CD4^+^ T-cells (b), and NK cells (c) in peripheral blood samples of wild-type mice by different antibodies.**

**Figure S10. B2m knockout rescues the growth of Gal-1 knockout MC38 tumors in wild-type mice.**

Gal-1/B2m double knockout and Gal-1 single knockout MC38 cells were inoculated subcutaneously into wild-type C57BL/6J mice (1×10^6^ MC38 cells/mouse). The p value was calculated by two-way ANOVA.

Data are shown as the mean ± SEM.

**Figure S11. FACS analysis of surface Gal-1 expression on CD8^+^ T-cells in spleen, peripheral blood, and MC38 tumor samples of wild-type mice.**

**Figure S12. B4GALT1 regulates TCR-CD8 colocalization and TCR activation via surface Gal-1.**

a. Schematic view of rGal-1 pulldown-MS with an rGal-1 affinity column.

b. Volcano plot showing identified Gal-1 binding proteins in wild-type and B4GALT1 knockout OT-1 cells. Proteins among the top list were annotated and labeled with red (decreased in B4GALT1 knockout) and green (increased in B4GALT1 knockout). Proteins in the TCR signaling pathway are underlined. The p values were calculated by Limma in DEqMS (V1.8.0).

c. Bar graph showing KEGG pathways significantly changed in B4GALT1 knockout OT-1 cells. The p value was calculated by the clusterProfiler (version 3.12.0) R package.

d. Western blot verification of the top list from the pulldown-MS data.

e. CD8β is a substrate of B4GALT1.

f. Schematic view of the FRET approach.

g. Fold change of FRET units for wild-type and B4GALT1 knockout OT-1 with or without rGal-1/rGal-1+lactose treatment was summarized. The p value was calculated by a two-tailed Student’s t test.

Data are shown as the mean ± SEM. ***P < 0.001. NS, not significant.

**Figure S13. Effect of B4GALT1 and CD8α on OT-1 CD8^+^ T-cells and anti-hCD19 CAR T-cells.**

a. CRISPR/Cas9 knockout of B4GALT1 in OT-1 CD8^+^ T-cells significantly increases in vitro specific killing activities on B16F10-OVA cells.

b. CRISPR/Cas9 knockout of CD8α in OT-1 CD8^+^ T-cells significantly reduces in vitro specific killing activities on B16F10-OVA cells.

c. CRISPR/Cas9 knockout of B4GALT1 in anti-hCD19 CAR T-cells does not affect in vitro killing of Nalm6 cells.

d. CRISPR/Cas9 knockout of CD8α in anti-hCD19 CAR T-cells does not affect in vitro killing of Nalm6 cells.

All of the p values were calculated by a two-tailed Student’s t test. Data are shown as the mean ± SEM. *P < 0.05; **P < 0.01; NS, not significant.

**Fig. S14. Lactose depletes surface Gal-1 on MC38 and OT-1 cells during coculture.**

a. Representative FACS plot for OT-1 cells cocultured with MC38 cells at a 2:1 ratio for 8 hours with MC38 cell conditioned medium.

b. MFIs of surface Gal-1 on MC38 cells were measured by FACS with PE-anti-Gal-1 antibody as indicated.

c. MFIs of surface Gal-1 on OT-1 cells were measured by FACS with PE-anti-Gal-1 antibody as indicated.

The p value was calculated by a two-tailed Student’s t test. Data are shown as the mean ± SEM. ***P < 0.001.

**Fig. S15. Lactose treatment does not affect TCR activation of OT-1 T-cells in the absence of exogenous or endogenous Gal-1.**

Lactose treatment has no significant effect on the mRNA expression of Tnfα and Ifnγ in OT-1 T-cells stimulated by anti-CD3/CD28 antibodies. The relative mRNA expression levels of Tnfα (a) and Ifnγ (b) were measured by quantitative RT‒qPCR. The p values were calculated by a two-tailed Student’s t test.

Data are shown as the mean ± SEM. NS, not significant.

**Fig. S16. Lactose reverses the inhibitory effect of MC38-transferred Gal-1 on TCR activation of OT-1 CD8^+^ T-cells.**

OT-1 T-cells cocultured with wild-type or Gal-1 knockout MC38 cells were sorted for anti-CD3/CD28 antibody stimulation in conditions with or without lactose. The relative mRNA expression levels of Tnfα(a) and Ifnγ(b) were measured by quantitative RT‒qPCR. The p values were calculated by a two-tailed Student’s t test.

Data are shown as the mean ± SEM. *P < 0.05; ***P < 0.001. NS, not significant.

**Fig. S17. Lactose treatment enhances TCR activation of tumor-infiltrated CD8^+^ T-cells in vitro.**

a. Lactose treatment depletes surface Gal-1 of MC38 tumor-infiltrated CD8^+^ T-cells. The MFIs of Gal-1 were measured by FACS.

b. Lactose treatment increases the expression of Tnfα and Ifnγ in MC38 tumor-infiltrated CD8^+^ T-cells stimulated in vitro by anti-CD3/CD28 antibodies. The mRNA expression levels of Tnfα and Ifnγ were measured by quantitative RT‒qPCR. The p values were calculated by a two-tailed Student’s t test.

Data are shown as the mean ± SEM. *P < 0.05; ***P < 0.001.

**Fig. S18. Dosage analysis of intravenous lactose administration to treat subcutaneous MC38 tumors in wild-type mice.**

**Lactose concentrations of** 50 mM (a), 250 mM (b), and 400 mM (c) in 200 μl PBS were intravenously injected every two days into wild-type mice subcutaneously inoculated with MC38 tumors. The p values were calculated by two-way ANOVA.

Data are shown as the mean ± SEM.

**Figure S19. Combinational treatment of MC38 tumors in wild-type mice.**

a-d. Combinational treatment of MC38 tumors in wild-type mice with lactose and anti-PD-1 (PDCD1) antibody.

e-f. Combinational treatment of MC38 tumors in wild-type mice with lactose and anti-PD-L1 (CD274) antibody.

The p values were calculated by two-way ANOVA. Data are shown as the mean ± SEM.

**Fig. S20. Properties of lactose and related derivates.**

a. Parallel artificial membrane permeability assay (PAMPA) of lactose. Testosterone and methotrexate were used as positive and negative controls, respectively.

b. Structures of lactose, sucrose, lacNAc, and lactose-BSA. SDS‒PAGE demonstrates efficient lactose coupling on BSA.

c. Competition curves of lactose, sucrose, lacNAc, and lactose-BSA on ECL, sWGA, Gal-1 binding on the MC38 cell surface.

d. The effect of intravenous lacNAc (250 μl of 20 mM in PBS) treatment on the growth of MC38 tumors in syngeneic C57BL/6J wild-type mice. The p value was calculated by two-way ANOVA.

Data are shown as the mean ± SEM.

**Fig. S21. Safety and toxicity profiles in wild-type C57BL/6J mice after short-term and long-term intravenous lactose injection.**

a. Short-term (24 hours after injection) effects of intravenous lactose injection on blood/serum biochemical and hematological parameters in wild-type mice. The p values were calculated by a two-tailed Student’s t test.

b. Long-term (24 days) effects of 12 times’ intravenous lactose injections on blood/serum biochemical and hematological parameters in wild-type mice. The p values were calculated by a two-tailed Student’s t test.

c. Body weight measurements of mice treated with intravenous lactose or PBS injection over 23 days. The p value was calculated by two-way ANOVA.

Data are shown as the mean ± SEM. *P < 0.05; NS, not significant.

ALT, alanine aminotransferase; AST, aspartate aminotransferase; CK, creatine kinase; LDH, lactate dehydrogenase; ALP, alkaline phosphatase; Pi, inorganic phosphorus; CR, creatinine; BUN, blood urea nitrogen; GLU, glucose; WBC, white blood cells; LYM, lymphocyte; MID, MID cells; GRA, granulocyte; RBC, red blood cells; HGB, hemoglobin; HCT, hematocrit; MCV, mean corpuscular volume; MCH, mean corpuscular hemoglobin; MCHC, mean corpuscular hemoglobin concentration; RDW-SD, red cell distribution width-standard deviation; RDW-CV, red cell distribution-coefficient of variation; PLT, platelets; PCT, procalcitonin; MPV, mean platelet volume; PDW, platelet distribution width.

**Figure S22. RNA-seq analysis of subcutaneous MC38 tumors in wild-type mice treated with control and lactose.**

a. Heatmap showing differentially expressed genes (DEGs) between control (PBS) and lactose-treated tumor samples. The genes in interferon γ (IFNγ) signaling are labeled on the left side.

b. Volcano plot showing upregulated and downregulated genes (p value< 0.05) in tumor samples treated with lactose. The genes in the interferon γ (IFNγ) signaling pathway are labeled with dark blue and dark red. The top genes and some genes in the IFNγ signaling pathway are annotated. The p value was calculated by the Wald test, and p.adjust was calculated by Benjamini‒Hochberg with the R package DESeq2 (version 1.22.2).

c. Volcano plot showing HALLMARKER gene sets significantly changed in tumor samples after intravenous lactose treatment. The annotated pathways represent the top downregulated and upregulated gene sets. The p value was calculated by the clusterProfiler (version 3.12.0) R package.

d-e. Transcriptional signature analysis of control and lactose-treated tumor samples. The p value was calculated by a two-tailed Student’s t test.

Data are shown as the mean ± SEM. **P < 0.01, ***P < 0.001.

**Fig. S23. Single-cell transcriptome analysis of infiltrated T-cells in MC38 tumors treated with control and lactose.**

a. Flow chart of single-cell RNA-seq and TCR analysis.

b. Control (6,607 cells) and lactose (7,737 cells) treated tumor samples were projected onto 9 T-cell clusters, respectively.

c. The distributions of each cluster in control- and lactose-treated tumor samples.

**Figure S24. Single-cell TCR sequencing analysis of infiltrated T-cells in MC38 tumors treated with control and lactose solutions.**

a. Clonal homeostatic space representations across control and lactose-treated samples using the CDR3 AA sequence for clonotype calling.

b. Relative proportional spade occupied by specific clonotypes across control and lactose-treated samples using the CDR3 AA sequence for clonotype calling.

c. Diversity measures based on clonotypes using Shannon, Inv Simpson, Chao, abundance-based coverage estimator (ACE), and Inv Pielou indices.

d. Morisita overlap quantifications for clonotypes across nine clusters.

e. UMAP projections of the largest T-cell clone, CN1, in control and lactose-treated tumor samples.

f. The distribution of the largest T-cell clone, CN1, in different T-cell clusters of control- and lactose-treated tumor samples.

**Figure S25. Exogenous expression of LALBA in MC38 cells leads to B4GALT1-dependent lactose synthesis.**

a. The amount of lactose in the cell pellets was quantified by LC‒MS. The p value was calculated by a two-tailed Student’s t test.

b. The lactose concentration in the culture medium was quantified by LC‒MS. The p value was calculated by a two-tailed Student’s t test.

c. The background of lactose measurement comes from the culture medium. Data are shown as the mean ± SEM. ***P < 0.001.

**Figure S26. Overexpression of LALBA in part of MC38 tumors reduces tumor growth in wild-type mice.**

MC38-LALBA cells were mixed with MC38-control cells at different ratios (1:1 and 1:9) and subcutaneously inoculated into wild-type mice. The p values were calculated between control samples and other samples by two-way ANOVA. Data are shown as the mean ± SEM. ***P < 0.001.

**Figure S27. The effect of intratumoral injection of lentivirus expressing the LALBA mutants D107A (a) and A126K (b) on the growth of MC38 tumors in syngeneic wild-type mice.**

Arrows indicate the time points for lentivirus injection. The p values were calculated by two-way ANOVA.

Data are shown as the mean ± SEM.

**Figure S28. Intravenous injection of lactose shows a similar effect as pembrolizumab, an anti-human PD-1 antibody, on the growth of human SGC7901 tumors in immunodeficient NPG mice reconstituted with the human immune system.**

The p value was calculated by two-way ANOVA. Data are shown as the mean ± SEM.

**Figure S29. Detailed information on ex vivo culture of patient-derived tumor fragments.**

a. Information on the 11 tumor samples used in the current study.

b. Comparisons of individual parameters assessed in untreated and lactose-treated patient-derived tumor fragments. The mRNA expression level was measured by quantitative RT‒qPCR and normalized by hGAPDH. The p values were calculated by a paired two-tailed Student’s t test.

c. Comparisons of individual parameters assessed in untreated and anti-PD-1-treated patient-derived tumor fragments. The mRNA expression level was measured by quantitative RT‒qPCR and normalized by hGAPDH. The p values were calculated by a paired two-tailed Student’s t test.

**Supplementary Table 1. List of sequences of gRNAs and primers used in current study.**

**Supplementary Table 2. List of plasmids used in current study.**

**Supplementary Table 3. Information of custom library used in current study.**

**Supplementary Table 4. Results of ex vivo CRISPR/Cas9 genome-wide screenings in Fig 1b.**

**Supplementary Table 5. Results of in vivo CRISPR/Cas9 screenings with custom small library Fig 1c.**

**Supplementary Table 6. Gene lists of signatures used in Fig S22.**

## References

1 Sung, H. et al. Global Cancer Statistics 2020: GLOBOCAN Estimates of Incidence and Mortality Worldwide for 36 Cancers in 185 Countries. CA Cancer J Clin 71, 209–249, doi:10.3322/caac.21660 (2021).

2 Esfahani, K. et al. A review of cancer immunotherapy: from the past, to the present, to the future. Curr Oncol 27, S87–S97, doi:10.3747/co.27.5223 (2020).

3 Kraehenbuehl, L., Weng, C.-H., Eghbali, S., Wolchok, J. D. & Merghoub, T. Enhancing immunotherapy in cancer by targeting emerging immunomodulatory pathways. Nature Reviews Clinical Oncology 19, 37–50 (2022).

4 Waldman, A. D., Fritz, J. M. & Lenardo, M. J. A guide to cancer immunotherapy: from T cell basic science to clinical practice. Nature Reviews Immunology 20, 651–668 (2020).

5 Miller, K. D. et al. Cancer treatment and survivorship statistics, 2022. CA Cancer J Clin 72, 409–436, doi:10.3322/caac.21731 (2022).

6 Spencer, K. R. et al. Biomarkers for Immunotherapy: Current Developments and Challenges. American Society of Clinical Oncology educational book. American Society of Clinical Oncology. Annual Meeting 35, e493–503, doi:10.14694/EDBK_160766

7 Havel, J. J., Chowell, D. & Chan, T. A. The evolving landscape of biomarkers for checkpoint inhibitor immunotherapy. Nature reviews. Cancer 19, 133–150, doi:10.1038/s41568-019-0116-x (2019).

8 Hodi, F. S. et al. Biologic activity of cytotoxic T lymphocyte-associated antigen 4 antibody blockade in previously vaccinated metastatic melanoma and ovarian carcinoma patients. Proceedings of the National Academy of Sciences of the United States of America 100, 4712–4717, doi:10.1073/pnas.0830997100 (2003).

9 Brahmer, J. R. et al. Safety and activity of anti-PD-L1 antibody in patients with advanced cancer. The New England journal of medicine 366, 2455–2465, doi:10.1056/NEJMoa1200694 (2012).

10 Topalian, S. L. et al. Safety, activity, and immune correlates of anti-PD-1 antibody in cancer. The New England journal of medicine 366, 2443–2454, doi:10.1056/NEJMoa1200690 (2012).

11 Ribas, A. & Wolchok, J. D. Cancer immunotherapy using checkpoint blockade. Science 359, 1350–1355 (2018).

12 Cercek, A. et al. PD-1 Blockade in Mismatch Repair-Deficient, Locally Advanced Rectal Cancer. The New England journal of medicine, doi:10.1056/NEJMoa2201445 (2022).

13 June, C. H., O’Connor, R. S., Kawalekar, O. U., Ghassemi, S. & Milone, M. C. CAR T cell immunotherapy for human cancer. Science 359, 1361–1365, doi:10.1126/science.aar6711 (2018).

14 Rafiq, S., Hackett, C. S. & Brentjens, R. J. Engineering strategies to overcome the current roadblocks in CAR T cell therapy. Nature reviews. Clinical oncology 17, 147–167, doi:10.1038/s41571-019-0297-y (2020).

15 Agata, Y. et al. Expression of the PD-1 antigen on the surface of stimulated mouse T and B lymphocytes. International immunology 8, 765–772, doi:10.1093/intimm/8.5.765 (1996).

16 Yu, Y. et al. Single-cell RNA-seq identifies a PD-1(hi) ILC progenitor and defines its development pathway. Nature 539, 102–106, doi:10.1038/nature20105 (2016).

17 Gordon, S. R. et al. PD-1 expression by tumour-associated macrophages inhibits phagocytosis and tumour immunity. Nature 545, 495–499, doi:10.1038/nature22396 (2017).

18 Park, B. V. et al. TGFbeta1-Mediated SMAD3 Enhances PD-1 Expression on Antigen-Specific T Cells in Cancer. Cancer Discov 6, 1366–1381, doi:10.1158/2159-8290.CD-15-1347 (2016).

19 Stephen, T. L. et al. SATB1 Expression Governs Epigenetic Repression of PD-1 in Tumor-Reactive T Cells. Immunity 46, 51–64, doi:10.1016/j.immuni.2016.12.015 (2017).

20 Okada, M. et al. Blockage of Core Fucosylation Reduces Cell-Surface Expression of PD-1 and Promotes Anti-tumor Immune Responses of T Cells. Cell Rep 20, 1017–1028, doi:10.1016/j.celrep.2017.07.027 (2017).

21 Meng, X. et al. FBXO38 mediates PD-1 ubiquitination and regulates anti-tumour immunity of T cells. Nature 564, 130–135, doi:10.1038/s41586-018-0756-0 (2018).

22 Zhou, X. A. et al. KLHL22 maintains PD-1 homeostasis and prevents excessive T cell suppression. Proceedings of the National Academy of Sciences of the United States of America 117, 28239–28250, doi:10.1073/pnas.2004570117 (2020).

23 Wei, J. et al. Targeting REGNASE-1 programs long-lived effector T cells for cancer therapy. Nature 576, 471–476, doi:10.1038/s41586-019-1821-z (2019).

24 Zhao, H. et al. Genome-wide fitness gene identification reveals Roquin as a potent suppressor of CD8 T cell expansion and anti-tumor immunity. Cell Rep 37, 110083, doi:10.1016/j.celrep.2021.110083 (2021).

25 Ma, X. et al. CD36-mediated ferroptosis dampens intratumoral CD8(+) T cell effector function and impairs their antitumor ability. Cell Metab 33, 1001–1012 e1005, doi:10.1016/j.cmet.2021.02.015 (2021).

26 Xu, S. et al. Uptake of oxidized lipids by the scavenger receptor CD36 promotes lipid peroxidation and dysfunction in CD8(+) T cells in tumors. Immunity 54, 1561–1577 e1567, doi:10.1016/j.immuni.2021.05.003 (2021).

27 Rodeheffer, C. & Shur, B. D. Targeted mutations in beta1,4-galactosyltransferase I reveal its multiple cellular functions. Biochim Biophys Acta 1573, 258–270, doi:10.1016/s0304-4165(02)00392-6 (2002).

28 Cheng, X., Wang, X., Han, Y. & Wu, Y. The expression and function of beta-1,4-galactosyltransferase-I in dendritic cells. Cell Immunol 266, 32–39, doi:10.1016/j.cellimm.2010.08.008 (2010).

29 Han, Y. et al. Expression of beta-1,4-galactosyltransferase-I affects cellular adhesion in human peripheral blood CD4+ T cells. Cell Immunol 262, 11–17, doi:10.1016/j.cellimm.2009.08.004 (2010).

30 Gomez-Henao, W. et al. Relevance of glycans in the interaction between T lymphocyte and the antigen presenting cell. Int Rev Immunol 40, 274–288, doi:10.1080/08830185.2020.1845331 (2021).

31 Rechavi, O., Goldstein, I. & Kloog, Y. Intercellular exchange of proteins: the immune cell habit of sharing. FEBS Lett 583, 1792–1799, doi:10.1016/j.febslet.2009.03.014 (2009).

32 Davis, D. M. Intercellular transfer of cell-surface proteins is common and can affect many stages of an immune response. Nat Rev Immunol 7, 238–243, doi:10.1038/nri2020 (2007).

33 Demotte, N. et al. Restoring the association of the T cell receptor with CD8 reverses anergy in human tumor-infiltrating lymphocytes. Immunity 28, 414–424, doi:10.1016/j.immuni.2008.01.011 (2008).

34 Smith, L. K. et al. Interleukin-10 Directly Inhibits CD8(+) T Cell Function by Enhancing N-Glycan Branching to Decrease Antigen Sensitivity. Immunity 48, 299–312 e295, doi:10.1016/j.immuni.2018.01.006 (2018).

35 Demetriou, M., Granovsky, M., Quaggin, S. & Dennis, J. W. Negative regulation of T-cell activation and autoimmunity by Mgat5 N-glycosylation. Nature 409, 733–739, doi:10.1038/35055582 (2001).

36 Blouin, C. M. et al. Glycosylation-Dependent IFN-gammaR Partitioning in Lipid and Actin Nanodomains Is Critical for JAK Activation. Cell 166, 920–934, doi:10.1016/j.cell.2016.07.003 (2016).

37 Gordon-Alonso, M., Hirsch, T., Wildmann, C. & van der Bruggen, P. Galectin-3 captures interferon-gamma in the tumor matrix reducing chemokine gradient production and T-cell tumor infiltration. Nat Commun 8, 793, doi:10.1038/s41467-017-00925-6 (2017).

38 Dings, R. P. M., Miller, M. C., Griffin, R. J. & Mayo, K. H. Galectins as Molecular Targets for Therapeutic Intervention. Int J Mol Sci 19, doi:10.3390/ijms19030905 (2018).

39 Sato, Y., Fu, Y., Liu, H., Lee, M. Y. & Shaw, M. H. Tumor-immune profiling of CT-26 and Colon 26 syngeneic mouse models reveals mechanism of anti-PD-1 response. BMC Cancer 21, 1222, doi:10.1186/s12885-021-08974-3 (2021).

40 Jin, Y., et al. Abstract A010: Development of OVA-expressing immunogenic syngeneic mouse tumor models: CT26-OVA and B16-OVA. Molecular Cancer Therapeutics 18, A010–A010 (2019).

41 Yao, S. et al. Preclinical PET imaging of HIP/PAP using 1’-(18)F-fluoroethyl-beta-D-lactose. Oncotarget 8, 75162–75173, doi:10.18632/oncotarget.20654 (2017).

42 Weser, E. & Sleisenger, M. H. Metabolism of circulating disaccharides in man and the rat. The Journal of clinical investigation 46, 499–505, doi:10.1172/JCI105552 (1967).

43 Boratyn ski, J. & Roy, R. High temperature conjugation of proteins with carbohydrates. 15, 131–138 (1998).

44 Tao, C., Chuah, Y. J., Xu, C. & Wang, D. A. Albumin conjugates and assemblies as versatile bio-functional additives and carriers for biomedical applications. Journal of materials chemistry. B 7, 357–367, doi:10.1039/c8tb02477d (2019).

45 Jiang, P. et al. Signatures of T cell dysfunction and exclusion predict cancer immunotherapy response. Nat Med 24, 1550–1558, doi:10.1038/s41591-018-0136-1 (2018).

46 Mathewson, N. D. et al. Inhibitory CD161 receptor identified in glioma-infiltrating T cells by single-cell analysis. Cell 184, 1281–1298 e1226, doi:10.1016/j.cell.2021.01.022 (2021).

47 Andreatta, M. et al. Interpretation of T cell states from single-cell transcriptomics data using reference atlases. Nature communications 12, 2965, doi:10.1038/s41467-021-23324-4 (2021).

48 Gottschalk, E. et al. Discovering tumor-reactive T-cell receptors through single-cell sequencing of tumor-infiltrating lymphocytes. bioRxiv, 2021.2011.2030.470597, doi:10.1101/2021.11.30.470597 (2021).

49 Malinovskii, V. A., Tian, J., Grobler, J. A. & Brew, K. Functional site in alpha-lactalbumin encompasses a region corresponding to a subsite in lysozyme and parts of two adjacent flexible substructures. Biochemistry 35, 9710–9715, doi:10.1021/bi960437c (1996).

50 Sadovnikova, A., Garcia, S. C. & Hovey, R. C. A Comparative Review of the Cell Biology, Biochemistry, and Genetics of Lactose Synthesis. J Mammary Gland Biol Neoplasia 26, 181–196, doi:10.1007/s10911-021-09490-7 (2021).

51 Voabil, P. et al. An ex vivo tumor fragment platform to dissect response to PD-1 blockade in cancer. Nat Med 27, 1250–1261, doi:10.1038/s41591-021-01398-3 (2021).

52 Gonzalez, H., Hagerling, C. & Werb, Z. Roles of the immune system in cancer: from tumor initiation to metastatic progression. Genes & development 32, 1267–1284, doi:10.1101/gad.314617.118 (2018).

53 Kalbasi, A. & Ribas, A. Tumour-intrinsic resistance to immune checkpoint blockade. Nature reviews. Immunology 20, 25–39, doi:10.1038/s41577-019-0218-4 (2020).

54 Jhunjhunwala, S., Hammer, C. & Delamarre, L. Antigen presentation in cancer: insights into tumour immunogenicity and immune evasion. Nature reviews. Cancer 21, 298–312, doi:10.1038/s41568-021-00339-z (2021).

55 Wellenstein, M. D. & de Visser, K. E. Cancer-Cell-Intrinsic Mechanisms Shaping the Tumor Immune Landscape. Immunity 48, 399–416, doi:10.1016/j.immuni.2018.03.004 (2018).

56 Anderson, N. M. & Simon, M. C. The tumor microenvironment. Curr Biol 30, R921–R925, doi:10.1016/j.cub.2020.06.081 (2020).

57 Arner, E. N. & Rathmell, J. C. Metabolic programming and immune suppression in the tumor microenvironment. Cancer cell 41, 421–433, doi:10.1016/j.ccell.2023.01.009 (2023).

58 Mendez-Huergo, S. P., Blidner, A. G. & Rabinovich, G. A. Galectins: emerging regulatory checkpoints linking tumor immunity and angiogenesis. Current opinion in immunology 45, 8–15, doi:10.1016/j.coi.2016.12.003 (2017).

59 Griffioen, A. W. & Thijssen, V. L. Galectins in tumor angiogenesis. Annals of translational medicine 2, 90, doi:10.3978/j.issn.2305-5839.2014.09.01 (2014).

60 Hamieh, M. et al. CAR T cell trogocytosis and cooperative killing regulate tumour antigen escape. Nature 568, 112–116, doi:10.1038/s41586-019-1054-1 (2019).

61 Poggio, M. et al. Suppression of Exosomal PD-L1 Induces Systemic Anti-tumor Immunity and Memory. Cell 177, 414–427 e413, doi:10.1016/j.cell.2019.02.016 (2019).

62 Platt, R. J. et al. CRISPR-Cas9 knockin mice for genome editing and cancer modeling. Cell 159, 440–455, doi:10.1016/j.cell.2014.09.014 (2014).

63 Kurachi, M. et al. Optimized retroviral transduction of mouse T cells for in vivo assessment of gene function. Nature protocols 12, 1980–1998, doi:10.1038/nprot.2017.083 (2017).

64 Robbins, P. F. et al. Single and dual amino acid substitutions in TCR CDRs can enhance antigen-specific T cell functions. J Immunol 180, 6116–6131, doi:10.4049/jimmunol.180.9.6116 (2008).

65 Martin, M. Cutadapt removes adapter sequences from high-throughput sequencing reads. *EMBnet*. journal 17, 10–12 (2011).

66 Magoc, T. & Salzberg, S. L. FLASH: fast length adjustment of short reads to improve genome assemblies. Bioinformatics 27, 2957–2963, doi:10.1093/bioinformatics/btr507 (2011).

67 Li, W. et al. MAGeCK enables robust identification of essential genes from genome-scale CRISPR/Cas9 knockout screens. Genome biology 15, 554, doi:10.1186/s13059-014-0554-4 (2014).

68 Yu, G., Wang, L.-G., Han, Y. & He, Q.-Y. clusterProfiler: an R package for comparing biological themes among gene clusters. Omics: a journal of integrative biology 16, 284–287 (2012).

69 Prato, C. A., Carabelli, J., Cattaneo, V., Campetella, O. & Tribulatti, M. V. Purification of recombinant galectins expressed in bacteria. STAR protocols 1, 100204 (2020).

70 Zhu, Y., et al. DEqMS: A Method for Accurate Variance Estimation in Differential Protein Expression Analysis. Molecular & cellular proteomics : MCP 19, 1047–1057, doi:10.1074/mcp.TIR119.001646 (2020).

71 Wu, T. et al. clusterProfiler 4.0: A universal enrichment tool for interpreting omics data. Innovation (Camb) 2, 100141, doi:10.1016/j.xinn.2021.100141 (2021).

72 Putri, G. H., Anders, S., Pyl, P. T., Pimanda, J. E. & Zanini, F. Analysing high-throughput sequencing data in Python with HTSeq 2.0. Bioinformatics 38, 2943–2945, doi:10.1093/bioinformatics/btac166 (2022).

73 Love, M. I., Huber, W. & Anders, S. Moderated estimation of fold change and dispersion for RNA-seq data with DESeq2. Genome biology 15, 550, doi:10.1186/s13059-014-0550-8 (2014).

