## Supplemental figres 1-29 for "Lactose blocks intercellular spreading of Galectin-1 from cancer cells to T-cells and activates tumor immunological control"

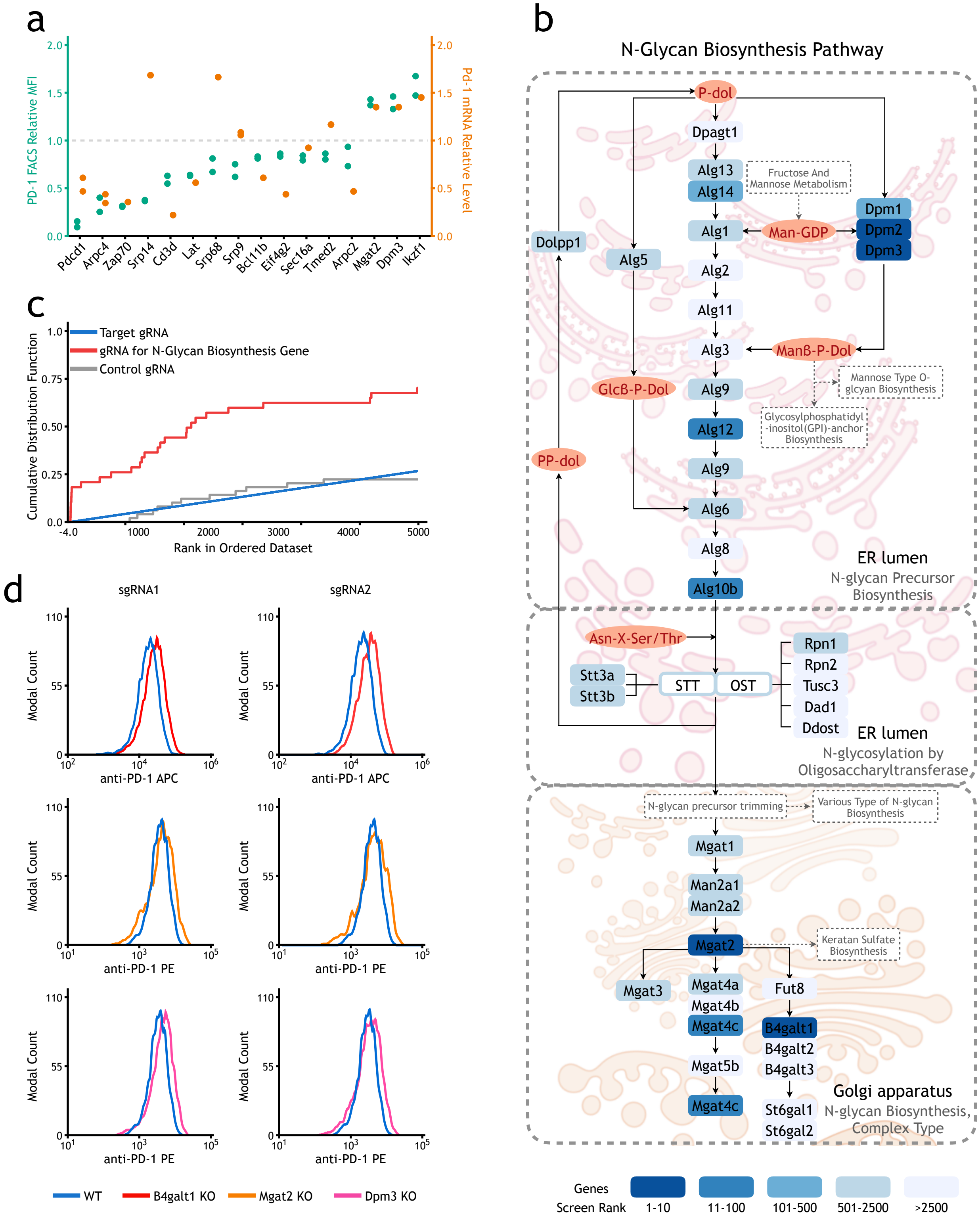
a
PD-1 FACS Relative MFI
Pd-1 mRNA Relative Level
Pdcd1
Arpc4
Zap70
Srp14
Cd3d
Lat
Srp68
Srp9
Bcl11b
Eif4g2
Sec16a
Tmed2
Arpc2
Mgat2
Dpm3
Ikzf1
b
N-Glycan Biosynthesis Pathway
P-dol
Dpagt1
Alg13
Alg14
Fructose And Mannose Metabolism
Dpm1
Dpm2
Dpm3
Man-GDP
Alg1
Dolpp1
Alg5
Alg2
Alg11
Alg3
ManB-P-Dol
Mannose Type O-glcyan Biosynthesis
GlcB-P-Dol
Alg9
Glycosylphosphatidyl-inositol(GPI)-anchor Biosynthesis
Alg12
PP-dol
Alg6
Alg8
Alg10b
ER lumen
N-glycan Precursor Biosynthesis
Asn-X-Ser/Thr
Stt3a
Stt3b
STT
OST
Rpn1
Rpn2
Tusc3
Dad1
Ddost
N-glycosylation by Oligosaccharyltransferase
N-glycan precursor trimming
Various Type of N-glycan Biosynthesis
Mgat1
Man2a1
Man2a2
Mgat2
Keratan Sulfate Biosynthesis
Mgat3
Mgat4a
Mgat4b
Mgat4c
Fut8
B4galt1
B4galt2
B4galt3
Mgat5b
St6gal1
St6gal2
Golgi apparatus
N-glycan Biosynthesis, Complex Type
Genes
Screen Rank
1-10
11-100
101-500
501-2500
>2500
c
Target gRNA
gRNA for N-Glycan Biosynthesis Gene
Control gRNA
Cumulative Distribution Function
Rank in Ordered Dataset
d
sgRNA1
sgRNA2
Modal Count
anti-PD-1 APC
anti-PD-1 PE
WT
B4galt1 KO
Mgat2 KO
Dpm3 KO

Figure S1

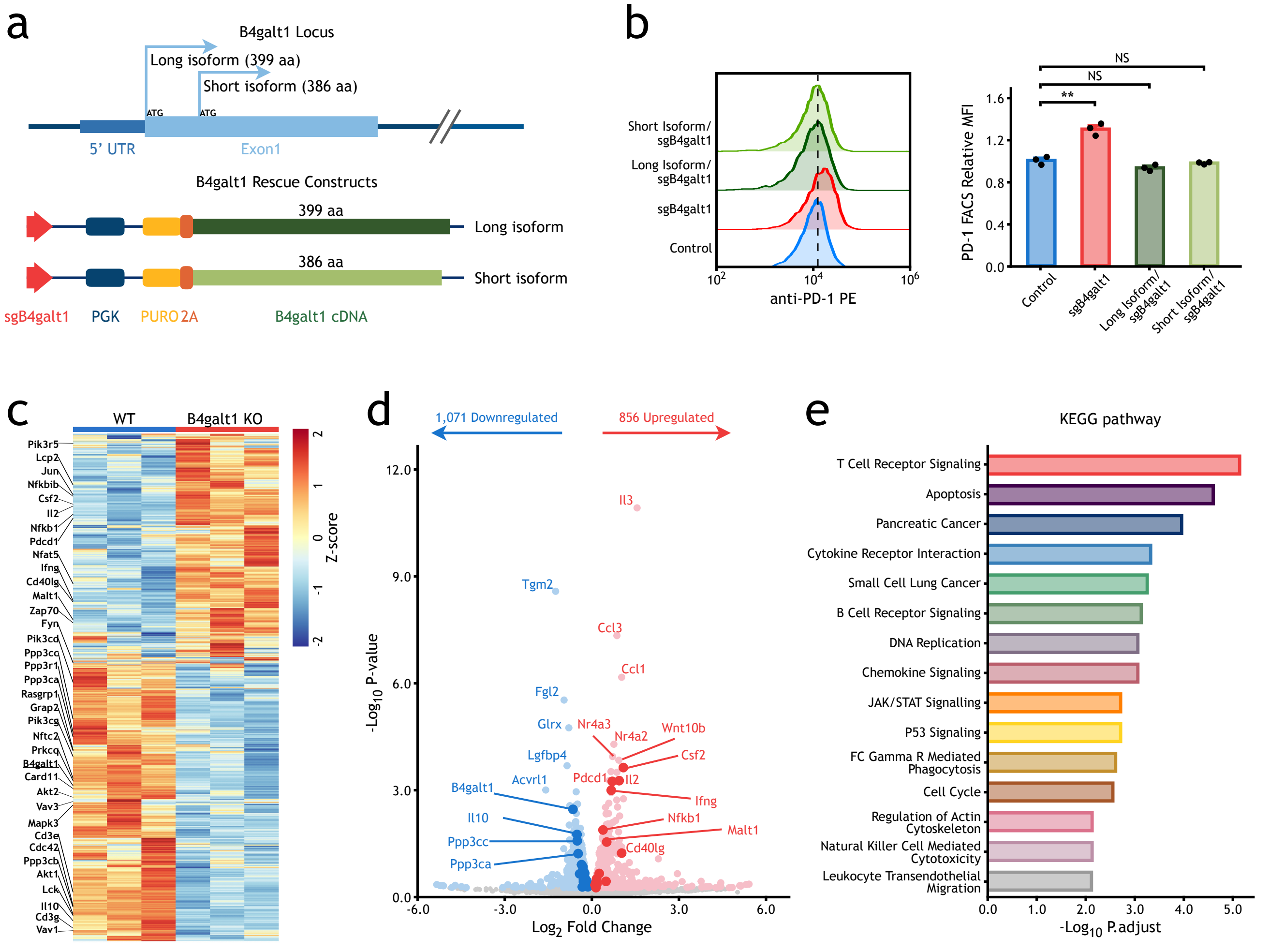

a
B4galt1 Locus
Long isoform (399 aa)
Short isoform (386 aa)
ATG
ATG
5' UTR
Exon1
B4galt1 Rescue Constructs
399 aa
Long isoform
386 aa
Short isoform
sgB4galt1
PGK
PURO
2A
B4galt1 cDNA
b
Short Isoform/ sgB4galt1
Long Isoform/ sgB4galt1
sgB4galt1
Control
anti-PD-1 PE
PD-1 FACS Relative MFI
NS
NS
**
Control
sgB4galt1
Long Isoform/ sgB4galt1
Short Isoform/ sgB4galt1
c
WT
B4galt1 KO
Z-score
Pik3r5
Lcp2
Jun
Nfkbib
Csf2
Il2
Nfkb1
Pdcd1
Nfat5
Ifng
Cd40lg
Malt1
Zap70
Fyn
Pik3cd
Ppp3cc
Ppp3r1
Ppp3ca
Rasgrp1
Grap2
Pik3cg
Nftc2
Prkcq
B4galt1
Card11
Akt2
Vav3
Mapk3
Cd3e
Cdc42
Ppp3cb
Akt1
Lck
Il10
Cd3g
Vav1
d
1,071 Downregulated
856 Upregulated
-Log10 P-value
Log2 Fold Change
Il3
Tgm2
Ccl3
Ccl1
Fgl2
Glrx
Nr4a3
Nr4a2
Wnt10b
Lgfbp4
Csf2
Acvrl1
Pdcd1
Il2
B4galt1
Ifng
Il10
Nfkb1
Malt1
Ppp3cc
Cd40lg
Ppp3ca
e
KEGG pathway
T Cell Receptor Signaling
Apoptosis
Pancreatic Cancer
Cytokine Receptor Interaction
Small Cell Lung Cancer
B Cell Receptor Signaling
DNA Replication
Chemokine Signaling
JAK/STAT Signalling
P53 Signaling
FC Gamma R Mediated Phagocytosis
Cell Cycle
Regulation of Actin Cytoskeleton
Natural Killer Cell Mediated Cytotoxicity
Leukocyte Transendothelial Migration
-Log10 P.adjust


Figure S2

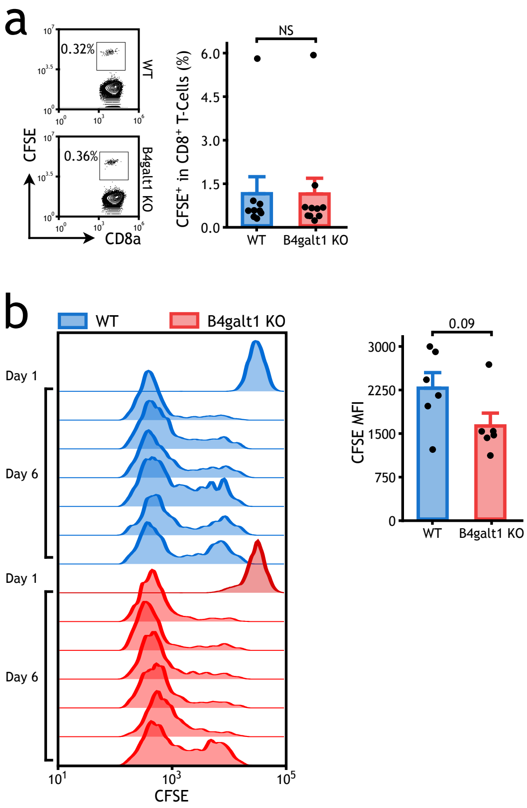
a
0.32%
WT
0.36%
B4galt1 KO
CFSE
CD8a
NS
CFSE+ in CD8+ T-Cells (%)
6.0
4.5
3.0
1.5
0.0
WT
B4galt1 KO
b
WT
B4galt1 KO
Day 1
Day 6
Day 1
Day 6
CFSE
0.09
3000
2250
1500
750
0
CFSE MFI
WT
B4galt1 KO

### Figure S3

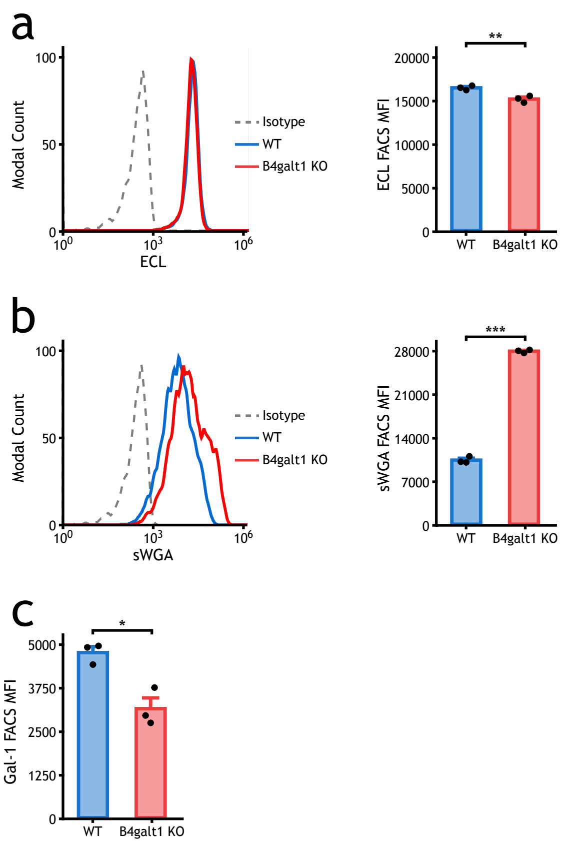
a
Modal Count
100
50
0
$10^0$
$10^3$
$10^6$
ECL
Isotype
WT
B4galt1 KO
**
ECL FACS MFI
20000
15000
10000
5000
0
WT
B4galt1 KO
b
Modal Count
sWGA
***
sWGA FACS MFI
28000
21000
14000
7000
c
*
Gal-1 FACS MFI
5000
3750
2500
1250

Figure S4

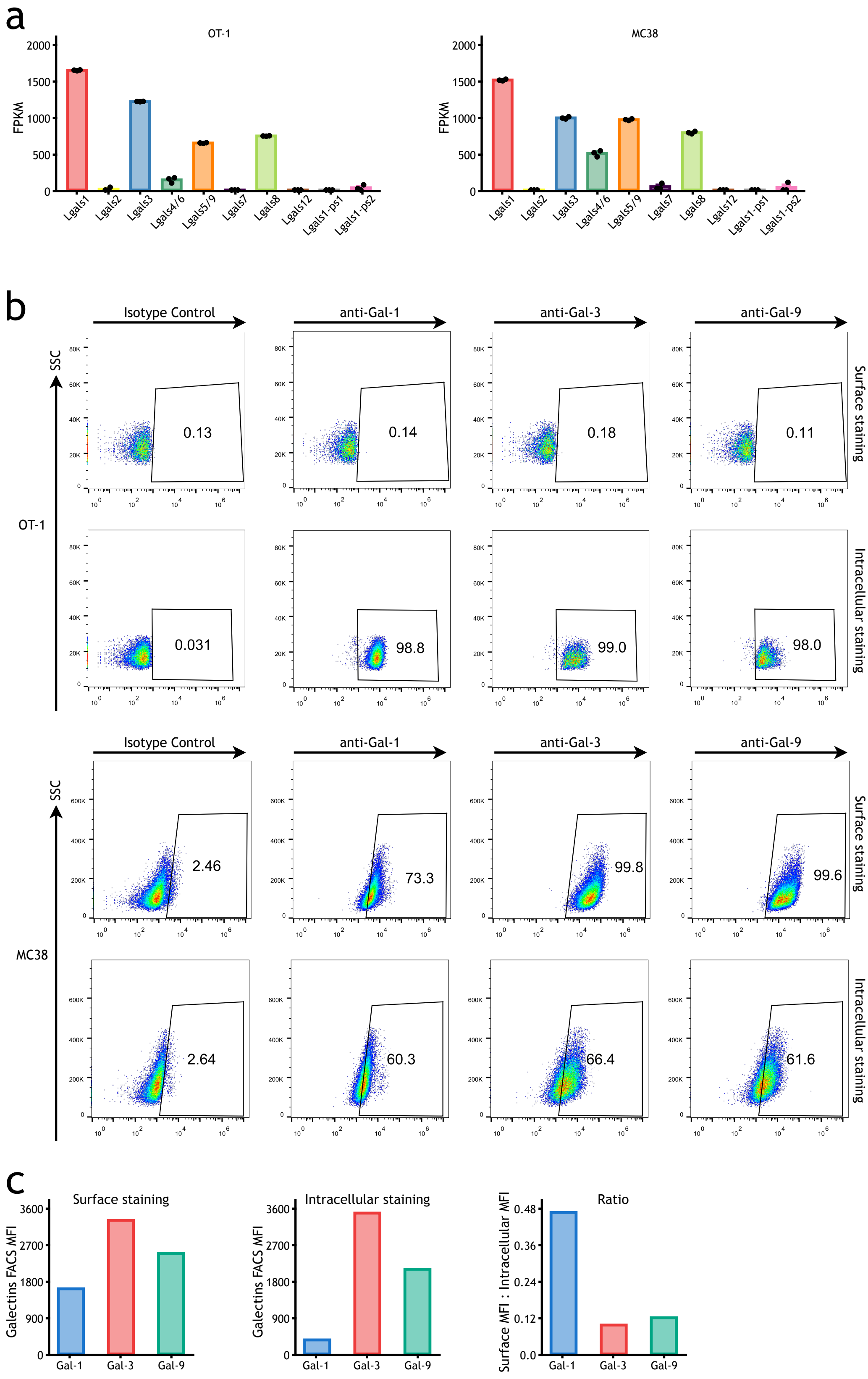
a
OT-1
MC38
FPKM
Lgals1
Lgals2
Lgals3
Lgals4/6
Lgals5/9
Lgals7
Lgals8
Lgals12
Lgals1-ps1
Lgals1-ps2
b
Isotype Control
anti-Gal-1
anti-Gal-3
anti-Gal-9
SSC
OT-1
MC38
Surface staining
Intracellular staining
0.13
0.14
0.18
0.11
0.031
98.8
99.0
98.0
2.46
73.3
99.8
99.6
2.64
60.3
66.4
61.6
c
Surface staining
Intracellular staining
Ratio
Galectins FACS MFI
Surface MFI : Intracellular MFI
Gal-1
Gal-3
Gal-9

Figure S5

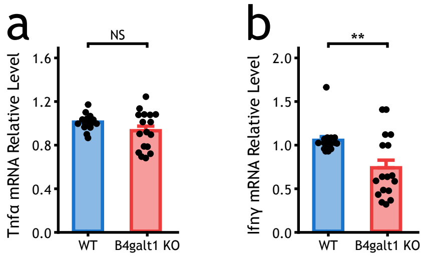

a
NS
Tnfα mRNA Relative Level
1.6
1.2
0.8
0.4
0.0
WT
B4galt1 KO
b
**
Ifnγ mRNA Relative Level
2.0
1.5
1.0
0.5
0.0
WT
B4galt1 KO


### Figure S6

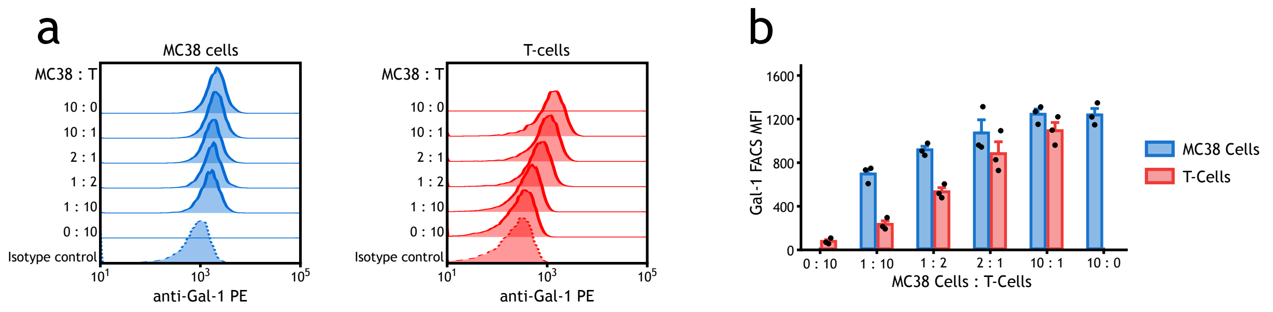
a
MC38 cells
T-cells
MC38 : T
10 : 0
10 : 1
2 : 1
1 : 2
1 : 10
0 : 10
Isotype control
anti-Gal-1 PE
b
Gal-1 FACS MFI
MC38 Cells : T-Cells
0 : 10
1 : 10
1 : 2
2 : 1
10 : 1
10 : 0
MC38 Cells
T-Cells

Figure S7

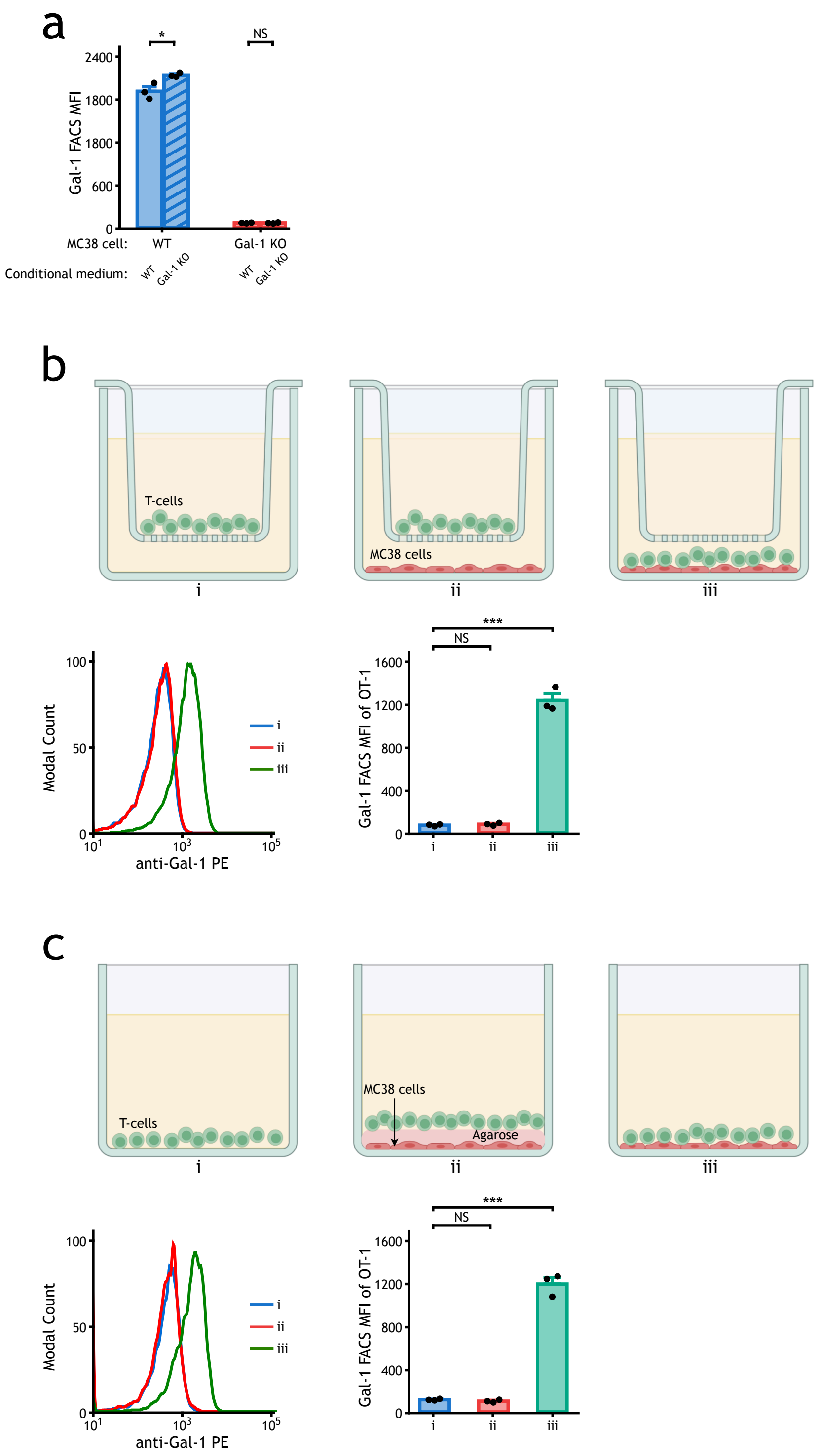
a
*
NS
2400
1800
1800
600
0
Gal-1 FACS MFI
MC38 cell:
WT
Gal-1 KO
Conditional medium:
WT
Gal-1 KO
WT
Gal-1 KO
b
T-cells
MC38 cells
i
ii
iii
100
50
0
Modal Count
$10^1$
$10^3$
$10^5$
anti-Gal-1 PE
i
ii
iii
***
NS
1600
1200
800
400
0
Gal-1 FACS MFI of OT-1
i
ii
iii
c
T-cells
MC38 cells
Agarose
i
ii
iii
100
50
0
Modal Count
$10^1$
$10^3$
$10^5$
anti-Gal-1 PE
i
ii
iii
***
NS
1600
1200
800
400
0
Gal-1 FACS MFI of OT-1
i
ii
iii

Figure S8

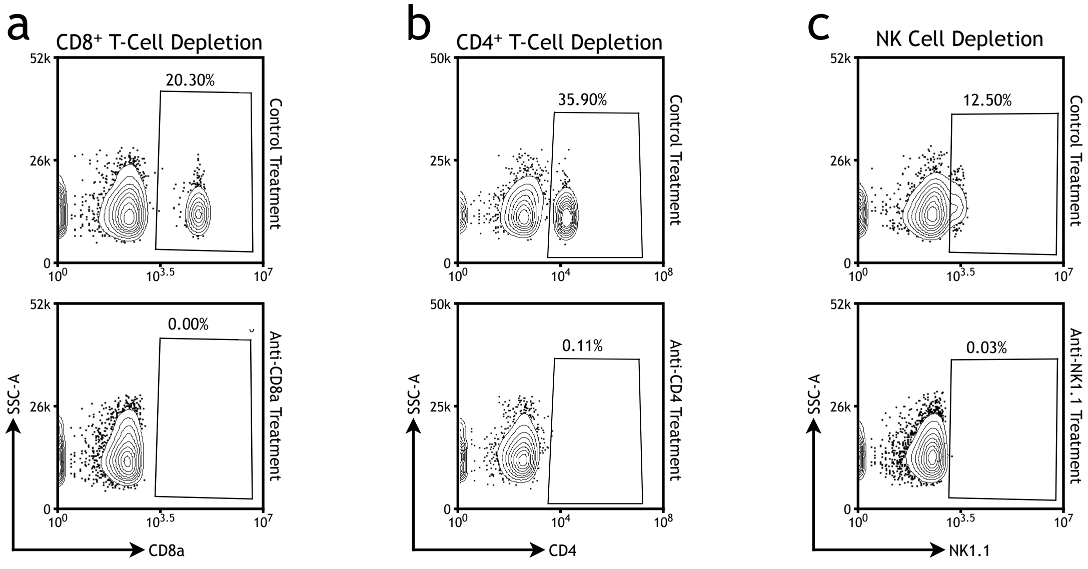
a
CD8+ T-Cell Depletion
b
CD4+ T-Cell Depletion
c
NK Cell Depletion
Control Treatment
Anti-CD8a Treatment
Anti-CD4 Treatment
Anti-NK1.1 Treatment
20.30%
0.00%
35.90%
0.11%
12.50%
0.03%
SSC-A
CD8a
CD4
NK1.1
52k
26k
50k
25k
0
$10^0$
$10^{3.5}$
$10^7$
$10^4$
$10^8$

Figure S9

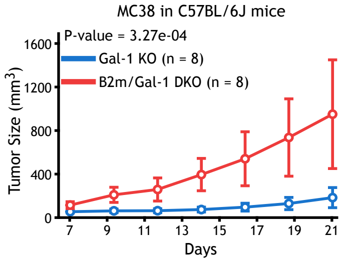
MC38 in C57BL/6J mice
P-value = 3.27e-04
Gal-1 KO (n = 8)
B2m/Gal-1 DKO (n = 8)
Tumor Size (mm$^3$)
1600
1200
800
400
0
7
9
11
13
15
17
19
21
Days

### Figure S10

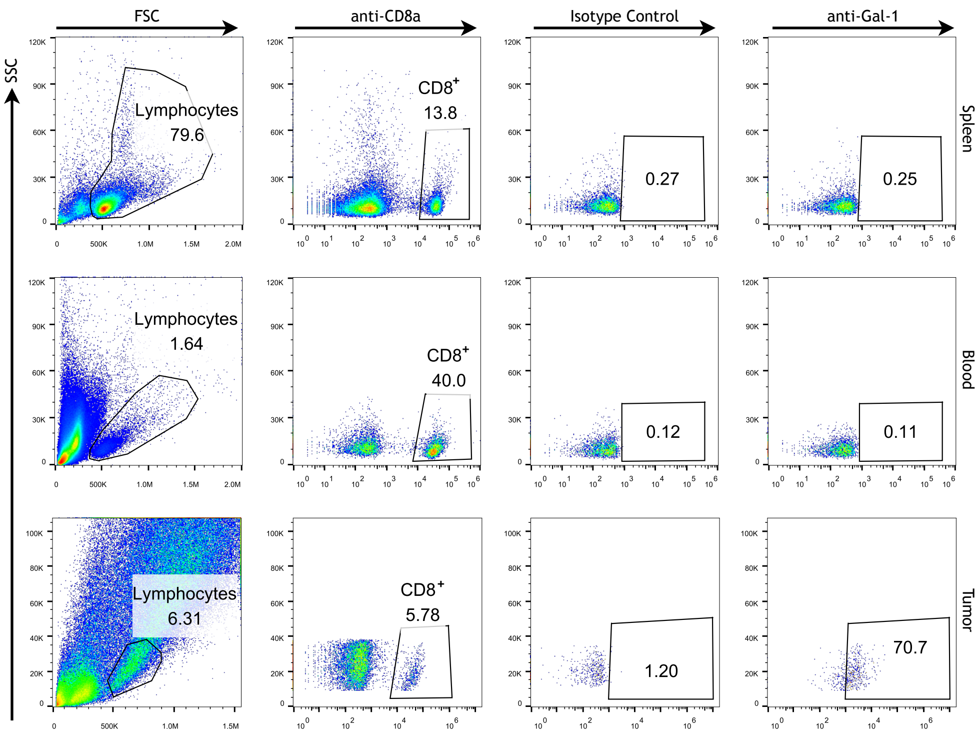
FSC
anti-CD8a
Isotype Control
anti-Gal-1
SSC
Spleen
Blood
Tumor
Lymphocytes
79.6
CD8+
13.8
0.27
0.25
Lymphocytes
1.64
CD8+
40.0
0.12
0.11
Lymphocytes
6.31
CD8+
5.78
1.20
70.7

### Figure S11

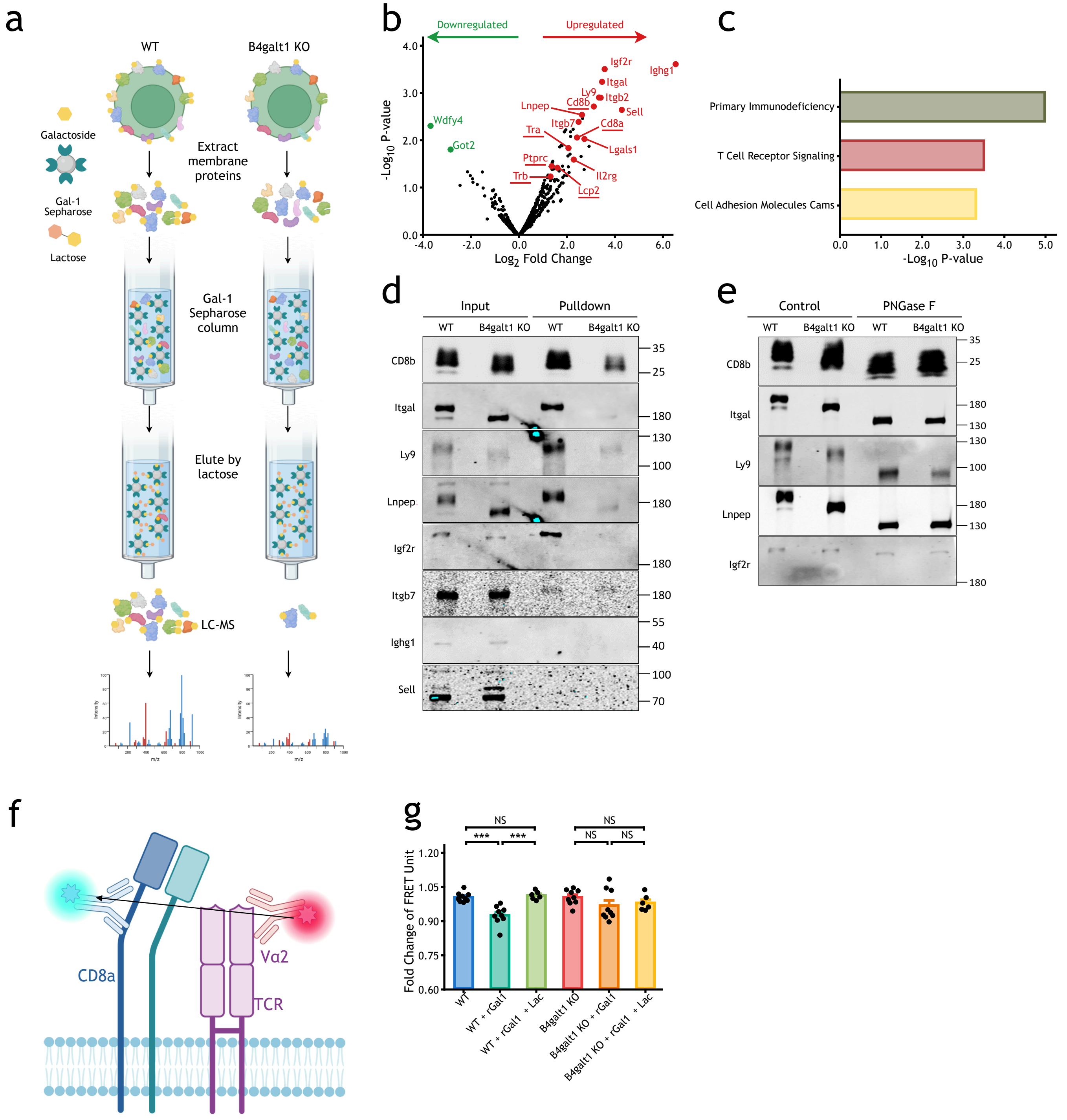
a
WT
B4galt1 KO
Galactoside
Gal-1 Sepharose
Lactose
Extract membrane proteins
Gal-1 Sepharose column
Elute by lactose
LC-MS
b
Downregulated
Upregulated
-Log10 P-value
Log2 Fold Change
Igf2r
Ighg1
Itgal
Ly9
Itgb2
Cd8b
Lnpep
Sell
Itgb7
Cd8a
Tra
Lgals1
Wdfy4
Got2
Ptprc
Trb
Il2rg
Lcp2
c
Primary Immunodeficiency
T Cell Receptor Signaling
Cell Adhesion Molecules Cams
-Log10 P-value
d
Input
Pulldown
WT
B4galt1 KO
CD8b
Itgal
Ly9
Lnpep
Igf2r
Itgb7
Ighg1
Sell
e
Control
PNGase F
f
CD8a
Vα2
TCR
g
Fold Change of FRET Unit
NS
***
WT
WT + rGal1
WT + rGal1 + Lac
B4galt1 KO
B4galt1 KO + rGal1
B4galt1 KO + rGal1 + Lac

Figure S12

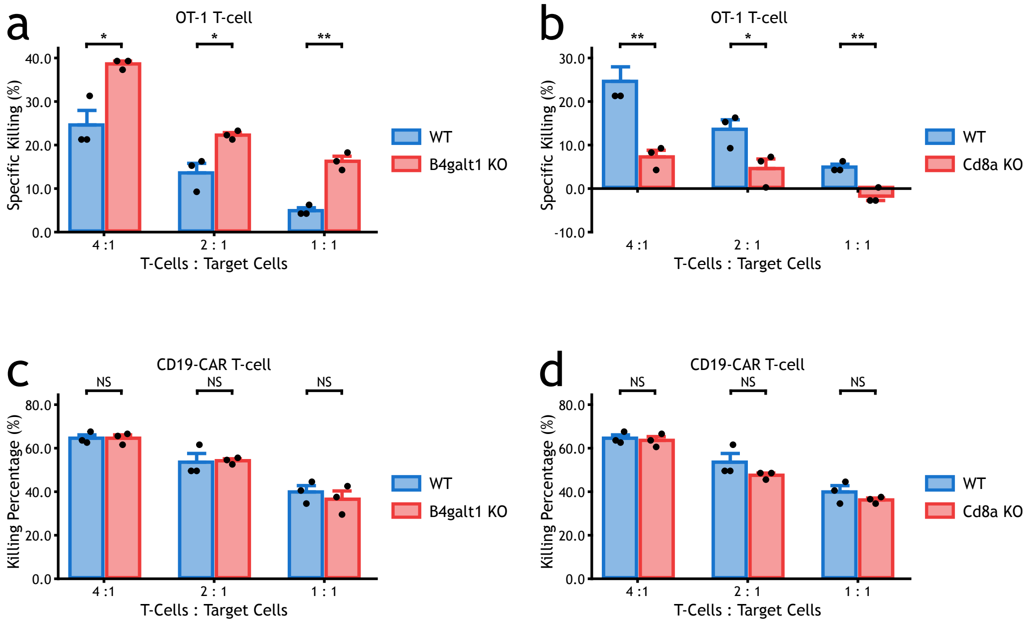
a
OT-1 T-cell
Specific Killing (%)
T-Cells : Target Cells
WT
B4galt1 KO
b
OT-1 T-cell
Specific Killing (%)
T-Cells : Target Cells
WT
Cd8a KO
c
CD19-CAR T-cell
Killing Percentage (%)
T-Cells : Target Cells
WT
B4galt1 KO
d
CD19-CAR T-cell
Killing Percentage (%)
T-Cells : Target Cells
WT
Cd8a KO

Figure S13

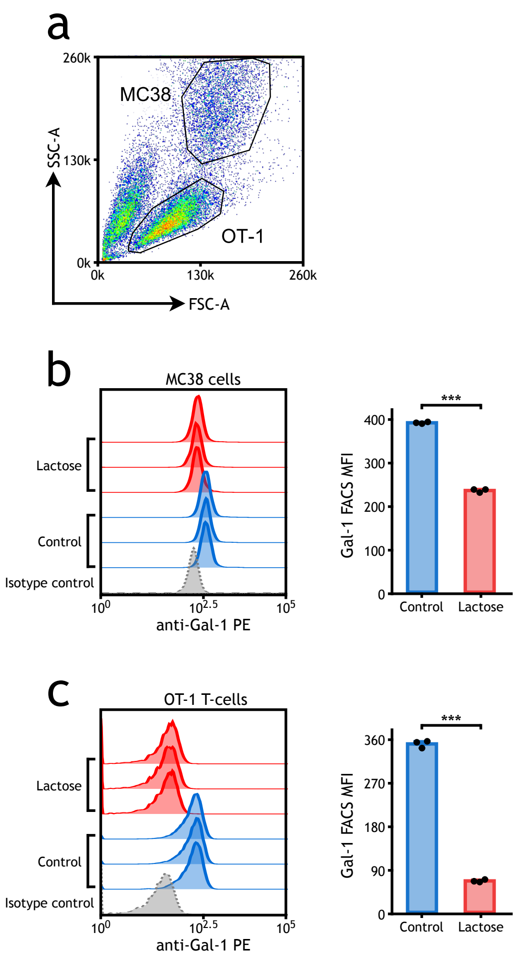
a
260k
130k
0k
0k
130k
260k
SSC-A
FSC-A
MC38
OT-1
b
MC38 cells
Lactose
Control
Isotype control
anti-Gal-1 PE
Gal-1 FACS MFI
400
300
200
100
0
***
Control
Lactose
c
OT-1 T-cells
Lactose
Control
Isotype control
anti-Gal-1 PE
Gal-1 FACS MFI
360
270
180
90
0
***
Control
Lactose

Figure S14

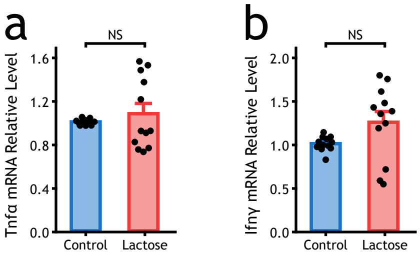
a
NS
Tnfα mRNA Relative Level
1.6
1.2
0.8
0.4
0.0
Control
Lactose
b
NS
Ifnγ mRNA Relative Level
2.0
1.5
1.0
0.5
0.0
Control
Lactose

### Figure S15

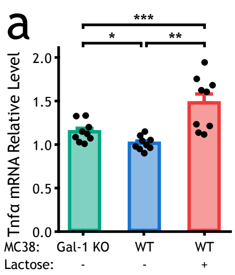
a
***
*
**
Tnfα mRNA Relative Level
2.0
1.5
1.0
0.5
0.0
MC38: Gal-1 KO WT WT
Lactose: - - +

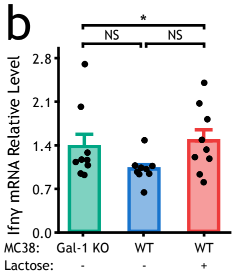
b
*
NS
NS
Ifnγ mRNA Relative Level
2.8
2.1
1.4
0.7
0.0
MC38: Gal-1 KO WT WT
Lactose: - - +

### Figure S16

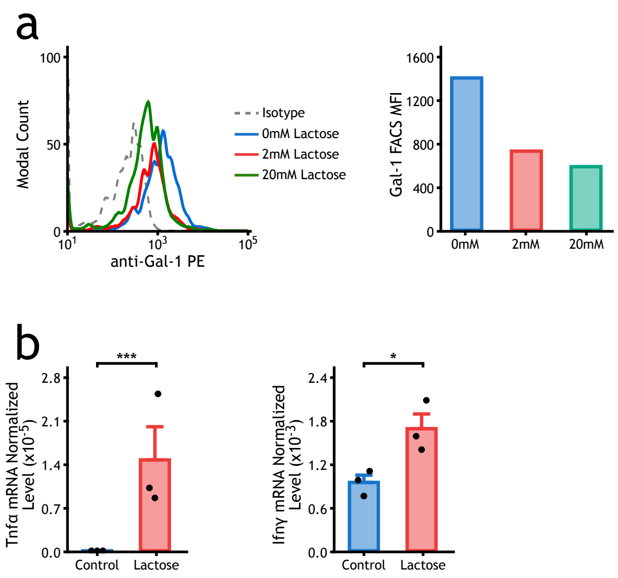
a
100
50
0
Modal Count
$10^1$
$10^3$
$10^5$
anti-Gal-1 PE
Isotype
0mM Lactose
2mM Lactose
20mM Lactose
1600
1200
800
400
0
Gal-1 FACS MFI
0mM
2mM
20mM
b
***
2.8
2.1
1.4
0.7
0.0
Tnfα mRNA Normalized Level (x10$^{-5}$)
Control
Lactose
*
2.4
1.8
1.2
0.6
0.0
Ifnγ mRNA Normalized Level (x10$^{-3}$)
Control
Lactose

### Figure S17

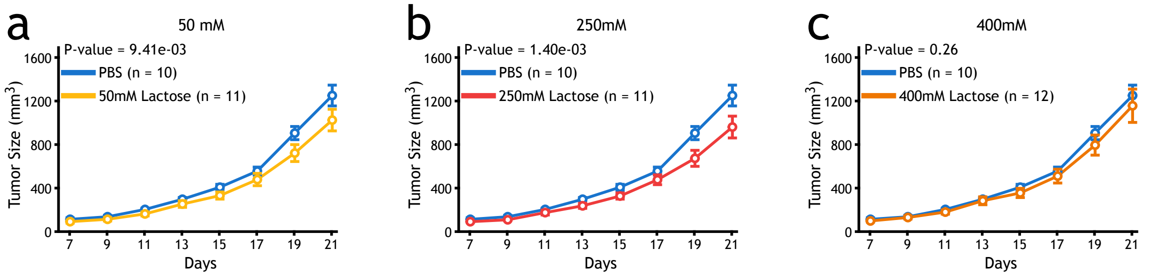
a
50 mM
P-value = 9.41e-03
PBS (n = 10)
50mM Lactose (n = 11)
Tumor Size (mm3)
1600
1200
800
400
0
7
9
11
13
15
17
19
21
Days
b
250mM
P-value = 1.40e-03
PBS (n = 10)
250mM Lactose (n = 11)
Tumor Size (mm3)
Days
c
400mM
P-value = 0.26
PBS (n = 10)
400mM Lactose (n = 12)
Tumor Size (mm3)
Days

Figure S18

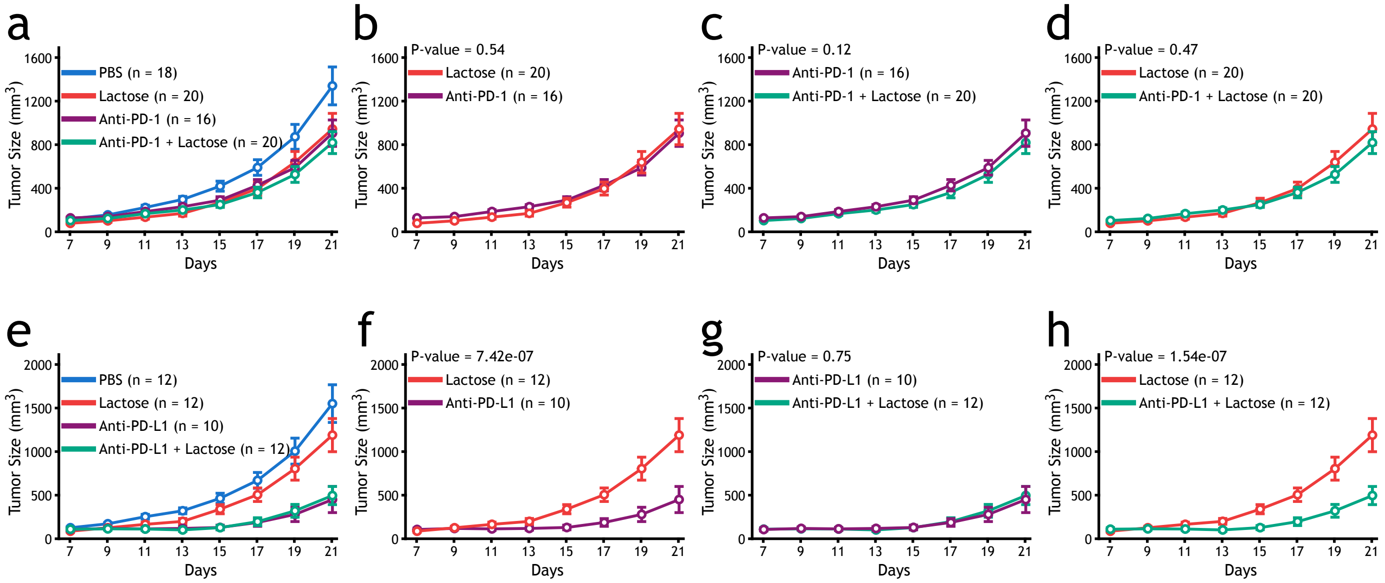
a
PBS (n = 18)
Lactose (n = 20)
Anti-PD-1 (n = 16)
Anti-PD-1 + Lactose (n = 20)
Tumor Size (mm³)
Days
b
P-value = 0.54
Lactose (n = 20)
Anti-PD-1 (n = 16)
c
P-value = 0.12
Anti-PD-1 (n = 16)
Anti-PD-1 + Lactose (n = 20)
d
P-value = 0.47
Lactose (n = 20)
Anti-PD-1 + Lactose (n = 20)
e
PBS (n = 12)
Lactose (n = 12)
Anti-PD-L1 (n = 10)
Anti-PD-L1 + Lactose (n = 12)
f
P-value = 7.42e-07
Lactose (n = 12)
Anti-PD-L1 (n = 10)
g
P-value = 0.75
Anti-PD-L1 (n = 10)
Anti-PD-L1 + Lactose (n = 12)
h
P-value = 1.54e-07
Lactose (n = 12)
Anti-PD-L1 + Lactose (n = 12)

Figure S19

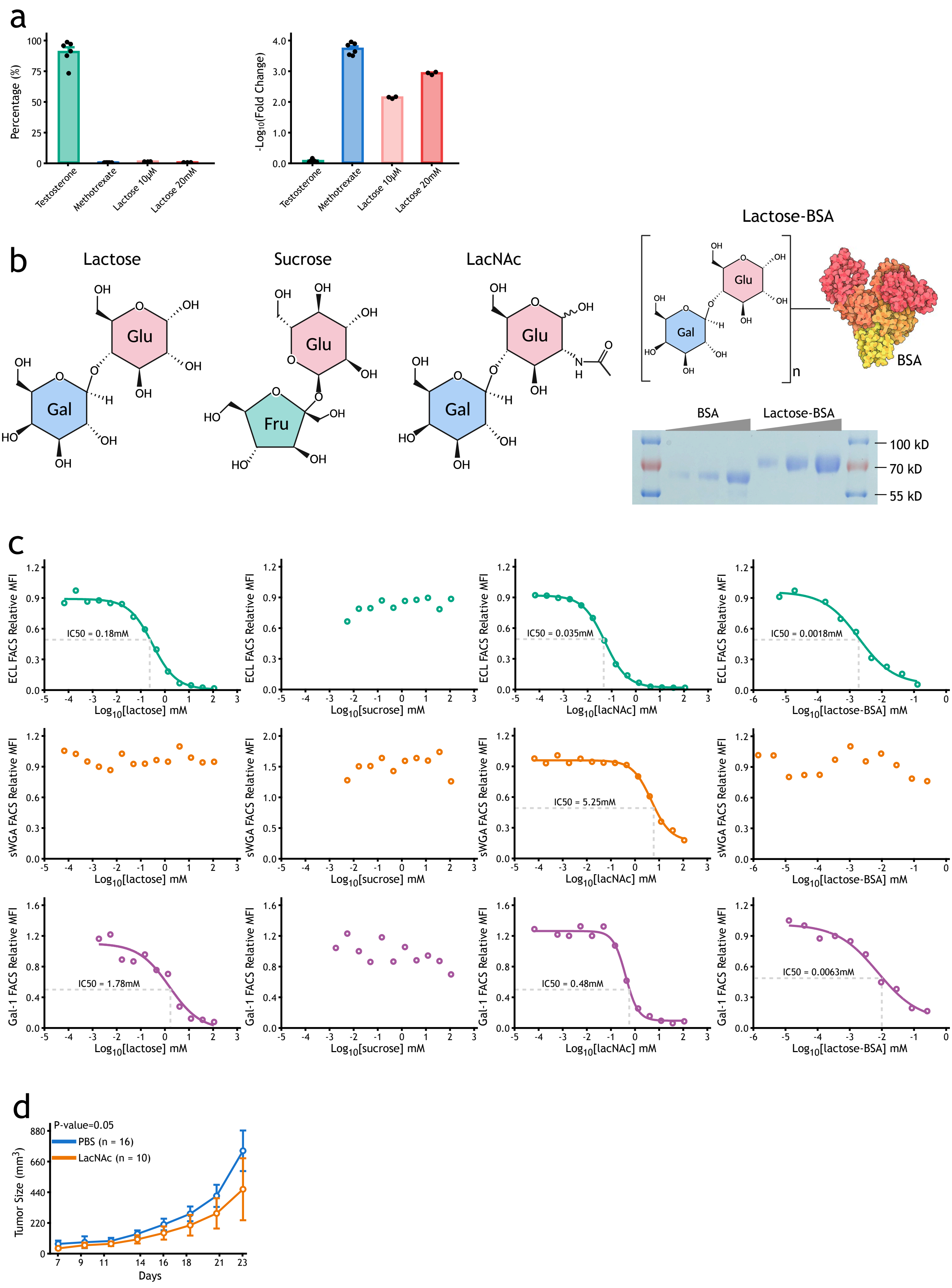
a
Percentage (%)
Testosterone
Methotrexate
Lactose 10μM
Lactose 20mM
-Log10(Fold Change)
b
Lactose
Sucrose
LacNAc
Lactose-BSA
Glu
Gal
Fru
BSA
100 kD
70 kD
55 kD
c
ECL FACS Relative MFI
sWGA FACS Relative MFI
Gal-1 FACS Relative MFI
IC50 = 0.18mM
IC50 = 0.035mM
IC50 = 0.0018mM
IC50 = 5.25mM
IC50 = 1.78mM
IC50 = 0.48mM
IC50 = 0.0063mM
Log10[lactose] mM
Log10[sucrose] mM
Log10[lacNAc] mM
Log10[lactose-BSA] mM
d
P-value=0.05
PBS (n = 16)
LacNAc (n = 10)
Tumor Size (mm3)
Days

### Figure S20

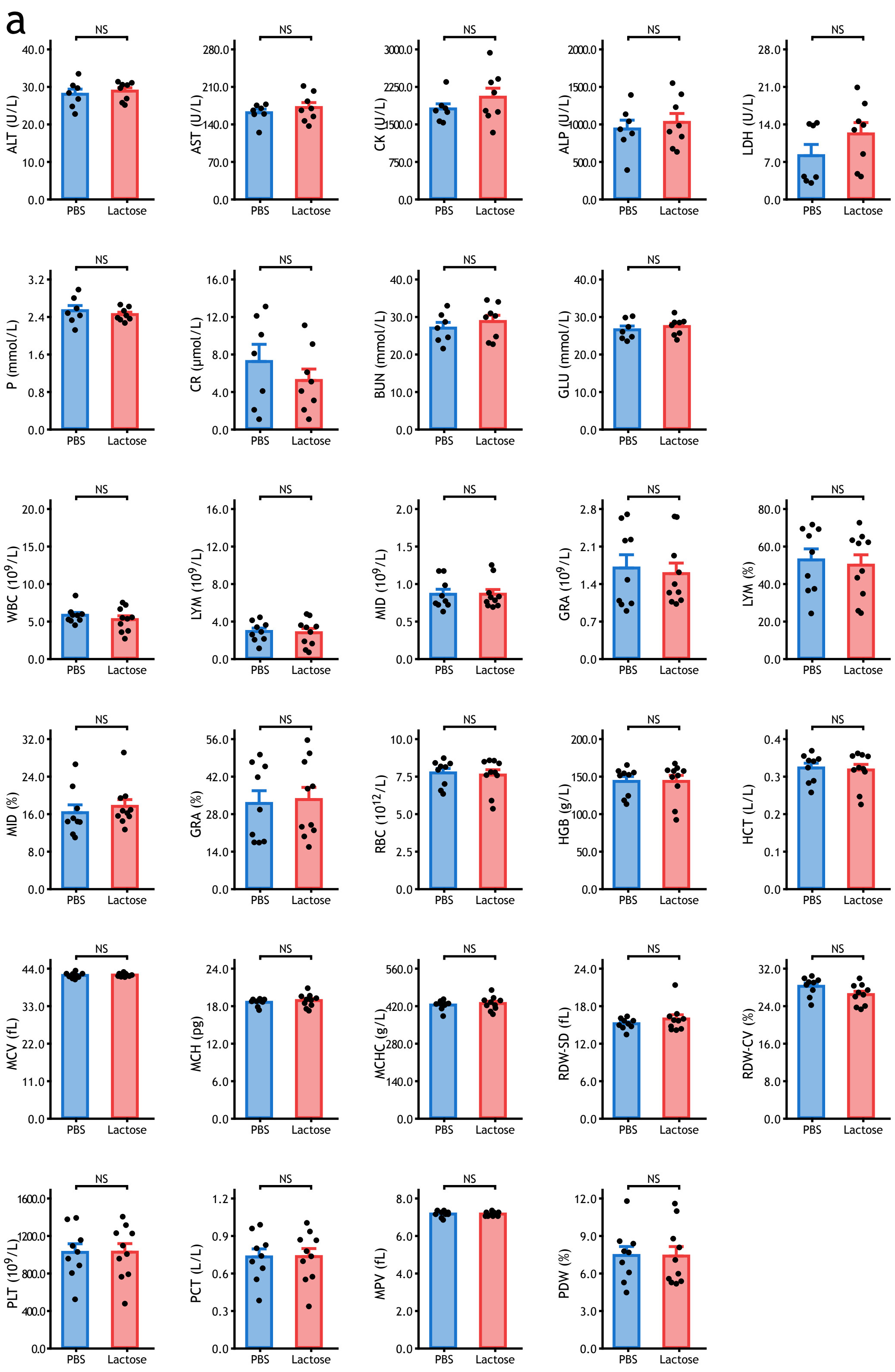
a
NS
PBS
Lactose
ALT (U/L)
AST (U/L)
CK (U/L)
ALP (U/L)
LDH (U/L)
P (mmol/L)
CR (μmol/L)
BUN (mmol/L)
GLU (mmol/L)
WBC (10^9/L)
LYM (10^9/L)
MID (10^9/L)
GRA (10^9/L)
LYM (%)
MID (%)
GRA (%)
RBC (10^12/L)
HGB (g/L)
HCT (L/L)
MCV (fL)
MCH (pg)
MCHC (g/L)
RDW-SD (fL)
RDW-CV (%)
PLT (10^9/L)
PCT (L/L)
MPV (fL)
PDW (%)

Figure S21

b

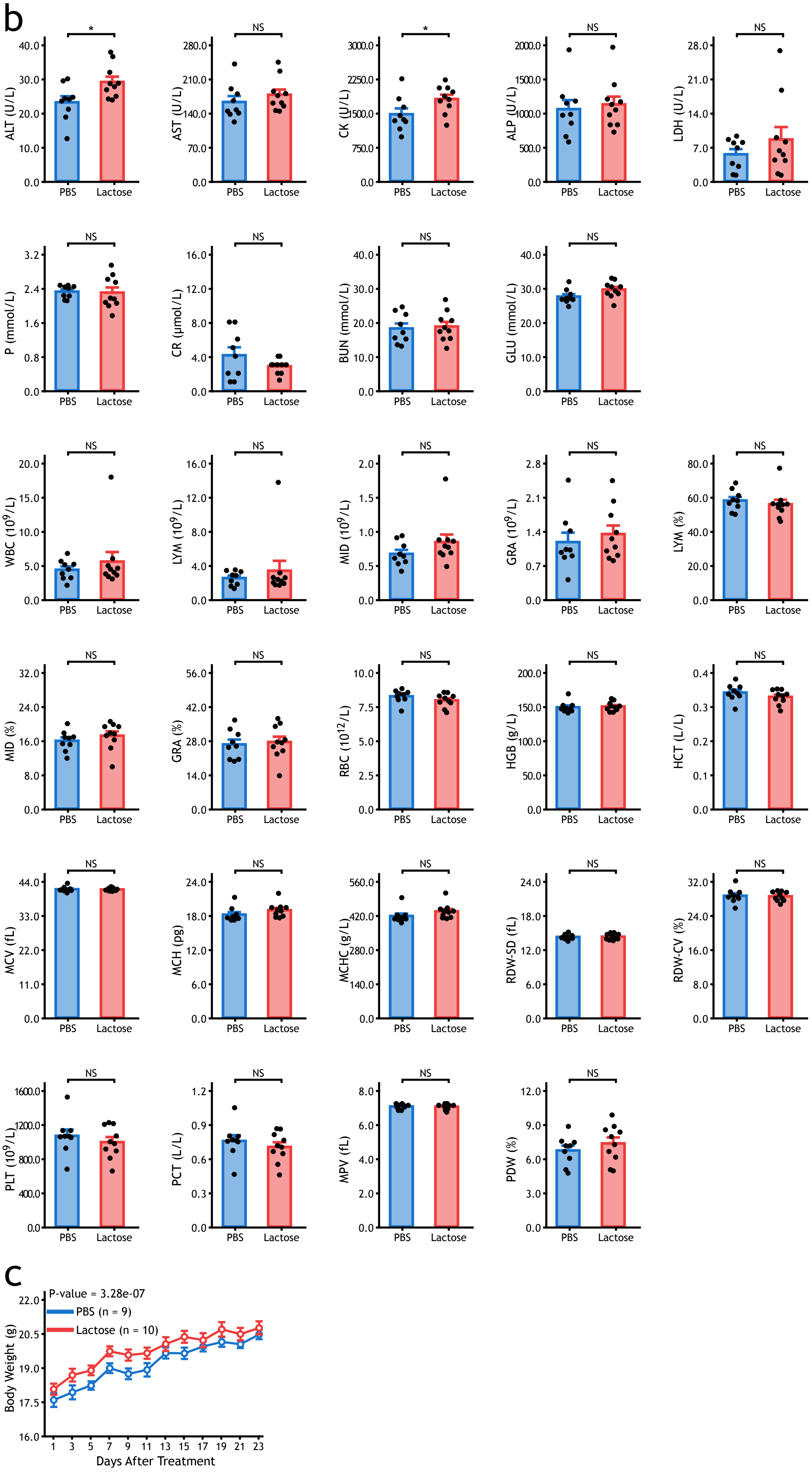

ALT (U/L)
AST (U/L)
CK (U/L)
ALP (U/L)
LDH (U/L)
P (mmol/L)
CR (μmol/L)
BUN (mmol/L)
GLU (mmol/L)
WBC (10^9/L)
LYM (10^9/L)
MID (10^9/L)
GRA (10^9/L)
LYM (%)
MID (%)
GRA (%)
RBC (10^12/L)
HGB (g/L)
HCT (L/L)
MCV (fL)
MCH (pg)
MCHC (g/L)
RDW-SD (fL)
RDW-CV (%)
PLT (10^9/L)
PCT (L/L)
MPV (fL)
PDW (%)
PBS
Lactose
*
NS
c
P-value = 3.28e-07
PBS (n = 9)
Lactose (n = 10)
Body Weight (g)
Days After Treatment


Figure S21

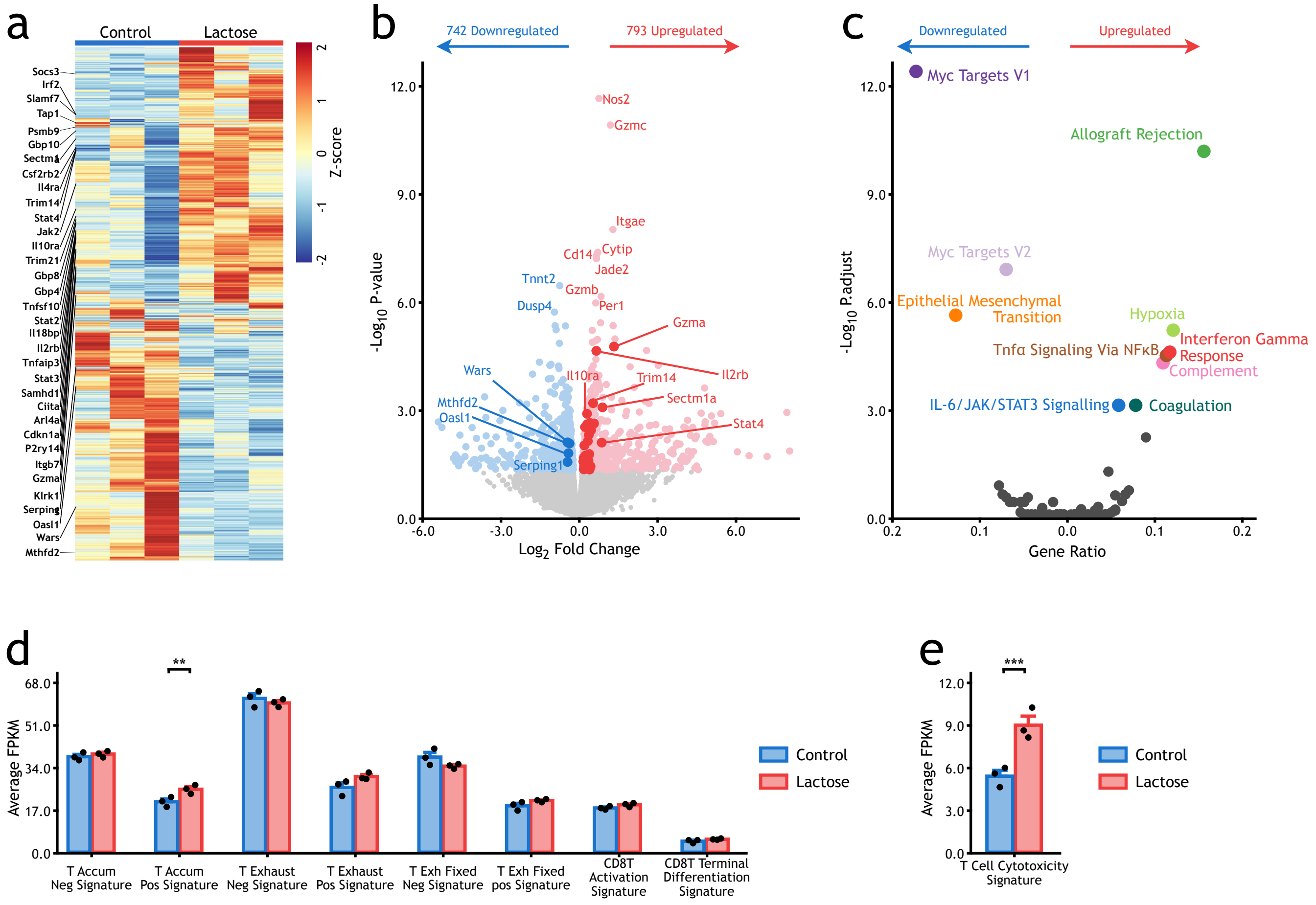

a
Control
Lactose
Z-score
Socs3
Irf2
Slamf7
Tap1
Psmb9
Gbp10
Sectm1a
Csf2rb2
Il4ra
Trim14
Stat4
Jak2
Il10ra
Trim21
Gbp8
Gbp4
Tnfsf10
Stat2
Il18bp
Il2rb
Tnfaip3
Stat3
Samhd1
Ciita
Arl4a
Cdkn1a
P2ry14
Itgb7
Gzma
Klrk1
Serping1
Oasl1
Wars
Mthfd2
b
742 Downregulated
793 Upregulated
-Log10 P-value
Log2 Fold Change
Nos2
Gzmc
Itgae
Cd14
Cytip
Jade2
Tnnt2
Gzmb
Dusp4
Per1
Gzma
Il2rb
Wars
Il10ra
Trim14
Mthfd2
Oasl1
Sectm1a
Stat4
Serping1
c
Downregulated
Upregulated
-Log10 P.adjust
Gene Ratio
Myc Targets V1
Allograft Rejection
Myc Targets V2
Epithelial Mesenchymal Transition
Hypoxia
Interferon Gamma Response
Tnfα Signaling Via NFκB
Complement
IL-6/JAK/STAT3 Signalling
Coagulation
d
Average FPKM
Control
Lactose
T Accum Neg Signature
T Accum Pos Signature
T Exhaust Neg Signature
T Exhaust Pos Signature
T Exh Fixed Neg Signature
T Exh Fixed pos Signature
CD8T Activation Signature
CD8T Terminal Differentiation Signature
e
Average FPKM
T Cell Cytotoxicity Signature


### Figure S22

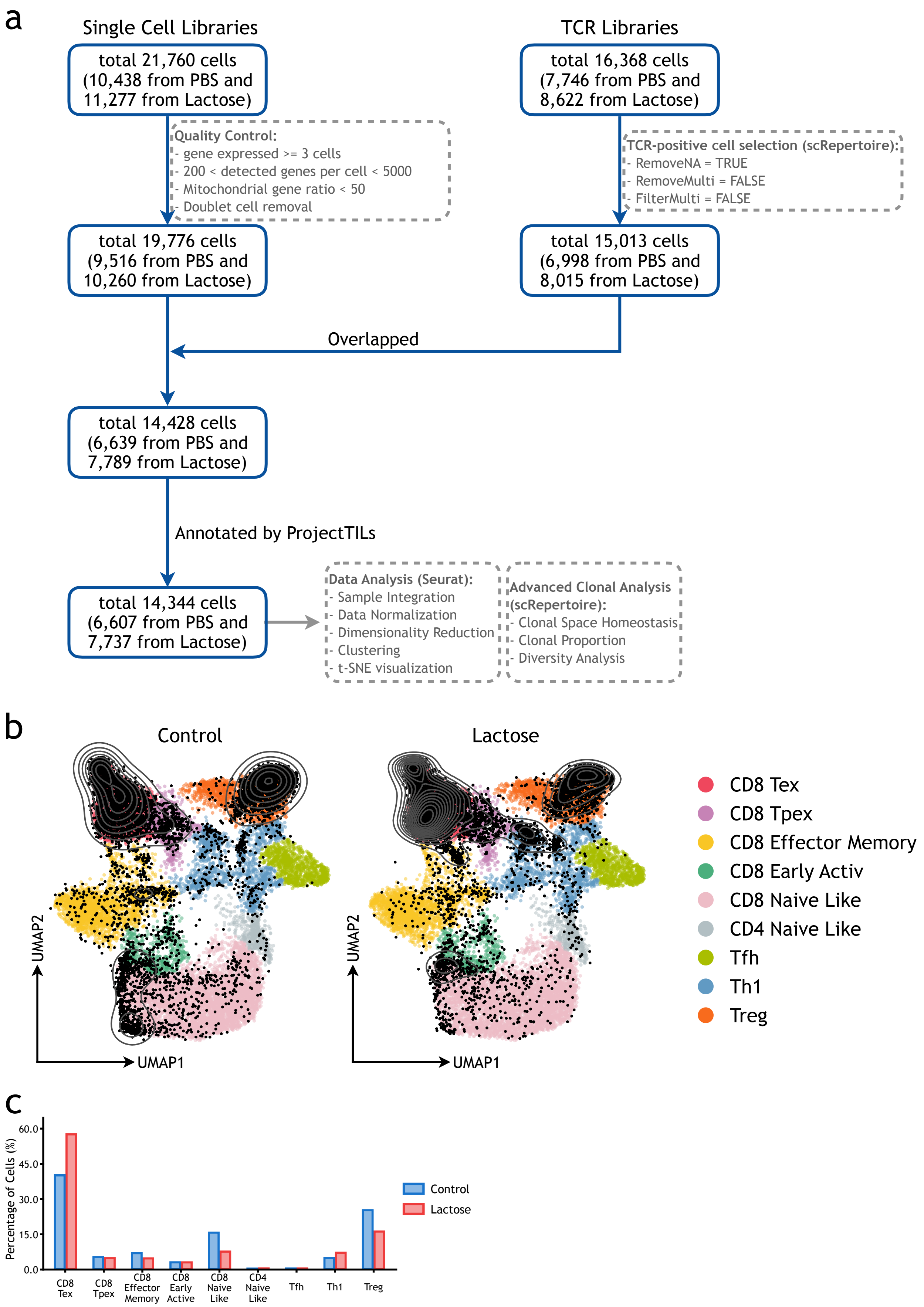
a
Single Cell Libraries
total 21,760 cells (10,438 from PBS and 11,277 from Lactose)
Quality Control:
- gene expressed >= 3 cells
- 200 < detected genes per cell < 5000
- Mitochondrial gene ratio < 50
- Doublet cell removal
total 19,776 cells (9,516 from PBS and 10,260 from Lactose)
TCR Libraries
total 16,368 cells (7,746 from PBS and 8,622 from Lactose)
TCR-positive cell selection (scRepertoire):
- RemoveNA = TRUE
- RemoveMulti = FALSE
- FilterMulti = FALSE
total 15,013 cells (6,998 from PBS and 8,015 from Lactose)
Overlapped
total 14,428 cells (6,639 from PBS and 7,789 from Lactose)
Annotated by ProjectTILs
total 14,344 cells (6,607 from PBS and 7,737 from Lactose)
Data Analysis (Seurat):
- Sample Integration
- Data Normalization
- Dimensionality Reduction
- Clustering
- t-SNE visualization
Advanced Clonal Analysis (scRepertoire):
- Clonal Space Homeostasis
- Clonal Proportion
- Diversity Analysis
b
Control
Lactose
UMAP2
UMAP1
CD8 Tex
CD8 Tpex
CD8 Effector Memory
CD8 Early Activ
CD8 Naive Like
CD4 Naive Like
Tfh
Th1
Treg
c
Percentage of Cells (%)
60.0
45.0
30.0
15.0
0.0
CD8 Tex
CD8 Tpex
CD8 Effector Memory
CD8 Early Active
CD8 Naive Like
CD4 Naive Like
Tfh
Th1
Treg
Control
Lactose

Figure S23

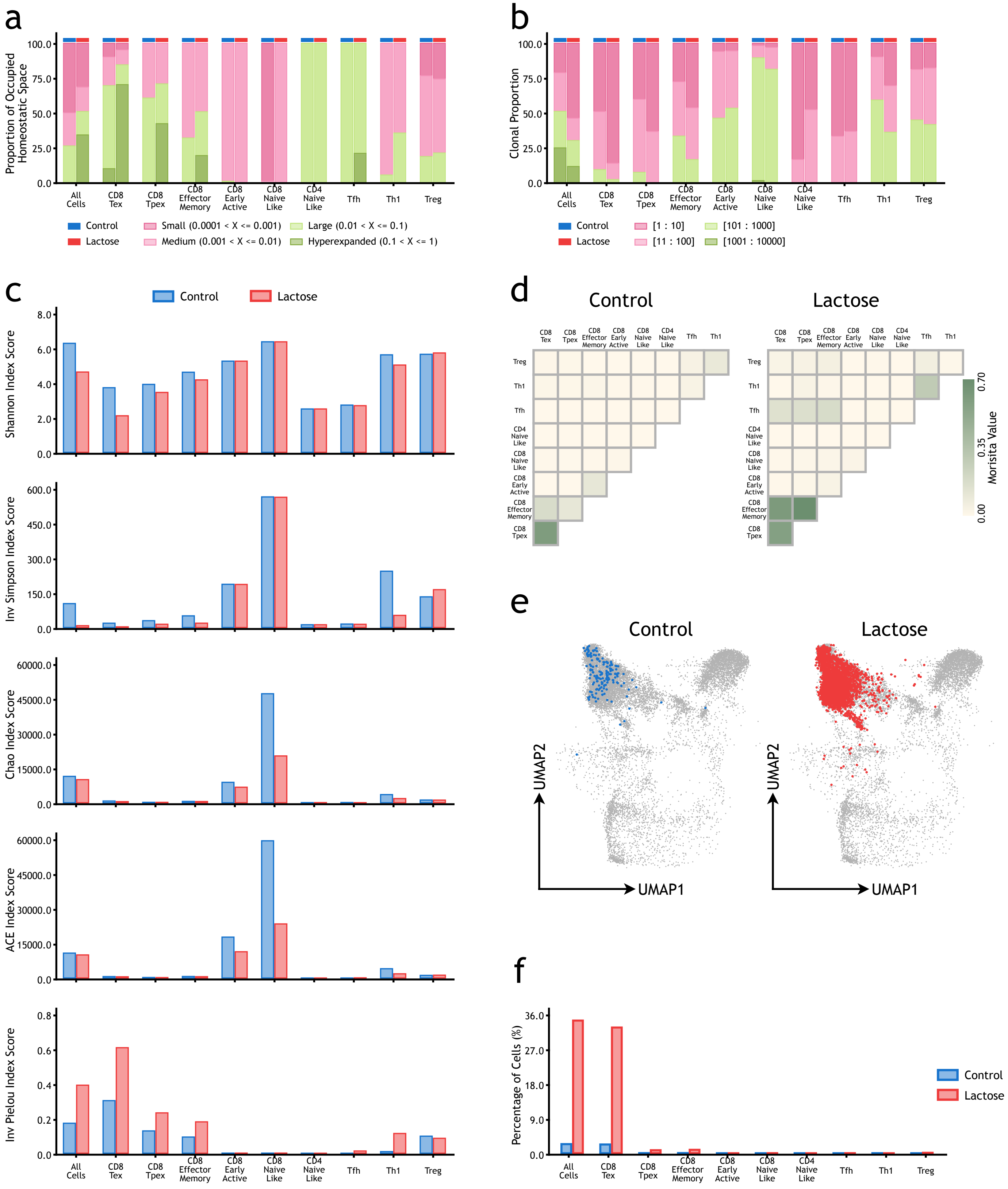
a
Proportion of Occupied Homeostatic Space
100.0
75.0
50.0
25.0
0.0
All Cells
CD8 Tex
CD8 Tpex
CD8 Effector Memory
CD8 Early Active
CD8 Naive Like
CD4 Naive Like
Tfh
Th1
Treg
Control
Lactose
Small (0.0001 < X <= 0.001)
Medium (0.001 < X <= 0.01)
Large (0.01 < X <= 0.1)
Hyperexpanded (0.1 < X <= 1)
b
Clonal Proportion
[1 : 10]
[11 : 100]
[101 : 1000]
[1001 : 10000]
c
Shannon Index Score
Inv Simpson Index Score
Chao Index Score
ACE Index Score
Inv Pielou Index Score
d
Control
Lactose
Morisita Value
0.70
0.35
0.00
e
Control
Lactose
UMAP1
UMAP2
f
Percentage of Cells (%)
36.0
27.0
18.0
9.0
0.0

### Figure S24

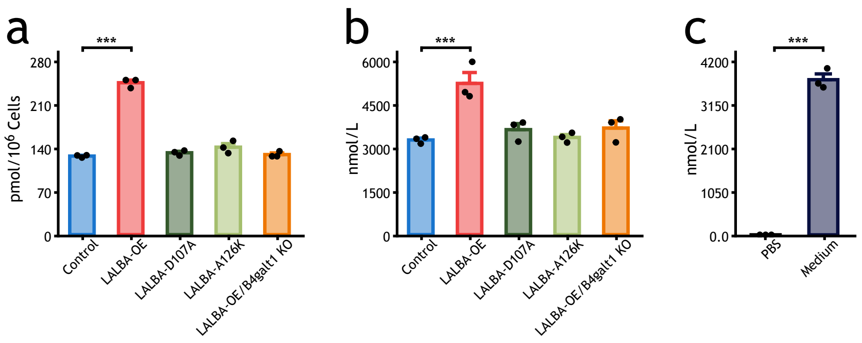
a
b
c
***
pmol/10^6 Cells
280
210
140
70
0
Control
LALBA-OE
LALBA-D107A
LALBA-A126K
LALBA-OE/B4galt1 KO
nmol/L
6000
4500
3000
1500
0
4200
3150
2100
1050
0.0
PBS
Medium

### Figure S25

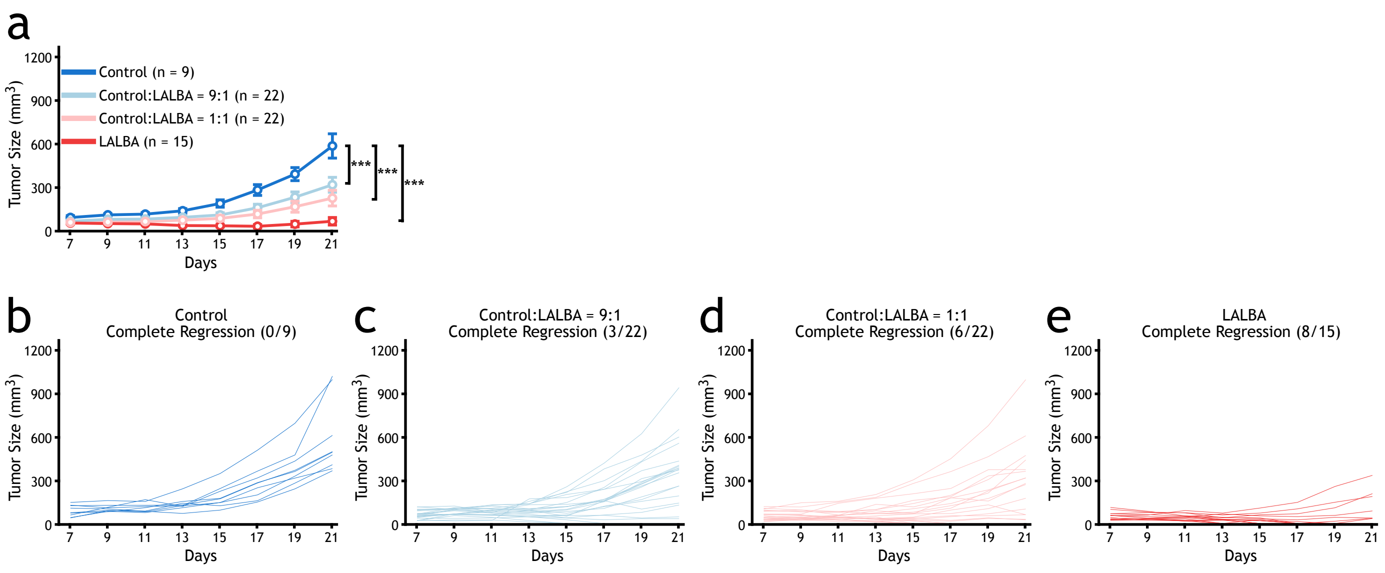

a
Control (n = 9)
Control:LALBA = 9:1 (n = 22)
Control:LALBA = 1:1 (n = 22)
LALBA (n = 15)
Tumor Size (mm$^3$)
Days
***
***
***
b
Control
Complete Regression (0/9)
c
Control:LALBA = 9:1
Complete Regression (3/22)
d
Control:LALBA = 1:1
Complete Regression (6/22)
e
LALBA
Complete Regression (8/15)


### Figure S26

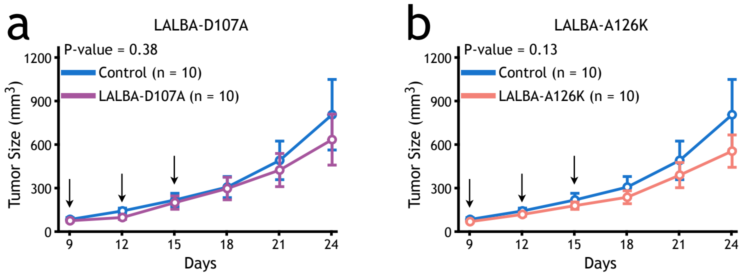
a
LALBA-D107A
P-value = 0.38
Control (n = 10)
LALBA-D107A (n = 10)
Tumor Size (mm3)
1200
900
600
300
0
9
12
15
18
21
24
Days
b
LALBA-A126K
P-value = 0.13
Control (n = 10)
LALBA-A126K (n = 10)
Tumor Size (mm3)
1200
900
600
300
0
9
12
15
18
21
24
Days

### Figure S27

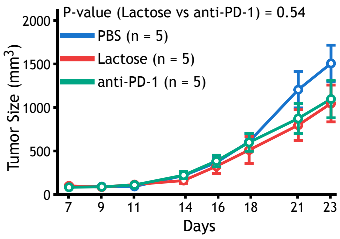
P-value (Lactose vs anti-PD-1) = 0.54
PBS (n = 5)
Lactose (n = 5)
anti-PD-1 (n = 5)
Tumor Size (mm$^3$)
2000
1500
1000
500
0
7
9
11
14
16
18
21
23
Days

### Figure S28

a

| Sample | Tumor type | Histology | Tumor site | Systematic treatment before surgery |
|---|---|---|---|---|
| Lung1 | Lung cancer | Poorly differentiated adenocarcinoma | Primary | No |
| Lung2 | Lung cancer | Poorly differentiated adenocarcinoma | Primary | No |
| Lung3.1 | Lung cancer | Poorly differentiated adenocarcinoma | Primary | No |
| Lung3.2 | Lung cancer | Poorly differentiated adenocarcinoma | Primary | No |
| Lung4 | Lung cancer | Moderately differentiated squamous cell carcinoma | Primary | No |
| CRC1 | Colon cancer | Moderately differentiated adenocarcinoma | Primary | No |
| CRC2 | Rectal cancer | Moderately differentiated adenocarcinoma | Primary | No |
| CRC3 | Rectal cancer | Moderately differentiated adenocarcinoma | Primary | No |
| CRC4 | Colon cancer | Moderately differentiated adenocarcinoma | Primary | No |
| CRC5 | Rectal cancer | Moderately differentiated adenocarcinoma, partial well differentiated adenocarcinoma | Primary | No |
| EC1 | Esophageal cancer | Squamous cell carcinoma | Primary | No |

b

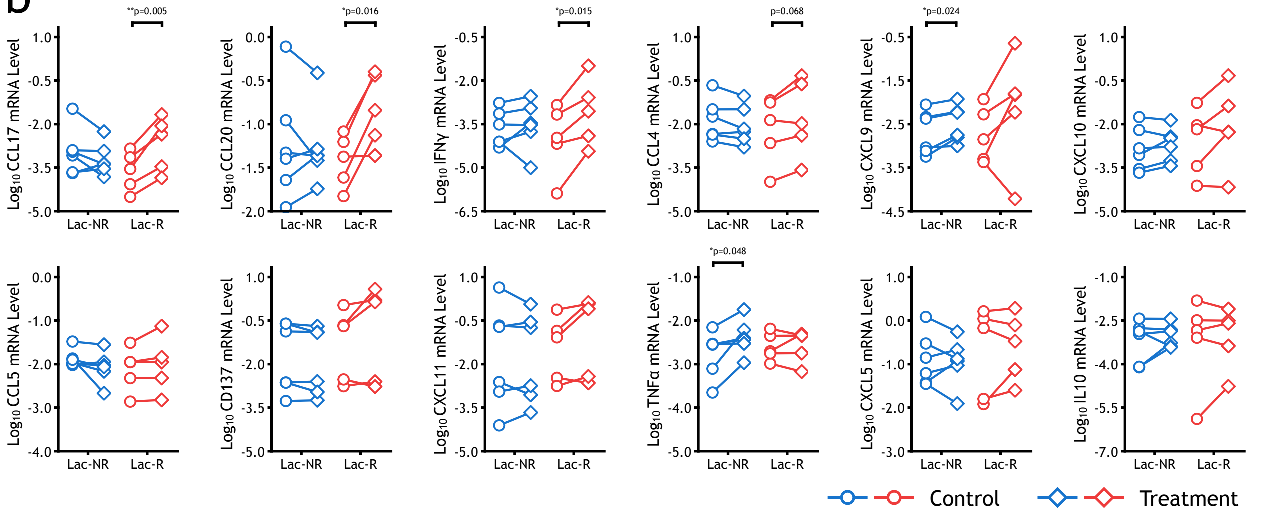


c

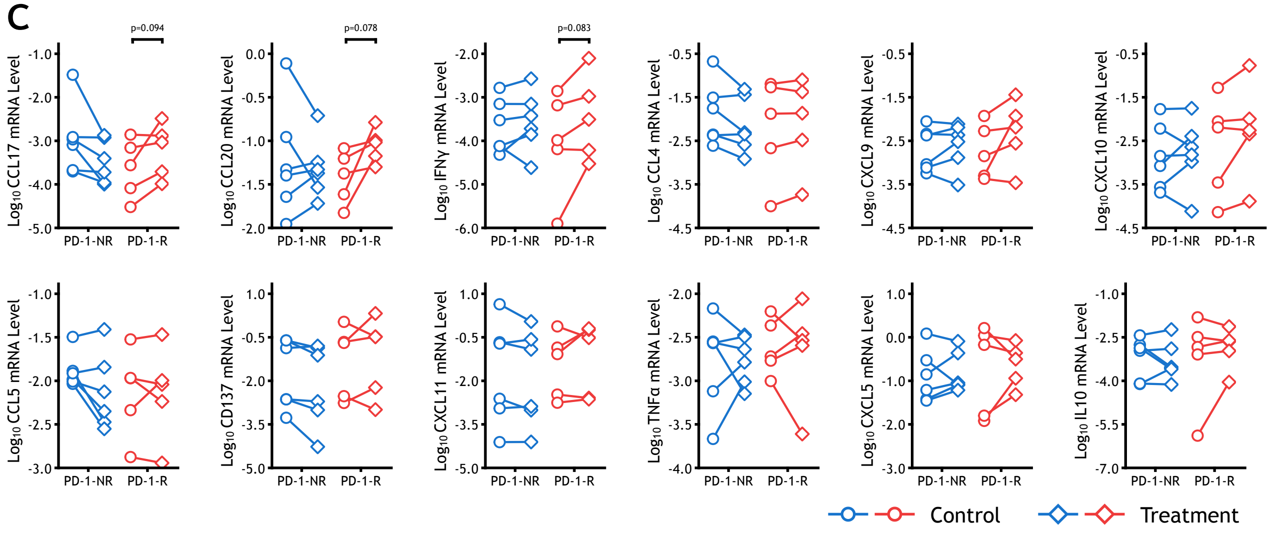


Figure S29
